# A lipid code-dependent phosphoswitch directs PIN-mediated auxin efflux in *Arabidopsis* development

**DOI:** 10.1101/755504

**Authors:** Shutang Tan, Xixi Zhang, Wei Kong, Xiao-Li Yang, Gergely Molnár, Jiří Friml, Hong-Wei Xue

## Abstract

Directional intercellular transport of the phytohormone auxin mediated by PIN FORMED (PIN) efflux carriers plays essential roles in both coordinating patterning processes and integrating multiple external cues by rapidly redirecting auxin fluxes. Multilevel regulations of PIN activity under internal and external cues are complicated; however, the underlying molecular mechanism remains elusive. Here we demonstrate that 3’-Phosphoinositide-Dependent Protein Kinase1 (PDK1), which is conserved in plants and mammals, functions as a molecular hub integrating the upstream lipid signalling and the downstream substrate activity through phosphorylation. Genetic analysis uncovers that loss-of-function *Arabidopsis* mutant *pdk1.1 pdk1.2* exhibits a plethora of abnormalities in organogenesis and growth, due to the defective PIN-dependent auxin transport. Further cellular and biochemical analyses reveal that PDK1 phosphorylates D6 Protein Kinase to facilitate its activity towards PIN proteins. Our studies establish a lipid-dependent phosphorylation cascade connecting membrane composition-based cellular signalling with plant growth and patterning by regulating morphogenetic auxin fluxes.

## Introduction

Due to a sessile lifestyle, plants have evolved strict developmental programming as well as adaptive plasticity in response to diverse environmental stimuli, which is ensured by a framework of multiple signalling pathways. The phytohormone auxin (indole-3-acetic acid, IAA) plays a pivotal role in shaping plant growth and development, in a concentration-dependent manner. The establishment and maintenance of local auxin minima, maxima and gradients rely on directional intercellular transport, which is mediated by a subset of auxin influx carriers AUXIN1 (AUX1)/LIKE AUX1s (LAXs)^1^, PIN (PIN-FORMED) efflux carriers^2, 3^ and ABCB-type transporters^4^. It has been well documented that the activity and polar localization of PIN proteins are regulated by intracellular trafficking as well as post-translational modifications, including reversible phosphorylation^5^.

PINs are phosphorylated at multiple serine/threonine residues by various protein kinases, including PINOID (PID)/WAVY ROOT GROWTHs (WAGs)^6–8^, D6 PROTEIN KINASEs (D6PK/D6PKL1∼3, hereafter as D6PKs)^9, 10^, PROTEIN KINASE ASSOCIATED WITH BRX (PAX)^11^, MITOGEN PROTEIN KINASE3/6 (MPK3/6)^12^ and CDPK-RELATED KINASE5 (CRK5)^13^, and dephosphorylated by Protein Phosphatase 2A (PP2A)^8, 14^, Protein Phosphatase 6 (PP6)^15^ and PP1^16^. PID-dependent phosphorylation and PP2A-mediated dephosphorylation antagonize with each other^6, 14^, acting as a binary switch determining the apical-basal targeting of PIN proteins. More PID kinase activity results in more apical localization of PINs; conversely, less PIN phosphorylation by PID gives rise to the reduced apical (or more basal) localization. In contrast to PID, D6PKs and PAX phosphorylate basal-localized PINs to regulate their activity^17^. To date, little is known about the upstream regulatory mechanism by which this phosphorylation increases or declines, except for that both PID^18, 19^ and D6PK^20^ have been reported to bind phospholipids, which might determine their subcellular localizations.

PDK1 acts as a master regulator of the AGC [named after the cAMP dependent (PKA) and cGMP-dependent protein kinases (PKG), and protein kinase C (PKC)] kinases in eukaryotes^21^. In animals, PDK1 is activated by PI(3,4,5)P_3_ and is critical for growth and development, as well as multiple metabolic and signalling pathways. The PDK1 loss-of-function mutants of fruitflies^22^ and mice^23^ exhibited a smaller size compared to that of wild type. By contrast, PI(3,4,5)P_3_ is undetectable and does not bind to the PDK1 orthologue in plants. *Arabidopsis* PDK1 was reported to be activated by phosphatidic acid (PA), as well as a wide spectra of lipids^24^. To date, several of the 39 AGC kinases in *Arabidopsis* have been functionally characterized as putative downstream substrates of PDK1, including PID and AGC2-1 (also called OXI1, OXIDATIVE SIGNAL INDUCIBLE1)^25–29^. As mentioned above, PID phosphorylates PIN auxin efflux carriers to regulate their polar localization as well as activity, and thus the directionality of intercellular auxin transport^6, 10, 30^. AGC2-1 regulates root hair growth and development^25, 28^, as well as plants’ response to the beneficial endophytic fungus *Piriformospora indica*^27^. However, due to a lack of well characterized knockout lines, the exact roles of PDK1 during development as well as its functional substrates remain poorly understood.

Here we demonstrate the biological function of *Arabidopsis* PDK1 as well as its regulation by phospholipids. Our comprehensive biochemical and cell biology analyses illustrate an important role of PDK1 in regulating PIN-dependent polar auxin transport through phosphorylating the D6PK subclade of AGC kinases, and thus the accompanied organogenesis processes and tropic responses.

## Results

### Loss-of-function mutants of *Arabidopsis PDK1.1* and *PDK1.2* exhibit auxin related defects

In *Arabidopsis* genome, there are two homologous genes encoding PDK1 proteins, *PDK1.1* and *PDK1.2* (*AT5G04510* and *AT3G10540*, also known as *PDK1-1* and *PDK1-2*^28^). These two paralogues exhibit 91.06% identity in amino acid sequences (Supplementary Fig. 1a) and conserved structures, i.e. a kinase domain (PDK1N) and a Pleckstrin Homology (PH) domain (PDK1C) at N- and C-terminus respectively (Supplementary Fig. 1b). Analysis of promoter::β-glucuronidase (GUS) lines revealed a broad and similar expression pattern, implying functional redundancy for *PDK1.1* and *PDK1.2.* Both *PDK1s* showed predominant transcription in vascular tissues of roots and cotyledons, columella cells, lateral root primordia, as well as dark-grown etiolated seedlings (Supplementary Fig. 2). Interestingly, *PDK1.1* was expressed in root columella cells, but *PDK1.2* not (Supplementary Fig. 2c, h), suggesting possible functional divergence.

To explore the biological function of PDK1s, two knock-out T-DNA insertional mutants, *pdk1.1-2* (hereafter as *pdk1.1*) and *pdk1.2-4* (*pdk1.2*) (Supplementary Fig. 3a), were verified and used for further analysis (Supplementary Fig. 3b, c). Neither *pdk1.1* nor *pdk1.2* single mutant exhibited any obvious phenotype compared to wild type (WT, Col-0) under normal conditions, whereas the *pdk1.1 pdk1.2* double mutant displayed severe defects in various growth and developmental processes, including smaller rosettes, shorter siliques, and lower fertility than those of Col-0 (Fig. 1a, Supplementary Fig. 4). Notably, *pdk1.1 pdk1.2* exhibited a dramatic decrease in lateral root number compared with that of Col-0 at the seedling stage (Fig. 1b, c). By transforming a *pPDK1.1::PDK1.1* construct into *pdk1.1 pdk1.2* (Supplementary Fig. 4a), growth defects of various tissues at different stages were completely rescued (Supplementary Fig. 4b-e), confirming the causative role of *PDK1* deficiency in the *pdk1.1 pdk1.2* double mutant.

**Figure 1.**
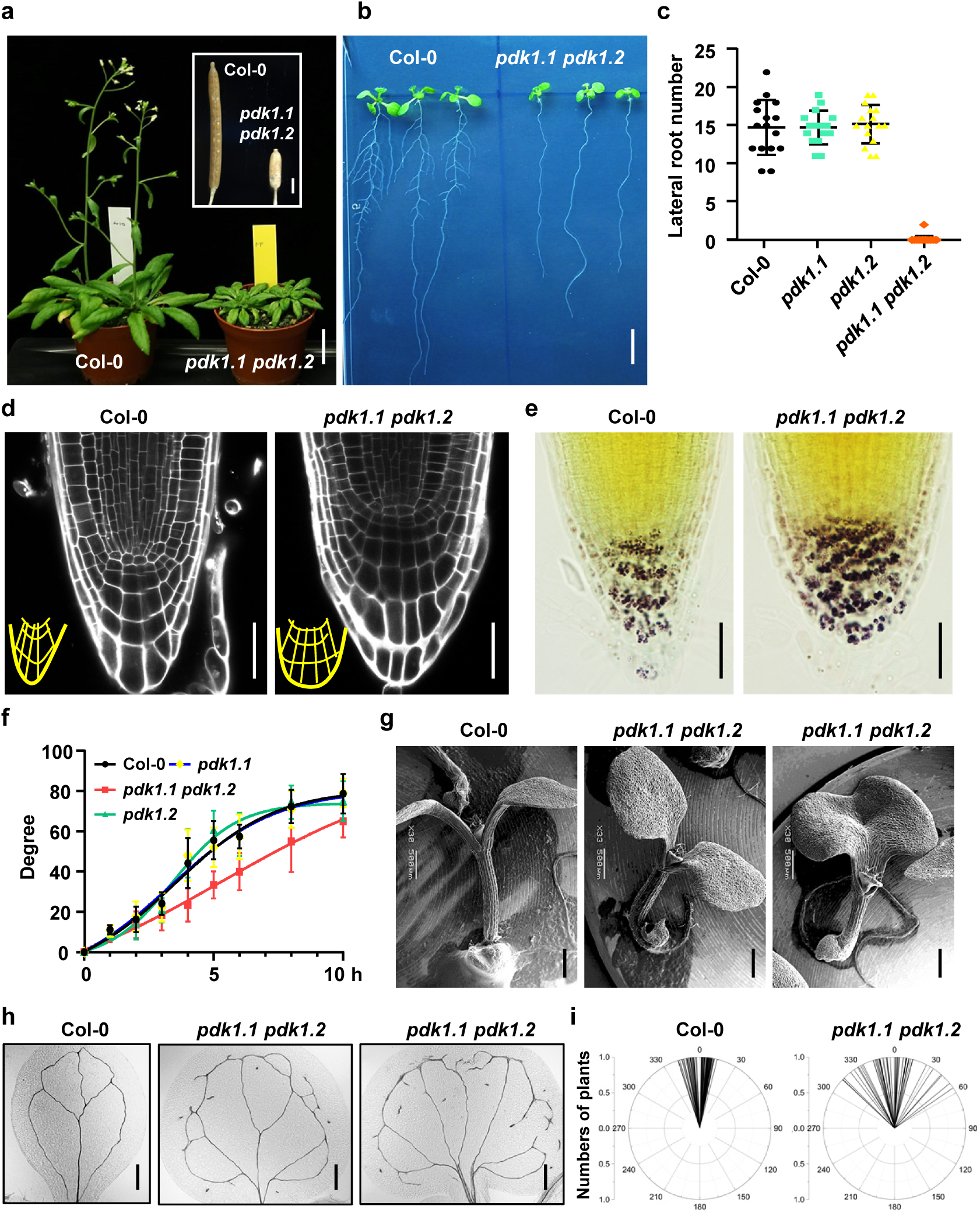
Loss of function of *PDK1.1* and *PDK1.2* attenuates *Arabidopsis* growth and development. a. Observations of 35-day-old adult plants of Col-0 and *pdk1.1 pdk1.2* indicated that *PDK1.1* and *PDK1.2* were required for the normal growth and development of *Arabidopsis*. Scale bar, 2 cm. The internal panel showed that *pdk1.1 pdk1.2* produced shorter siliques and fewer seeds (from 50-day-old plants; scale bar, 1 mm). b. Deficiency of *PDK1.1* and *PDK1.2* impaired lateral root formation. A representative photo of 12-day-old Col-0 and *pdk1.1 pdk1.2* seedlings was shown. Scale bar, 1 cm. c. Reduced lateral root numbers of *pdk1.1 pdk1.2*. Lateral root number of plate-grown 12-day-old Col-0 and *pdk1.1 pdk1.2* seedlings was counted and shown as mean ± standard deviation (s.d., n = 15∼20 per genotype). d. PI staining indicated an overgrowth of root columella cells in *pdk1.1 pdk1.2*. Roots of 7-day-old Col-0 and *pdk1.1 pdk1.2* seedlings were stained by PI and imaged by CLSM. Representative images were shown. Scale bars, 20 µm. e. Lugol staining of 7-day-old seedlings indicates overgrowth of root columella cells and over-accumulation of starch granules in *pdk1.1 pdk1.2*. Representative photos are shown. Scale bars, 20 µm. f. *pdk1.1 pdk1.2* seedlings exhibited a reduced root gravitropic response. Five-day-old Col-0, *pdk1.1*, *pdk1.2*, and *pdk1.1 pdk1.2* seedlings were gravistimulated by 90° reorientation, and root inclination at each hour were measured and shown as mean ± s.d. (n = 10∼24). g. Scanning electron microscopy (SEM) observation showed fused cotyledons of a subset of 7-day-old *pdk1.1 pdk1.2* seedlings (right). Scale bars, 500 µm. h. Differential interference contrast (DIC) microscopy images of the vasculature pattern of Col-0 and *pdk1.1 pdk1.2*. Representative images of 7-day-old seedlings were shown. Scale bars, 1 mm. i. Deficiency of *PDK1.1* and *PDK1.2* led to less gravitropic hypocotyls under dark. Etiolated seedlings of Col-0 (n=51) and *pdk1.1 pdk1.2* (n=68) were grown under dark for 90 hours and hypocotyl angles were measured by Image J.

Microscopic analysis revealed that *pdk1.1 pdk1.2* exhibited a broader region of columella cells (Fig. 1d), which are filled with starch granules (statoliths) responsible for gravity perception^31, 32^. Lugol staining confirmed the overgrowth of columella cells, suggesting a potential role of PDK1 in gravitropism (Fig. 1e). Consistently, the root of *pdk1.1 pdk1.2* responded to gravistimulation more slowly than that of Col-0 by 90° reorientation (Fig. 1f). Notably, *pdk1.1 pdk1.2* exhibited a more pronounced agravitropic root phenotype under dark (Supplementary Fig. 5a, b). In addition, ∼18% of *pdk1.1 pdk1.2* seedlings exhibited fused cotyledons, which, however, were barely found in Col-0 or *pdk1* single mutants (Fig. 1g, and Table S1), indicating an essential role of PDK1s in boundary formation during organogenesis.

These above morphological defects of *pdk1.1 pdk1.2* are reminiscent of auxin synthesis, transport, or signalling deficiency as reported previously^3^. Therefore, we investigated more auxin related phenotypic outputs. Auxin is required for vasculature patterning^3, 33^. In comparison to the four-loop vasculature of Col-0, *pdk1.1 pdk1.2* showed a fragmented pattern in cotyledons (Fig. 1h). In addition, *pdk1.1 pdk1.2* hypocotyls were unable to grow upright in darkness (Fig. 1i, Supplementary Fig. 5a, c), and *pdk1.1 pdk1.2* bent slowly toward a directional light source (Supplementary Fig. 5d, e), indicating defects in both gravitropism and phototropism. Plants form an apical hook to protect the shoot apical meristem when protruding out of the earth during germination ^34^, as observed for Col-0. However, *pdk1.1 pdk1.2* failed to form a complete apical hook though it grew a comparable length of the etiolated hypocotyls (Supplementary Fig. 6a-c), even upon the treatment with the ethylene precursor 1-aminocyclopropane-1-carboxylic acid (ACC), an early initiation signal which leads to an exaggerated apical hook through crosstalk with the auxin pathway^34, 35^ (Supplementary Fig. 6d). Likewise, hypocotyl elongation in response to high temperature (29°C)^36, 37^ was suppressed in *pdk1.1 pdk1.2* as expected (Supplementary Fig. 6e-f). In summary, *PDK1* deficiency leads to a plethora of defects in auxin-regulated organogenesis processes and tropic responses, resembling those mutants defective in the biosynthesis, transport or signalling of auxin^3, 38, 39^.

### Deficiency of *PDK1.1* and *PDK1.2* affects the local auxin distribution

Further, we generated a cross between *pdk1.1 pdk1.2* and an auxin responsive reporter *DR5rev::GFP*^40^ to monitor the distribution or the response of auxin. In Col-0, *DR5rev::GFP* was expressed in vascular tissues, with maxima in root columella cells and leaf tips. Being consistent with the overgrowth of columella cells, there was a broader expression of *DR5rev::GFP* in *pdk1.1 pdk1.2* than that of Col-0 (Fig. 2a, b). Under gravistimulation by 90° reorientation, there was a polar auxin flow from the root tip to the lower side of the lateral root cap^41, 42^, revealed by a relocated expression pattern of *DR5rev::GFP* (Fig. 2c). However, this was undetectable in *pdk1.1 pdk1.2* (Fig. 2d, e), indicating a deficiency of basipetal auxin transport activity in root tips of *pdk1.1 pdk1.2*. In line with the dramatic morphological changes, two auxin maxima sites for the fused cotyledon were observed (Supplementary Fig. 7a, e), while none for the lateral root primordia was found in *pdk1.1 pdk1.2* (Supplementary Fig. 7b, f).

**Figure 2.**
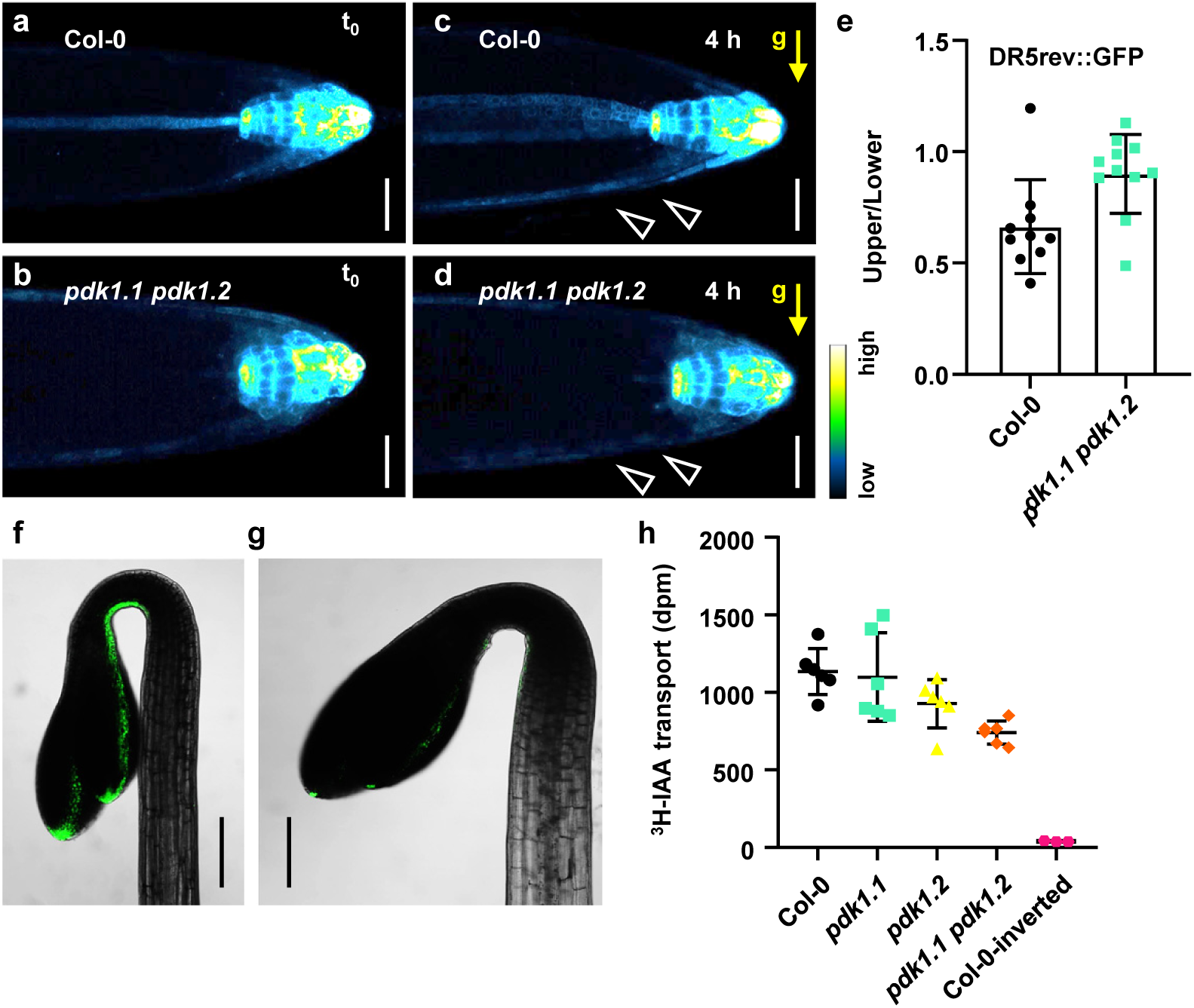
PDK1.1 and PDK1.2 regulate auxin distribution through PIN-mediated auxin transport. a-d. *pdk1.1 pdk1.2* showed defects in gravistimulation-induced DR5rev::GFP relocation. Four-day-old seedlings of *DR5rev::GFP* and *pdk1.1 pdk1.2 DR5rev::GFP* were gravistimulated by 90° reorientation for 4 h, then imaged by CLSM. The “Green Fire Blue” LUT was used for visualization based on fluorescence intensity by the FIJI program. Open arrowheads indicated the location of DR5rev::GFP at the lower side. Scale bars, 20 µm. e. *pdk1.1 pdk1.2* was defective in gravistimulation-induced DR5rev::GFP relocation, in comparison to Col-0. The ratio of GFP fluorescence between the upper side and the lower side was measured (n = ∼10). f-g. Observation of the auxin responsive reporter *DR5rev::GFP* indicated a decrease of the auxin maxima in shoots of *pdk1.1 pdk1.2*. Dark-grown 4-day-old *DR5rev::GFP* (f) and *pdk1.1 pdk1.2 DR5rev::GFP* (g) etiolated seedlings were imaged by CLSM. Scale bars, 200 µm. h. Deficiency of *PDK1.1* and *PDK1.2* resulted in decreased basipetal auxin transport. Decapitated hypocotyls of 7-day-old etiolated seedlings were used to measure the transport of [^3^H]-IAA, and 15 individuals were pooled as a biological replicate (n=6). Control: inverted hypocotyls of Col-0, with the [^3^H]-IAA droplets placed at the rootward end.

There was a dramatic depletion of the DR5rev::GFP reporter signal at the concave side of the apical hook of *pdk1.1 pdk1.2* etiolated seedlings under dark (Fig. 2f, g). Similarly, ACC treatment induced increase of *DR5rev::GFP* in the apical hook accompanied with the exaggerated curvature (Supplementary Fig. 7c) or roots under dark (Supplementary Fig. 7d) was undetectable (Supplementary Fig. 7g) or dramatically decreased (Supplementary Fig. 7h) in *pdk1.1 pdk1.2*. Collectively, *PDK1* deficiency abolished the plants’ ability of maintaining a functional local auxin concentration during various organogenesis processes, as well as that of redirecting the auxin flow in tropic responses.

Based on the scenario that PDK1 acts as a master regulator of AGC kinases in different organisms^21^, and the crucial roles of AGC kinases in regulating auxin transport^30, 43^, we speculate that PDK1 might function in auxin pathway through regulating specific AGC kinases. Of the 39 AGC members, PID/WAGs^6–8, 44^, D6PKs^9^ and PAX^11^ have been reported to regulate either the polarity (by PID/WAGs) or the activity (by all of these) of PIN proteins respectively through phosphorylation, and hence modulate auxin fluxes. Therefore, we firstly measured the basipetal auxin transport in intact etiolated hypocotyls with [^3^H]-IAA. Results showed that there was indeed a decrease in IAA transport in *pdk1.2* and *pdk1.1 pdk1.2*, but not in *pdk1.1*, compared to that of Col-0 (Fig. 2h).

These observations suggest that PDK1s are required for polar auxin transport, and the subsequent asymmetric auxin distribution, in which PIN auxin exporters play a critical role^3, 5^. Thus, we examined the polarity of PIN proteins firstly. Immunofluorescence with anti-PIN1 or anti-PIN2 antibodies indicated that there was no polarity change for PIN1 (Supplementary Fig. 8a) or PIN2 (Supplementary Fig. 8b) in *pdk1.1 pdk1.2*. Moreover, by crossing with fluorescent protein-fused PIN marker lines, it was found that though with an expanded region for *PIN3-GFP* expression in root columella cells, there was no change (i.e., no switch or decrease) in the polar localization of PIN1-YFP (Supplementary Fig. 8c), PIN2-GFP (Supplementary Fig. 8d), and PIN3-GFP (Supplementary Fig. 8e-h) respectively. This indicates that PDK1s do not play a major role in regulating PIN polarity. We speculate that the change of *PIN3-GFP* expression pattern might be result of, but not the driving force for, the corresponding morphological change in root tips.

Further immunofluorescence using an inducible overexpression line (*XVE>>Venus-PDK1.1*) revealed that *PDK1.1* overexpression did not affect the PIN1 or PIN2 polarity either (Supplementary Fig. 9a-d). Collectively, we conclude that PDK1s function in regulating polar auxin transport, through modulating the activity, but not the polarity, of PIN proteins.

### PDK1s bind to phospholipids and localizes to both cytoplasm and the basal side of plasma membrane (PM)

In animals, PDK1 is activated by PI(3,4,5)P_3_, which, however, seems not exist in plants^45^, suggesting a distinct regulatory mode for plant PDK1. *Arabidopsis* PDK1.1 was previously reported to bind to and being activated by phosphatidic acid (PA) *in vitro*^28^. Besides, PDK1.1 can also bind to a wide spectra of phospholipids^24^. To investigate which exact lipid species, PDK1 can bind and transduce the signal, recombinant proteins of the full length, N-termini, and C-termini of both PDK1.1 and PDK1.2 were expressed in *Escherichia coli* and purified proteins were used for biochemical analyses. Lipid-protein blot overlay assays using PIP strips with various phospholipids showed that, full length of PDK1.1 and PDK1.2 bound to an array of phospholipids, including PI monophosphates (PI3P, PI4P, PI5P), PI bisphosphates [PI(4,5)P_2_, PI(3,4)P_2_, PI(3,5)P_2_], and PA (Fig. 3a). The N-terminal kinase domain of PDK1.1 (PDK1.1N) showed a similar lipid binding spectrum, whereas PDK1.2N only bound to less lipid species, such as PI3P and PA (Fig. 3a and Supplementary Fig. 10). As expected, the C-termini His-PDK1C and His-PDK1.2C bound to the same lipid species as the full length did (Fig. 3a).

**Figure 3.**
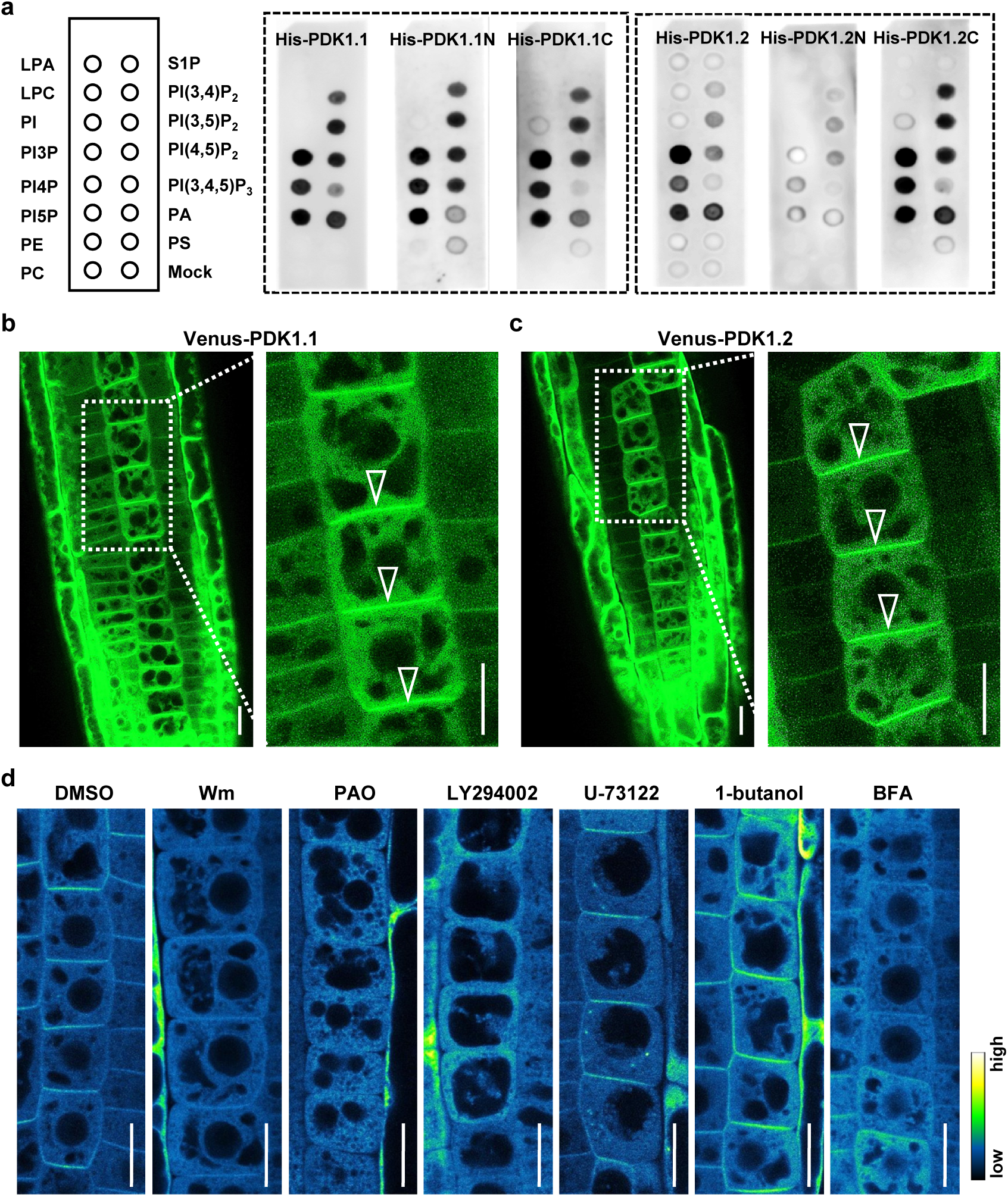
PDK1.1 and PDK1.2 bind to phospholipids and localize to both cytoplasm and the basal side of PM in root cells. a. Lipid-protein blot overlays revealed that recombinant His-PDK1.1 and His-PDK1.2 proteins directly bound to various phospholipids. PIP strips were incubated with His-tagged full-length PDK1.1/PDK1.2, the N-terminal kinase domain (PDK1.1/2N) or the C-terminal PH domain (PDK1.1/2C), respectively, and detected by an anti-His antibody. b-c. Venus fused PDK1.1 (b) and PDK1.2 (c) localized to both cytoplasm and the basal side of plasma membrane (PM). Open arrowheads indicate the basal polar localization. Scale bars, 10 µm. Four-day-old seedlings of *35S::Venus-PDK1.1* and *35S::Venus-PDK1.2* were imaged by CLSM. One representative of at least three independent lines was shown. d. Lipid biosynthesis inhibitors impaired the PM localization of Venus-PDK1.1. Four-day-old seedlings of *35S::Venus-PDK1.1* were treated with different chemicals for 30 min, and imaged by CLSM. Wortmannin (Wm, 33 µM, inhibitor of PI3K and PI4K), PAO (30 µM, an inhibitor of PI3K), LY294002 (50 µM, an inhibitor of PI3K), U73122 (5 µM, an inhibitor of PI-PLC), 1-butanol (0.8%, an inhibitor of PLD), and Brefeldin A (BFA, 50 µM, an inhibitor of ARF GEF) were applied. Scale bars, 10 µm.

To explore whether lipid binding PH domains determine the subcellular localizations, PDK1.1 and PDK1.2, together with the above truncations, with N-termini fused to mCherry or Venus driven by a 35S promoter were transformed into Col-0 and *pdk1.1 pdk1.2* respectively. Full length of PDK1.1 or PDK1.2 completely rescued the growth and developmental defects of *pdk1.1 pdk1.2* (Supplementary Fig. 11a-c), indicating that they were biologically functional. Observations of Venus-PDK1.1 and Venus-PDK1.2 (Fig. 3b, c) and mCherry-PDK1.1 and mCherry-PDK1.2 (Supplementary Fig. 11d-e) revealed their presence in the cytoplasm and at the basal side of PM.

Given the soluble properties of PDK1.1 and PDK1.2, the basal PM localizations might be determined by lipid binding preferences. Pharmacological treatments with various lipid biosynthesis inhibitors were performed to investigate the regulatory roles of different lipids in modulating the subcellular localization of PDK1s. The PI3K and PI4K inhibitors^18, 20^, wortmannin (Wm), phenylarsine oxide (PAO), and LY294002, completely abolished the PM localization of Venus-PDK1.1, however, the PI-PLC inhibitor U-73122 as well as the PA synthesis inhibitor 1-butanol, did not (Fig. 3d). These results indicate that PI3P and PI4P, but not PI(4,5)P_2_ or PA, majorly affect the polar PM targeting of PDK1s. Furthermore, an ARF GTPase inhibitor, brefeldin A (BFA, targeting ARF GEFs)^46^, also impaired the PM localization of Venus-PDK1.1, revealing the involvement of the intracellular trafficking machinery in the PM targeting of PDK1s. In addition, both Venus-PDK1.1/2N and Venus-PDK1.1/2C fusion proteins depleted from PM (Supplementary Fig. 11f-i), indicating that both the PH domain and the kinase domain are required for the PM localization of PDK1s. Additionally, analysis of the *pPDK1.1::Venus-PDK1.1* confirmed that Venus-PDK1.1 localized to both cytoplasm and PM. In line with the overexpression line, Venus-PDK1.1 exhibited a predominant localization at the basal side of PM in the stele (Supplementary Fig. 12).

### PDK1 regulates auxin transport through phosphorylating D6PKs

Plant AGC kinases share a conserved activation loop and a PDK1-interacting motif (PIF) ^30^ (Supplementary Fig. 13). Thus, one simple hypothesis is that PDK1 regulates distinct AGC kinase to participate in different life activities. Given the basal localization and the typical auxin defective phenotype, D6PKs from the AGC family were particularly analysed. Morphological analysis showed that *pdk1.1 pdk1.2* resembled the *d6pk-1 d6pkl1-1 d6pkl2-2* (*d0 d1 d2*) triple mutant in various aspects (Fig. 4A and Supplementary Fig. 14), and *pdk1.1 pdk1.2 d0 d1 d2* was similar to the parental lines, indicating that they functioned in the same pathway. Analysis by lipid-protein overlay assays further confirmed a similar lipid binding spectrum for PDK1 and D6PKs (Supplementary Fig. 15). As expected, YFP-D6PK co-localized with mCherry-PDK1s at the basal PM (Fig. 4b, c; the same results were observed with mCherry-D6PKs and Venus-PDK1.1, Supplementary Fig. 16a-c). However, it was noticed that Venus-PDK1s showed a stronger signal in the cytoplasm, than mCherry-D0∼D3 (Supplementary Fig. 16a-b), suggesting additional functions of PDK1s beyond the one at PM. In addition, Western blot analysis showed that D6PK exhibited a mobility shift due to phosphorylation, whereas this was impaired in *pdk1.1 pdk1.2* (Fig. 4d), indicating that PDK1s are crucial for the phosphorylation and activation of D6PK *in planta*.

**Figure 4.**
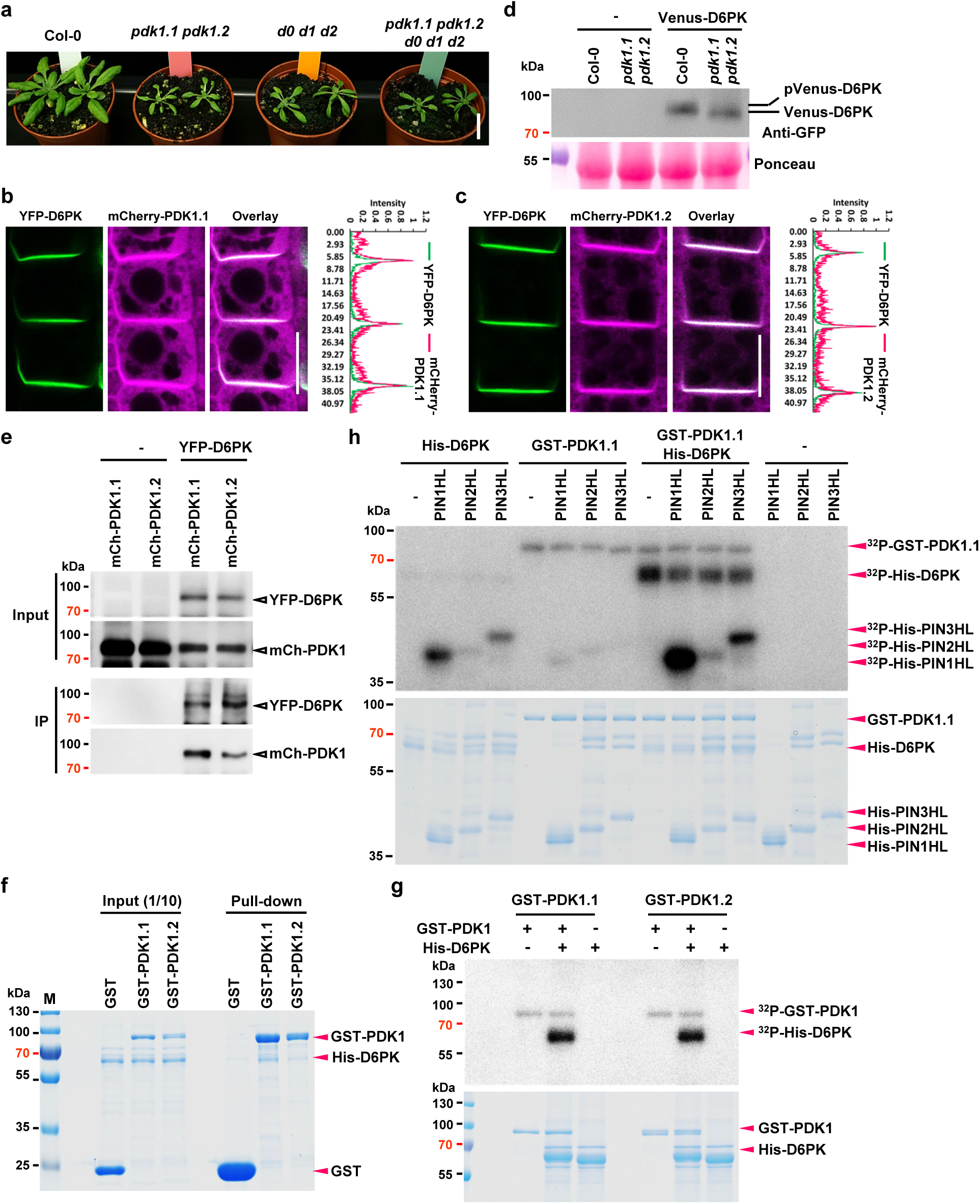
PDK1.1 and PDK1.2 regulate plant growth and development through phosphorylating D6PK/D6PKLs. a. Triple mutant *d0 d1 d2* phenocopied *pdk1.1 pdk1.2*. The quintuple mutant of *pdk1.1 pdk1.2 d0 d1 d2* exhibited a similar phenotype as parental lines. A representative photo of 25-day-old adult plants under long-day conditions was shown. Scale bar, 2 cm. b-c. mCherry-PDK1.1 (b) and mCherry-PDK1.2 (c) colocalized with YFP-D6PK at the basal side of PM in root epidermal cells. Scale bars, 10 µm. d. Phosphorylation of Venus-D6PK was impaired in *pdk1.1 pdk1.2*. Five-day-old seedlings of *35S::Venus-D6PK (Col-0)* and *35S::Venus-D6PK* (*pdk1.1 pdk1.2*) were used for protein extraction and Western blot analysis. Upper panel, Western blot with a GFP antibody; lower panel, Ponceau staining. e. Co-IP assay revealed the interaction of mCherry-PDK1s and YFP-D6PK. Crossing lines *35S::YFP-D6PK*; *RPS5A::mCherry-PDK1.1* and *35S::YFP-D6PK; RPS5A::mCherry-PDK1.2* were used for Co-IP assays. Transgenic lines *RPS5A::mCherry-PDK1.1* and *RPS5A::mCherry-PDK1.2* were used as controls. Anti-GFP beads were used to immunoprecipate YFP-D6PK. The “Input” and “IP” samples were detected with a GFP antibody and a RFP antibody sequentially. f. GST Pull-down assays revealed that both GST-PDK1.1 and GST-PDK1.1 directly interact with His-D6PK. Recombinant GST (as control), GST-PDK1.1, and GST-PDK1.1 proteins (100µg each) were incubated with His-D6PK (100 µg, 1 µg/ µL) with 100 µL GST binding resin for 30 min, and then washed three times before being subjected to SDS-PAGE. Gels were visualized by coomassie brilliant blue (CBB) staining. 1/10 of each reaction mixture before washing was used as “Input”. g. *In vitro* kinase assay revealed that GST-PDK1.1 and GST-PDK1.2 directly phosphorylated D6PK. Recombinant His-D6PK, GST-PDK1.1 and GST-PDK1.2 were used for *in vitro* phosphorylation reactions, which were initiated by adding [^32^P]-ATP. Upper panel, autoradiography of ^32^P; lower panel, CBB staining. h. *In vitro* kinase assay using [^32^P]-ATP revealed that GST-PDK1.1-conducted phosphorylation of D6PK facilitated its activity towards phosphorylation of PIN-HL (recombinant His-PIN1∼3-HL proteins). Upper panel, autoradiography; lower panel, CBB staining.

**Figure 5.**
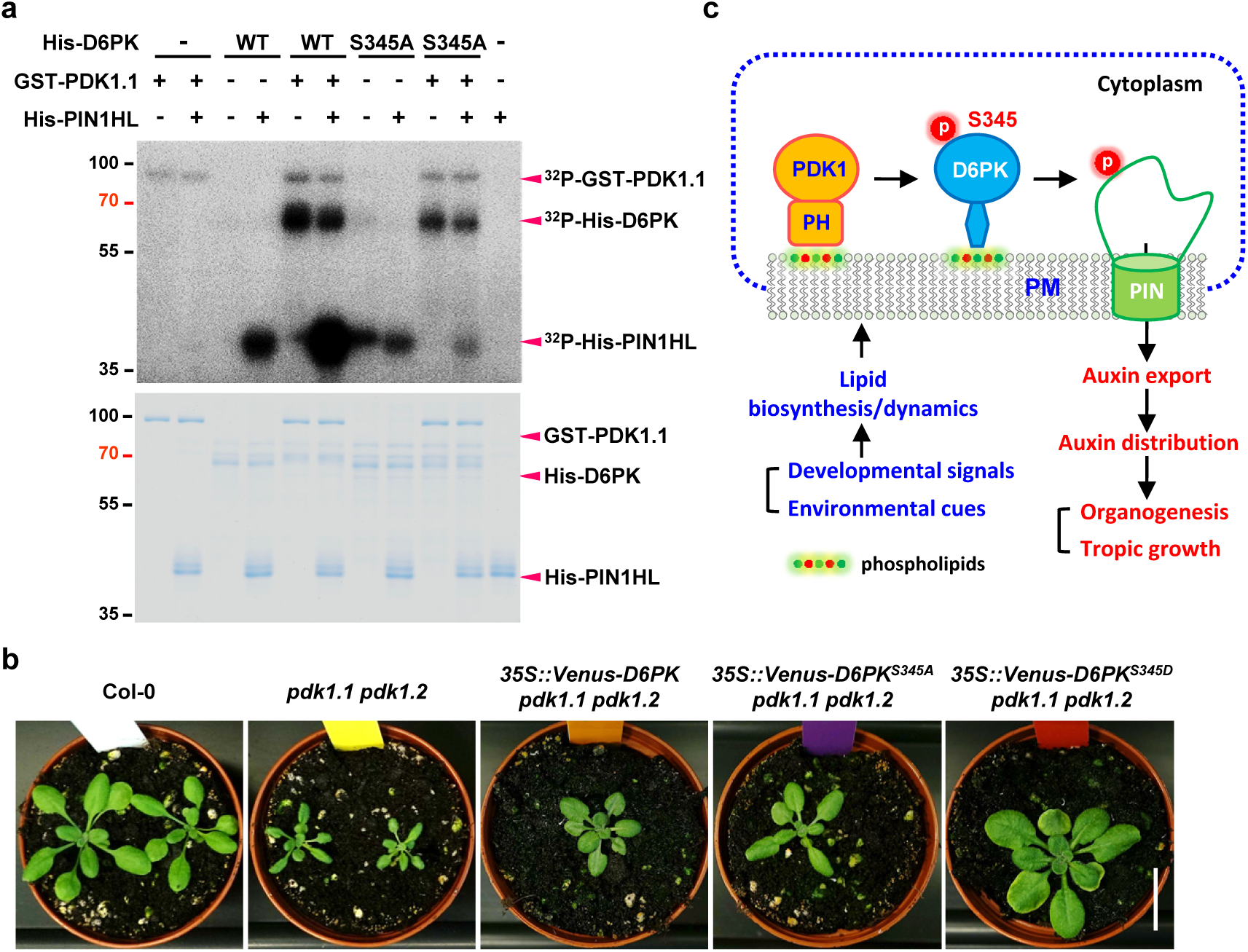
Phospholipids-PDK1-D6PK cascade regulates PIN-dependent polar auxin transport and plant growth. a. *In vitro* kinase assay revealed that GST-PDK1.1-conducted phosphorylation at Ser345 was required for the activation of D6PK, towards His-PIN1-HL phosphorylation. Upper panel, autoradiography; lower panel, CBB staining. b. Overexpression of the phosphomimic D6PK^S345D^, but not the wild-type D6PK and phosphodead D6PK^S345A^ versions, largely derepressed the growth attenuation of *pdk1.1 pdk1.2*. A representative photo of 30-day-old plants is shown. Scale bar, 2 cm. c. In root columella cells or at the basal side of PM in cells of the vascular system, local biosynthesis of an array of phospholipids, including PI monophosphates (PI3P, PI4P, PI5P), PI bisphosphates [PI(4,5)P_2_, PI(3,4)P_2_, PI(3,5)P_2_] and PA in response to distinct developmental signals or environmental cues, recruits PDK1 and D6PK to a membrane nanodomain or microdomain. Thereafter, PDK1 interacts with and phosphorylates D6PK at Ser345, the activated D6PK hence phosphorylates and stimulates the activity of auxin efflux carriers (PINs), facilitating the rootward auxin transport and leading to the fine-tuned plant growth the development.

A co-immunoprecipitation (Co-IP) assay revealed that YFP-D6PK interacted with both mCherry-PDK1.1 and mCherry-PDK1.2 *in vivo* (Fig. 4e). Further GST pull-down and *in vitro* kinase assays using recombinant GST-PDK1.1/2 and His-D6PK proteins revealed that both GST-PDK1.1 and GST-PDK1.2 interacted with and phosphorylated His-D6PK directly (Fig. 4f, g). On the other hand, the band intensities of GST-PDK1s did not change when incubated with or without D6PK, indicating that D6PK could not phosphorylate PDK1.1 or PDK1.2. D6PK has been well established as an essential activator of PIN proteins through direct phosphorylation^9, 10^. With recombinant His-tagged PIN1∼3 hydrophilic loops (His-PIN1∼3HL), further kinase assays were performed to test if PDK1-mediated phosphorylation affected the D6PK activity towards PIN-HL phosphorylation. His-D6PK could phosphorylate His-PIN-HLs, whereas GST-PDK1.1 (Fig. 4h) and GST-PDK1.2 (Supplementary Fig. 16d) could only slightly phosphorylate His-PIN1HL and His-PIN2HL *in vitro*. Notably, presence of GST-PDK1.1 or GST-PDK1.2 together with His-D6PK resulted in a much higher level of PIN-HL phosphorylation than D6PK alone (Fig. 4h and Supplementary Fig. 16d). Intriguingly, although His-PIN2-HL is not a preferential substrate of His-D6PK, after being phosphorylated by GST-PDK1s, His-D6PK exhibited a markedly higher activity towards PIN2-HL phosphorylation (Fig. 4h and Supplementary Fig. 16d).

PDK1 interacts with the PIF motif of the AGC kinase substrates, and thereby phosphorylates the activation loop at a conserved phosphosite^21, 30^, here Ser345 for D6PK (Supplementary Fig. 13). To test if D6PK plays a major role downstream of PDK1, we generated point mutants of D6PK, the phosphodead Ser345Ala (short as S345A) and phosphomimic Ser345Asp (S345D). *In vitro* kinase assays revealed that the S345A mutation largely decreased the phosphorylation of D6PK by PDK1s (Fig. 5a, Supplementary Fig. 16e, and Supplementary Fig. 17a-d), indicating that S345 one key phosphosite of D6PK by PDK1s. Furthermore, the S345A mutation abolished the D6PK activation by PDK1 towards PIN-HL phosphorylation (Fig. 5a, Supplementary Fig. 16e, and Supplementary Fig. 17a-d). Overexpression of Venus-fused D6PK variants, including wild-type, S345A and S345D, in *pdk1.1 pdk1.2* plants showed that *35S::Venus-D6PK^S345D^* largely rescued the growth defects, but *35S::Venus-D6PK* or *35S::Venus-D6PK^S345A^* did not (Fig. 5b), confirming the critical function of S345 in D6PK activation by PDK1 *in vivo*. In addition, both Venus-D6PK and Venus-D6PK^S345D^ showed basal localization at PM, whereas Venus-D6PK^S345A^ exhibited more intracellular signal in Col-0 (Supplementary Fig. 17e). In *pdk1.1 pdk1.2*, both Venus-D6PK and Venus-D6PK^S345A^ showed an increase in the intracellular signal, whereas Venus-D6PK^S345D^ kept more signal at PM, suggesting that phosphorylation at S345 was important for the PM targeting of D6PK. Taken together, PDK1 phosphorylates D6PK at S345, thereby increases its activity towards PIN proteins, as well as stabilizing its localization at PM. Additionally, more kinase assays with D6PKL2 further confirmed that PDK1 also phosphorylated and activated D6PKL2, with PIN-HLs as substrates (Supplementary Fig. 18a, b).

## Discussion

Plant cells perceive extracellular signals via plasma membrane-localized sensors, to coordinate with developmental programming or in response to environmental stimuli. In addition to the role as membrane constituents, phospholipids also function as signals to transduce local membrane information to downstream effectors^47^. Both internal developmental signals and external environment cues have been reported to regulate membrane lipid dynamics, thereby modulating the downstream signal transduction^5, 45, 47, 48^. Multiple lines of genetic evidence suggest that the local membrane status regulates polar localization and recycling of PIN proteins. For example, mutants defective in phospholipid biosynthesis or metabolism exhibit strong defects in various growth and developmental processes, for which mislocalization of PIN proteins account for many of those phenotypes^19, 48–53^. Our studies uncover a novel layer of participation of lipids, in regulating PIN transporters, at the level of activity via polarizing a PDK1-mediated phosphorylation cascade.

In animals, PDK1 plays a critical role in most life activities, as a master activator of the AGC kinases, and dysfunction of AGC kinases in humans cause different diseases, including cancer and diabetes^21^. For instance, PDK1 can phosphorylate and activate one key substrate, AKT, to promote cell proliferation and diverse metabolic processes^21^. Plants AGC family comprises 39 members, having evolved unique biological functions, via phosphorylating distinct substrates. Our findings establish a central role of PDK1 in transducing a local lipid signal to auxin transport activity (Fig. 5c): Local biosynthesis of certain phospholipids recruits both PDK1 and D6PK at the basal side of PM, where PDK1 phosphorylates and stimulates D6PK to amplify the signal. The activated D6PK then phosphorylates the basal-localized, plant-specific, PIN proteins, and thereby increases the export activity of PIN auxin efflux carriers, eventually enabling the rootward polar auxin transport. Notably, the PDK1-D6PK module, analogous to the PDK1-AKT pathway ^21^ of mammalian cells, acts as a phosphoswitch to strictly control the directional auxin flow and the subsequent developmental processes. Interestingly, except for the PM localization, PDK1 also locates at the cytoplasm. Whether PDK1 regulates more substrates, other than D6PKs, to participate in additional life activities, needs further investigation.

## Methods

### Plant materials and growth conditions

*Arabidopsis thaliana* (L.) mutants and transgenic lines are all in Columbia-0 (Col-0) background. The marker lines *pPIN1::PIN1-YFP*^54^, *pPIN1::PIN1-GFP*^55^, *pPIN2::PIN2-GFP*^56^, *pPIN3::PIN3-GFP*^34^, *DR5rev::GFP*^6^, and *35S::YFP-D6PK*^9^ were described previously. The mutant *d6pk-1 d6pkl1-1 d6pkl2-2*^9^ was also previously reported. The *pdk1.1* (SALK_113251) and *pdk1.2* (SALK_017433) mutants were obtained from the *Arabidopsis* Biological Resource Centre (ABRC) and verified by PCR. All the plant lines, including mutants, reporters and crosses, are listed in Supplementary Table 2. The primers used to genotype T-DNA mutants are listed in Supplementary Table 4.

The floral dip method^57^ was used for *Arabidopsis* transformation with an *Agrobacterium tumefaciens* strain GV3101. For phenotyping of seedlings or pharmacological experiments, surface-sterilized seeds by chlorine gas were sown on 0.5× Murashige and Skoog (MS) medium supplemented with 1% sucrose and 0.8% phytoagar (pH 5.9), stratified at 4°C for 2 days, and then grown in a phytotron at 21°C with a long-day photoperiod (16h light/8h dark). For dark treatment, plates were covered with aluminium foil. For adult plants, 7-day-old seedlings were transferred to soil and grown under long-day (16h light/8h dark) conditions. Light sources used were GreenPower LED production modules [in deep red (660 nm)/far red (720 nm)/blue (455 nm) combination, Philips], with a photon density of approximately 140 µmol/m^2^/s ± 20%.

### Molecular cloning

Restriction enzyme digestion or Gateway cloning was used to make constructs, for recombinant protein expression in *E.coli* or transgenic plants generation. For restriction enzyme digestion, the restriction enzyme sites were added to DNA fragments by PCR, and were then cloned into the corresponding plasmids. For Gateway cloning, DNA fragments were firstly cloned into entry vectors by BP reaction, and were then cloned into destination vectors by LR reaction. All the reagents, including restriction enzymes and Gateway clonases, are listed in Supplementary Table 3. The primers used for molecular cloning are listed in Supplementary Table 4. The resultant plasmids from this study are listed in Supplementary Table 5.

### Quantitative RT-PCR (qRT-PCR) analysis

qRT-PCR was used to examine the transcripts of *PDK1.1* and *PDK1.2* in *pdk1.1* and *pdk1.2* mutants respectively, with *ACTIN7* (*AT5G09810*) as an internal reference. In detail, total RNA was extracted from indicated tissues using TRIzol (Invitrogen), and DNase was added to digest genomic DNA. Then 1 µg of RNA sample was used for reverse transcription (Takara). The resultant cDNA of corresponding genes and *ACTIN7* was analysed using the SYBR Premix Ex Taq (Takara) with a Bio-Rad CFX Connect Real-Time System. Relative transcript level of examined genes was normalized by *ACTIN7*, and calculated by setting wild-type or a certain tissue as “1”, and was finally presented as average ± standard deviation (s.d.) from three biological replicates.

### Promoter::β-glucuronidase (GUS) Histochemical staining

For the promoter-GUS fusion studies, 1.2-kilobases (kb) and 1.1-kb genomic DNA fragments containing the promoter regions of the *PDK1.1* and *PDK1.2* genes were amplified by PCR respectively, and were subcloned into a modified pCambia1300 binary vector^58^ (The primers used for molecular cloning were listed in Supplementary Table 3). Positive transgenic lines were stained at 37°C for 1 h, in GUS staining solution [0.5 mg/mL 5-bromo-4-chloro-3-indolyl-β-d-glucuronic acid (X-Gluc, added just before staining with a 80× stock in DMSO), 0.5 mM potassium ferricyanide (K_4_[Fe(CN)_6_]·3H_2_O), 0.5 mM potassium ferrocyanide (K_3_[Fe(CN)_6_]), 0.1% (v/v) Triton X-100, 10 mM EDTA (ethylenediaminetetraacetic acid), and 0.1 M sodium phosphate (NaH_2_PO_4_), pH 7.0]^59^. More than ten independent transgenic lines for each transgene were tested for GUS activity at the seedling stage, and three representatives of them were further analysed in detail for different tissues and stages, and they all showed similar patterns.

### Scanning electron microscopy (SEM)

For imaging seedling cotyledons and pavement cells, the samples were mounted on an ironic platform, frozen in liquid nitrogen, and then observed by a JSM-B360LV scanning electron microscope (JEOL, Japan), with a cryo adaptor.

### Pharmacological treatments

For pharmacological treatments, seeds were sown on MS plates with indicated chemicals, such as ACC (Sigma). After 2-day stratification at 4°C, plates were transferred to a growth chamber under conditions as mentioned in the “Plant material and growth conditions” section, for 7 days or 10 days, respectively. The phenotype was then analysed by Image J (NIH)^60^.

For short-term treatment to study the subcellular localization of Venus-PDK1.1, 4-day-old *35S::Venus-PDK1.1* seedlings were incubated in liquid MS medium with DMSO (as the solvent control), 33 µM Wortmannin, 30 µM PAO, 50 µM LY294002, 5 µM U-73122, 0.8% 1-butanol, or 50 µM BFA, for 30 min. Afterwards, the samples were imaged with CLSM. All the reagents used in this study are listed in Supplementary Table 2.

### Imaging with confocal laser scanning microscopy (CLSM)

Fluorescence imaging was performed using a Zeiss LSM800 CLSM equipped with a GaAsP detector. The manufacturer’s default settings were used for imaging Green florescent protein (GFP) (excitation, 488 nm; emission, 495-545 nm)-, YFP (excitation, 514 nm; emission, 524-580 nm), Venus (excitation, 514 nm; emission, 524-580 nm)-, and mCherry (excitation, 561 nm; emission, 580-650 nm)-tagged proteins respectively. For propidium iodide (PI) staining, excitation, 561 nm; emission, 580-680 nm. All images were obtained in 8-bit depth, 2× line averaging.

### Propidium iodide staining

For PI (Sigma, P4170) staining, 4-day-old seedlings were stained with PI at the final concentration of 10 mg/ml in H_2_O for 1 min before imaging with CLSM.

### Statolith starch staining with Lugol solution

5-day-old seedlings from indicated genotypes were stained with Lugol solution for 2 min, and were then washed with liquid MS medium for 2 min. The samples were mounted with a clearing solution (30 mL H_2_O, 10 mL glycerol, 80 g chloral hydrate, with 100 mL in total). Finally, root tips were imaged with a differential interference contrast (DIC) microscopy (Nikon SMZ1500).

### Vasculature pattern analysis with microscope

12-day-old seedlings were firstly incubated in 100% ethanol for 30 min to remove the chlorophylls, and then gradually exchanged with 75% (v/v) ethanol, 50% (v/v) ethanol, and H_2_O, with each incubation for 30 min respectively. The samples were mounted with a clearing solution (30 mL H_2_O, 10 mL glycerol, 80 g chloral hydrate, with 100 mL in total), and the incubated for at least 1 h, and finally imaged with a DIC microscopy (Nikon).

### Immunolocalization analysis

PIN immunolocalization experiments with *Arabidopsis* primary roots were carried out as reported previously^33^. In detail, 4-day-old seedlings of Col-0, *pdk1.1 pdk1.2*, and *XVE::Venus-PDK1.1* (treated with or without β-Estradiol) were firstly fixed with FAA (formaldehyde-acetic acid-ethanol) solution. A rabbit anti-PIN1 antibody^61^ (1:1000) and a rabbit anti-PIN2 antibody ^62^ (1:1000) were used to detect PIN1 and PIN2 respectively. A secondary goat anti-rabbit antibody coupled to Cy3 (Sigma-Aldrich) was further used for the immunofluorescence reaction. Samples were imaged with a Zeiss LSM800 CLSM.

### Image analysis and morphological analysis

For root phenotype analysis, photos of plates were taken with a scanner (Epson Dual lens V800) and/or a camera (Sony A600 with a macro lens). The lateral root numbers were counted. The primary root length, apical hook curvature, hypocotyl growth direction, or the root tip angle were measured with and analysed with Image J or Fiji^60^.

### Auxin transport assays in *Arabidopsis* hypocotyls

The basipetal transport of auxin in intact hypocotyls of etiolated seedlings with [^3^H]-IAA was performed as reported previously^63^, with a few modifications. 6-day-old etiolated Col-0, *pdk1.1*, *pdk1.2*, and *pdk1.1 pdk1.2* seedlings were transferred to new MS plates, with 15 seedlings as one biological replicate, and 6 replicates for each genotype. Agarose droplets with [^3^H]-IAA (PerkinElmer) were prepared in MS medium with 1.25% agarose and 500 µM [^3^H]-IAA (5.8 µL in 10 mL this solution), 10 µL/droplet on a Petri-dish, in darkness to prevent the degradation of [^3^H]-IAA. The seedlings were decapitated and then placed with a [^3^H]-IAA droplet at the shootward end (plus three groups with [^3^H]-IAA droplets at the roodward end as controls). After incubation for 6 hours in the dark, the lower part of the hypocotyls were cut (the roots and the upper parts of the hypocotyls having direct contact with the [^3^H]-IAA droplets were removed) and collected into a 2 mL Eppendorf tube (with two iron beads inside), and then frozen in liquid nitrogen. The samples were ground completely and then homogenized in 1 mL scintillation liquid (PerkinElmer). These samples were incubated overnight to allow the radioactivity to evenly diffuse into the scintillation cocktail. Finally the radioactivity was measured with a scintillation counter (Hidex 300XL), with each sample counted for 300 secs, 3 times. Three samples with only the scintillation solution were used as background controls, and for subtraction in data analysis.

### Protein extraction and Western blot analysis

To examine the protein levels of Venus-tagged proteins, approximately 100 mg of seedlings (7 days old) were frozen in liquid nitrogen, ground totally, and homogenized in 100 μL plant extraction buffer (20 mM Tris-HCl, pH 7.5, 150 mM NaCl, 0.5% Tween-20, 1 mM EDTA, 1 mM DTT) containing a protease inhibitor cocktail (cOmplete, Roche) and a protein phosphatase inhibitor tablet (PhosSTOP, Roche). After addition of half a volume of SDS loading dye, the samples were boiled at 65°C for 5 min, resolved by 10% SDS-PAGE and transferred to a PVDF membrane by wet blotting. The membranes were incubated with a mouse anti-GFP HRP-conjugated antibody. mCherry proteins were detected by a rat anti-RFP antibody (5F8-100, 1:2,000, Chromotek), and then with a goat anti-rat IgG HRP-conjugated secondary antibody (AP202P, 1:10,000, Sigma). HRP activity was detected by the Supersignal Western Detection Reagents (Thermo Scientific).

### Co-immunoprecipitation (Co-IP) assay

To examine the interaction between YFP-D6PK and mCherry-PDK1s *in planta*, crossing lines *35S::YFP-D6PK*; *RPS5A::mCherry-PDK1.1* and *35S::YFP-D6PK; RPS5A::mCherry-PDK1.2* were used for Co-IP assays using a µMACS Epitope Tag Protein Isolation Kit (Miltenyi Biotec), with *RPS5A::mCherry-PDK1.1* and *RPS5A::mCherry-PDK1.2* transgenic lines as controls. In detail, ∼100 mg of 7-day-old seedlings were homogenized into 200 µL protein extraction buffer [20 mM Tris-HCl (pH 7.5), 150 mM NaCl, 0.5% Triton X-100, and 1 mM DTT] containing a protease inhibitor cocktail (cOmplete, Roche) and a protein phosphatase inhibitor tablet (PhosSTOP, Roche). After centrifuge, 200 µL supernatant of each sample was transferred to a new tube for incubation with 50 µL µMACS anti-GFP magnetic microbeads, with 20 µL lysate spared as the “Input” samples. The reaction samples were incubated at 4°C for 1h, and were then loaded onto a µ column on a magnetic holder. The beads were washed 5 times with the supplied wash buffer [150 mM NaCl, 1% Igepal CA-630, 0.5% sodium deoxycholate, 0.1% SDS, 50 mM Tris HCl (pH 8.0)], and were then eluted with elution buffer [50 mM Tris HCl (pH 6.8), 50 mM DTT, 1% SDS, 1 mM EDTA, 0.005% bromphenol blue, 10% glycerol]. Protein samples were separated by 10% SDS-PAGE, and were eventually detected by a mouse anti-GFP HRP-conjugated antibody and a RFP antibody sequentially.

### *In vivo* protein phosphorylation analysis

The phosphorylation status of Venus-D6PK was examined with a mobility shift assay by SDS-PAGE. ∼100 mg of *35S::Venus-D6PK (Col-0)* and *35S::Venus-D6PK (pdk1.1 pdk1.2)* seedlings (7 days old) were homogenized into 100 μL plant extraction buffer (20 mM Tris-HCl, pH 7.5, 150 mM NaCl, 0.5% Triton X-100, and 1 mM DTT) containing a protease inhibitor cocktail (cOmplete, Roche) and a protein phosphatase inhibitor tablet (PhosSTOP, Roche). Protein samples were analysed by Western blot with a mouse anti-GFP HRP-conjugated antibody.

### Recombinant protein expression and purification from *E. coli*

Recombinant proteins, including GST, GST-PDK1.1, GST-PDK1.2, GST-D6PKL1, His-D6PK, His-D6PK^S345A^, His-D6PK^S345D^, His-D6PKL2, His-PDK1.1N, His-PDK1.1C, His-PDK1.2N, His-PDK1.2C, His-PIN1HL, His-PIN2HL, and His-PIN3HL were expressed in an *E. coli* strain BL21 (DE3) with induction by 0.5 mM IPTG (Isopropyl β-D-1-Thiogalactopyranoside, 16°C, 12 h) and then purified using Ni-NTA His binding resin (Thermo Scientific) according to the manufacturer’s manual. The eluted protein samples were further purified with size exclusion chromatography, with a Superdex 200 (10/30L) increase column (GE Healthcare), on an ÄKTA pure chromatography system. Protein samples were collected every 500 μL, and then checked by SDS-PAGE, visualized by the Coomassie brilliant blue staining (Bio-Safe™ Coomassie Stain #1610786, Bio-Rad). The protein concentration was qualified with the Bradford method (Quick Start™ Bradford Reagent, Bio-Rad).

### Lipid-protein binding assay with PIP strips

Lipid binding assays, namely lipid-protein blot overlay assays^18^, were performed with PIP strips or PIP arrays (P-6001 or P-6100, Echelon Bioscience) and recombinant proteins expressed and purified from *E. coli*. Nitrocellulose membranes containing immobilized lipids (PIP strips or PIP arrays) were incubated in blocking solution [1×TBST (50 mM Tris-HCl, 150 mM NaCl, 0.1% Tween-20, pH 7.6) + 3% BSA] for 1 h. Membranes were then incubated for 2 h in 10 mL of 1×TBST containing 20 µl of recombinant proteins, His-PDK1.1, His-PDK1.2, His-PDK1.1N, His-PDK1.2N, His-PDK1.1C, His-PDK1.2C, His-D6PK, His-D6PKL2, GST-D6PKL1, and GST (control). After washing (5 min × 3 times) using 1×TBST, membranes were incubated for 2 h at room temperature with an anti-His HRP-(horseradish peroxidase) conjugated antibody (1:4000; Agrisera) or an anti-GST HRP-conjugated antibody (1:4000; Agrisera) accordingly diluted in blocking solution, and finally rinsed three times with 10 mL 1×TBST respectively before checking by chemiluminescence. HRP activity was detected by Supersignal Western Detection Reagents (Thermo Scientific), and was then visualized with an Amersham 600RGB molecular imaging system (GE Healthcare).

### GST Pull-down assay

To detect the interaction between PDK1s and D6PK, recombinant GST, GST-PDK1.1, GST-PDK1.2, and His-D6PK proteins were used to perform the GST pull-down assays. In detail, 1 mL of GST, GST-PDK1.1, and GST-PDK1.2 lysates were incubated with 100 µL GST resin for 30 min in 1.5 mL tubes at 25 °C. Samples were spun at 3,000 rpm for 1 min, and the supernatant was discarded. The pellet was then washed with 500 µL 1×TBS. 100 µL His-D6PK (1 µg/ µL) was added into each tube, plus another 400 µL 1×TBS, and then incubated at 25 °C for 30 min. 30 µL of the reaction mixture was transferred to a new tube, added with 15 µL loading dye as the “inputs”. Afterwards, the samples were spun at 3,000 rpm for 1 min, and washed three times, with 500 µL 1×TBS, 500 µL 1×TBS, and 1mL 1×TBS respectively. The resin was finally re-suspended in 30 µL elution buffer [1×TBS + 10 mM glutathione (GSH)], and then added with 15 µL loading dye. 10 µL samples were separated with 10 % (v/v) SDS-PAGE, and visualized by coomasie staining (Bio-Safe™ Coomassie Stain #1610786, Bio-Rad).

### *In vitro* kinase assay

*In vitro* phosphorylation assays were conducted as previously described^7^. Recombinant GST-PDK1.1, GST-PDK1.2, His-D6PK, His-D6PKL2, His-PIN1HL, His-PIN2HL, His-PIN3HL in the kinase reaction buffer (50 mM Tris-HCl pH 7.5, 10 mM MgCl_2_, 5 mM NaCl, 10 mM ATP (Adenosine 5’-triphosphate), and 1 mM DTT) at the presence of 5 μCi [γ-^32^P]-ATP (NEG502A001MC; Perkin-Elmer) at 25 °C for 90 min. Then, the reactions were terminated by adding SDS loading buffer. The resultant samples were then separated in a 10% (v/v) SDS-PAGE gel, developed with a phosphor-plate overnight, and finally imaged with a Fujifilm FLA 3000 plus DAGE system.

### Accession Numbers

Sequence data from this article can be found in the *Arabidopsis* Genome Initiative databases under the following accession numbers: PIN1 (AT1G73590), PIN2 (AT5G57090), PIN3 (AT1G70940), PIN4 (AT2G01420), PIN7 (AT1G23080), ARF7 (AT5G20730), ARF19 (AT1G19220), PDK1.1 (AT5G04510), PDK1.2 (AT3G10540), D6PK (AT5G55910), D6PKL1 (AT4G26610), D6PKL2 (AT5G47750), D6PKL3 (AT3G27580), PID (AT2G34650), WAG1 (AT1G53700), and WAG2 (AT3G14370).

## Acknowledgement

We thank Dr. Claus Schwechheimer (TUM) and Dr. Ben Scheres (WU) for sharing published plant lines, and Dr. Matouš Glanc for providing pET28a-PIN2/3 plasmids. We gratefully acknowledge the Nottingham *Arabidopsis* Stock Centre (NASC) and the *Arabidopsis* Biological Resource Centre (ABRC) for providing T-DNA insertional mutants. The research leading to these results has received funding from Chinese “Ten-Thousand Talent Program” (to HW) and the European Union’s Horizon2020 program (ERC grant agreement n° 742985 to JF). ST was supported by a European Molecular Biology Organization (EMBO) long-term postdoctoral fellowship (ALTF 723-2015). XZ was supported by a Ph.D. scholarship from China Scholarship Council.

## Author Contributions

S.T., J.F., and H.X. designed experiments. S.T., X.Z., W.K., and X.Y. performed experiments. G.M. provided [^32^P]-ATP and helped with the kinase assays. S.T., J.F., and H.X. analysed and interpreted the data. S.T., J.F., and H.X. wrote the manuscript with inputs from other co-authors, and all authors revised it.

## Data and materials availability

All data and materials necessary to evaluate the conclusions in the paper or the supplementary materials are available.

## Supplementary Information

Supplementary Figures. 1 to 18

Supplementary Tables. 1 to 5

## Figure legends of Supplemental figures

**Supplementary Figure 1.**
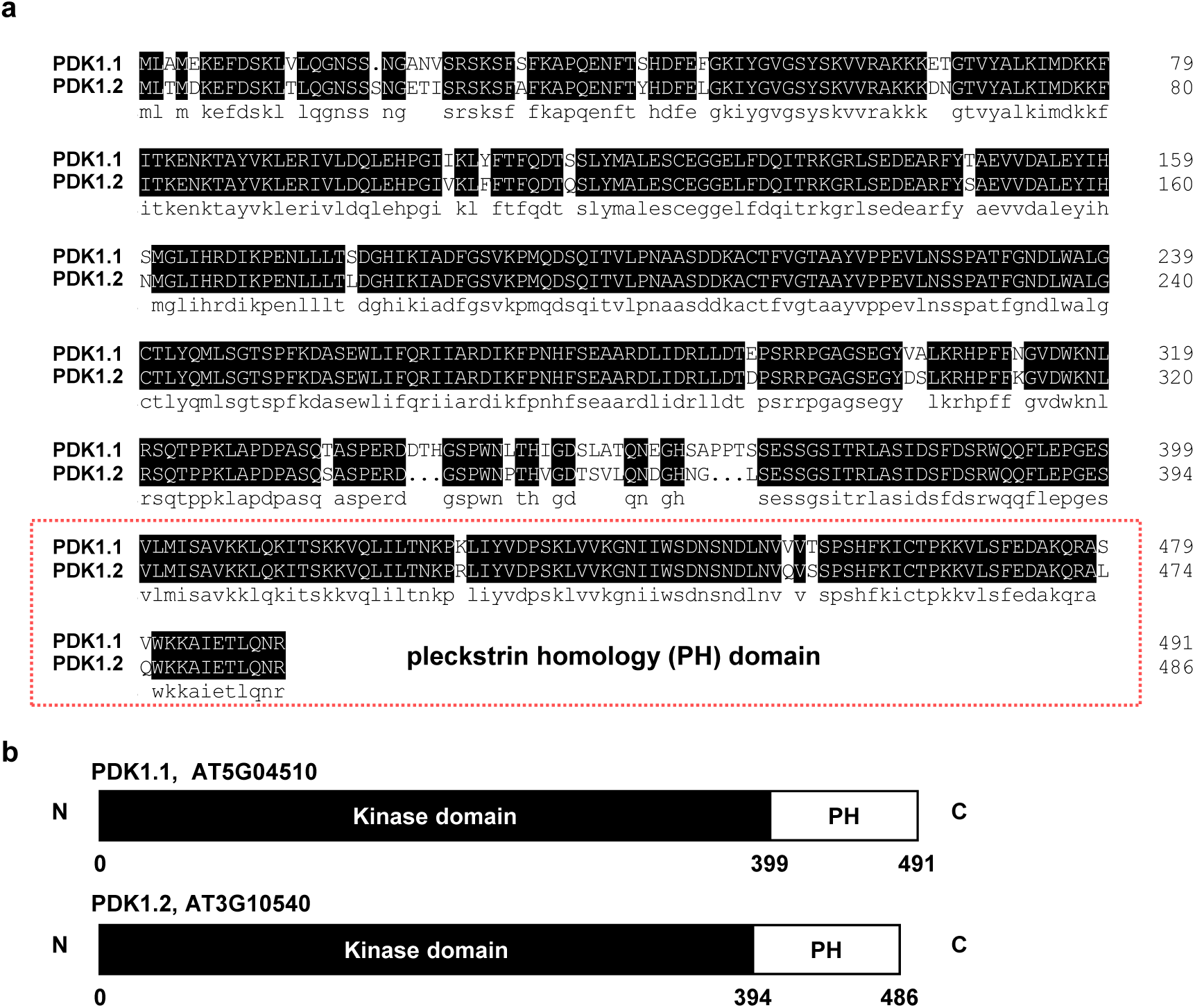
Alignment of the amino acid sequences and schematic graphs of PDK1.1 and PDK1.2. a. Alignment of PDK1.1 and PDK1.2 by the DNAMAN program. The two paralogues show 91.06% identity. The predicted pleckstrin homology (PH) domain is indicated by a red rectangle. b. Schematic graphs of the domain structure of PDK1.1 and PDK1.2 proteins.

**Supplementary Figure 2.**
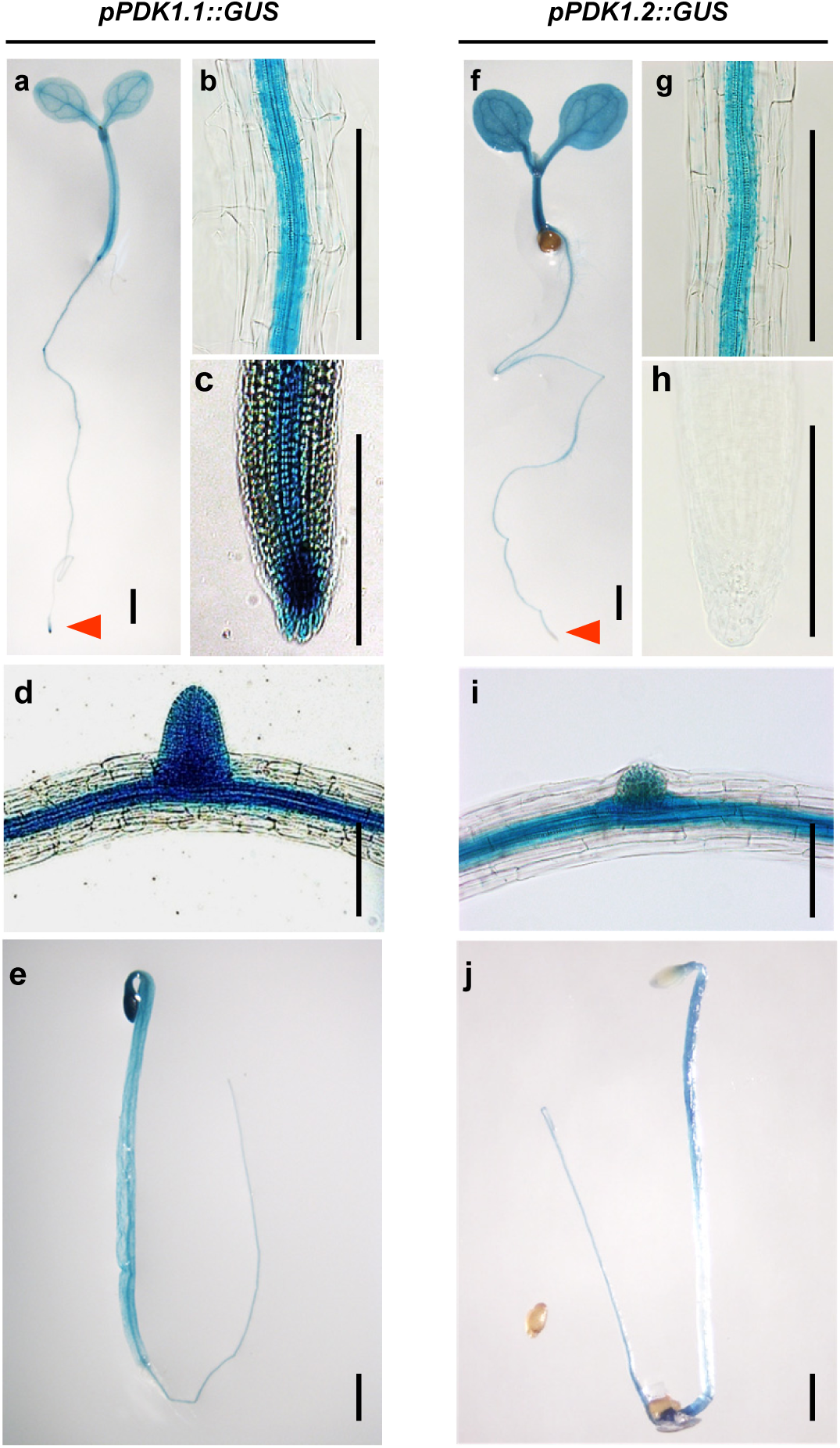
Expression pattern of *PDK1.1* and *PDK1.2*. GUS staining of *pPDK1.1::GUS* and *pPDK1.2::GUS* lines indicated that *PDK1.1* and *PDK1.2* were expressed in the vascular tissues in both roots and shoots at various developmental stages, including young seedlings (a, f; 7 days), root stele (b, g; 7 days), columella cells (c, h, only with expression detected for *PDK1.1*; 7 days), lateral root primordia (d, i; 12 days), and dark-grown seedlings (e, j; 4 days). Representative images of three independent homozygous lines were shown. Scale bars, 1 mm.

**Supplementary Figure 3.**
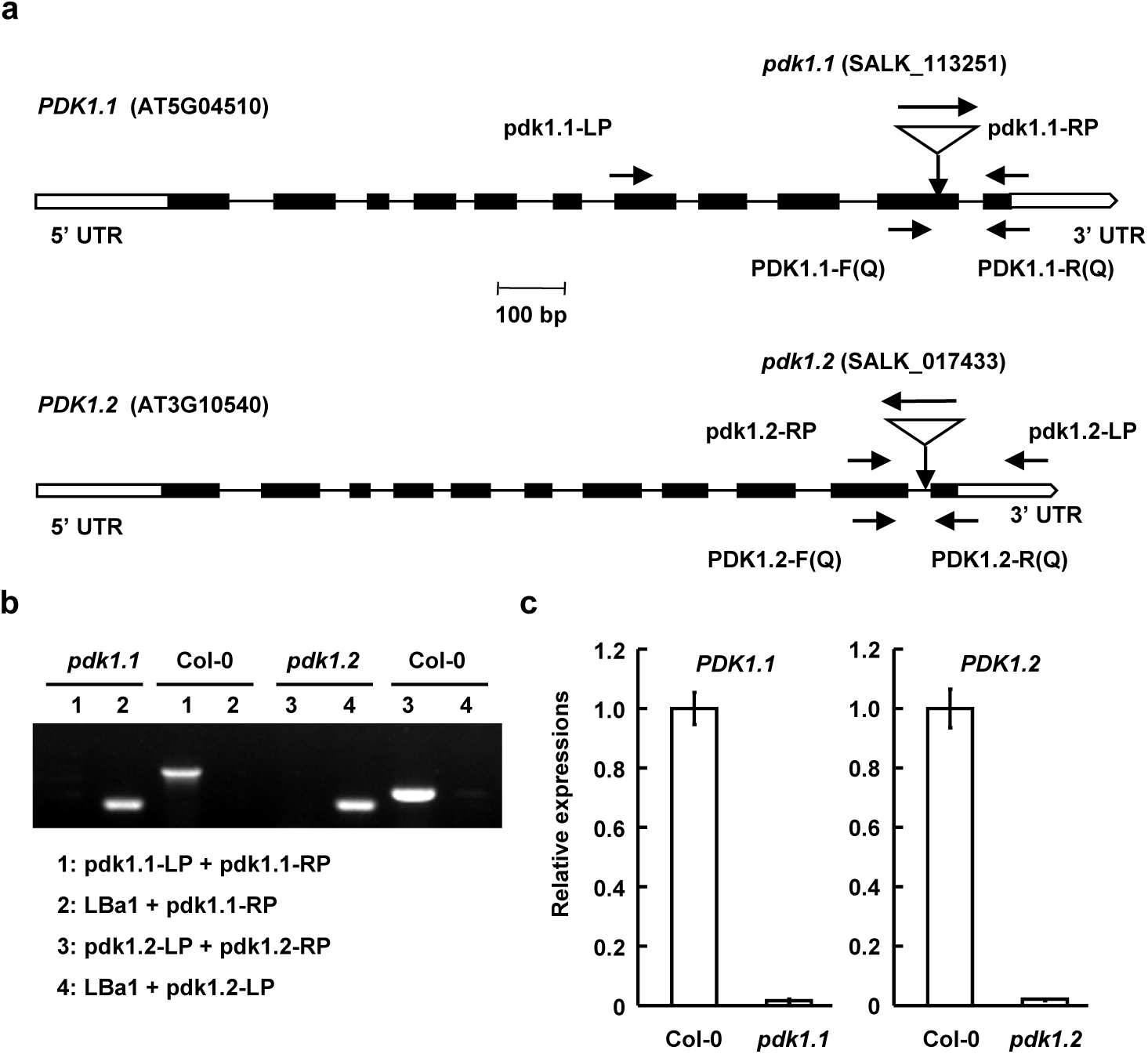
Identification of *Arabidopsis pdk1.1* and *pdk1.2* T-DNA insertional mutants. a. Schematic representation of *PDK1.1* and *PDK1.2* genes and positions of T-DNA insertions for *pdk1.1* and *pdk1.2*. Introns, exons, and non-coding regions are indicated by lines, black, or blank boxes respectively. Positions of primers are indicated. b. Identification of homozygous *pdk1.1* and *pdk1.2* mutants. Genomic DNA of *pdk1.1* and *pdk1.2* mutants was used as templates for PCR amplification. Homozygous lines have a single amplified DNA fragment when using LBa1/pdk1.1-RP or LBa1/pdk1.2-RP primers. c. qRT-PCR analysis confirmed the deficient expression of *PDK1.1* and *PDK1.2* genes in *pdk1.1* and *pdk1.2* mutants, respectively. Total RNA of 7-day-old WT, *pdk1.1* and *pdk1.2* seedlings was extracted, reversely transcribed, and then used for analysis. *ACTIN7* was amplified and used as an internal reference to normalize the expression of *PDK1.1* and *PDK1.2*. The experiments were biologically repeated for 3 times and results were presented as mean ± s.d..

**Supplementary Figure 4.**
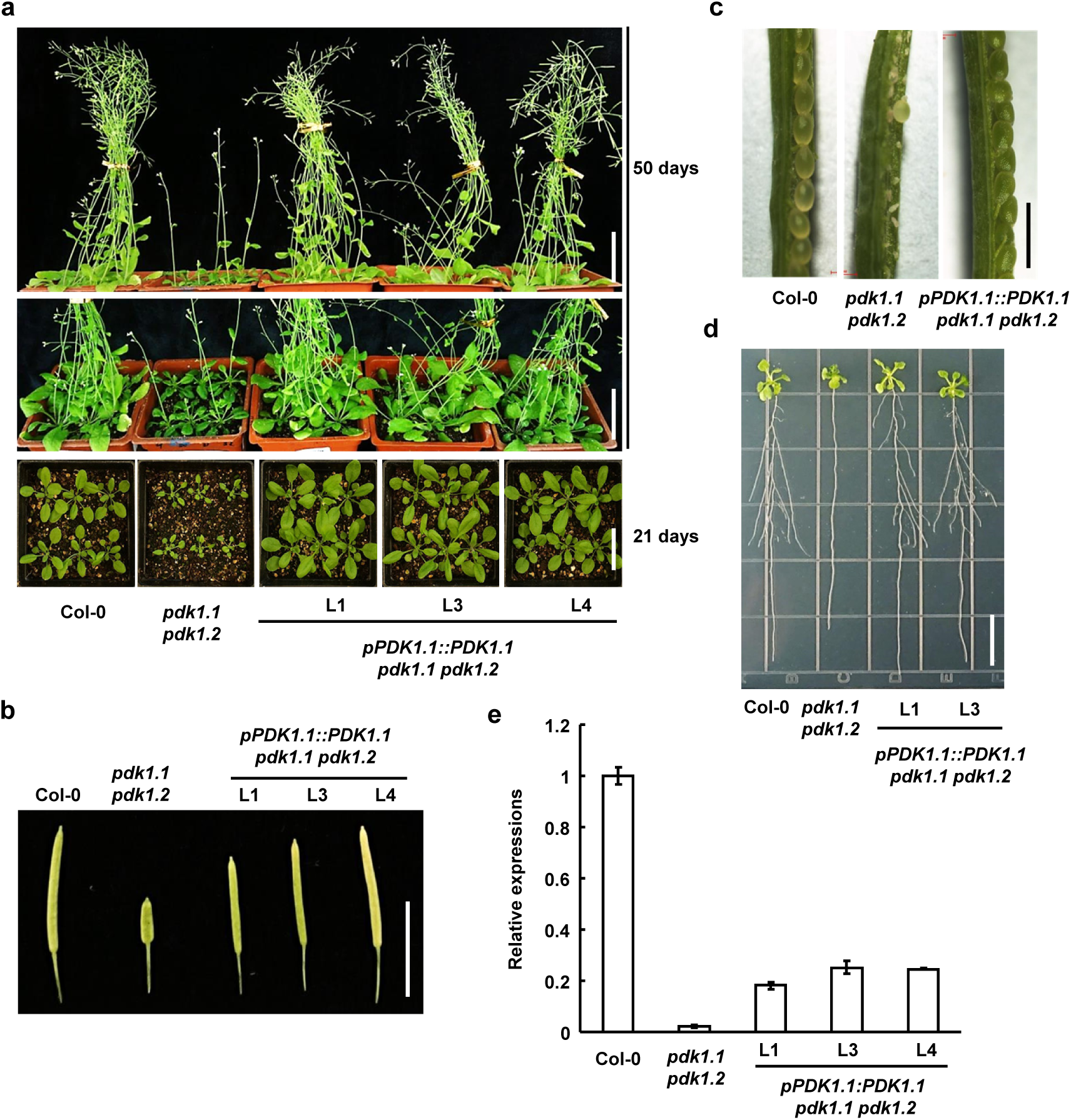
Expression of *PDK1.1* driven by native promoter (*pPDK1.1::PDK1.1*) rescued the defective growth of *pdk1.1 pdk1.2* double mutant. a-d. Complementary expression of *PDK1.1* recovered the reduced growth (a, scale bar, 5 cm), silique shortness (b; scale bar, 1 cm), low fertility (c; scale bar, 1 mm), and defective root elongation and lateral root formation (d, representative images of 2-week-old seedlings were shown; scale bar, 1 cm) of *pdk1.1 pdk1.2*. e. qRT-PCR analysis showed the complementary expression of *PDK1.1* in *pdk1.1 pdk1.2* plants. Total RNA of 7-day-old WT, *pdk1.1 pdk1.2* and transgenic *pdk1.1 pdk1.2* seedlings harbouring *pPDK1.1::PDK1.1* was extracted and used for analysis. *ACTIN7* was amplified and used as an internal reference. Experiments were biologically repeated and data were presented as mean ± s.d. (n = 3).

**Supplementary Figure 5.**
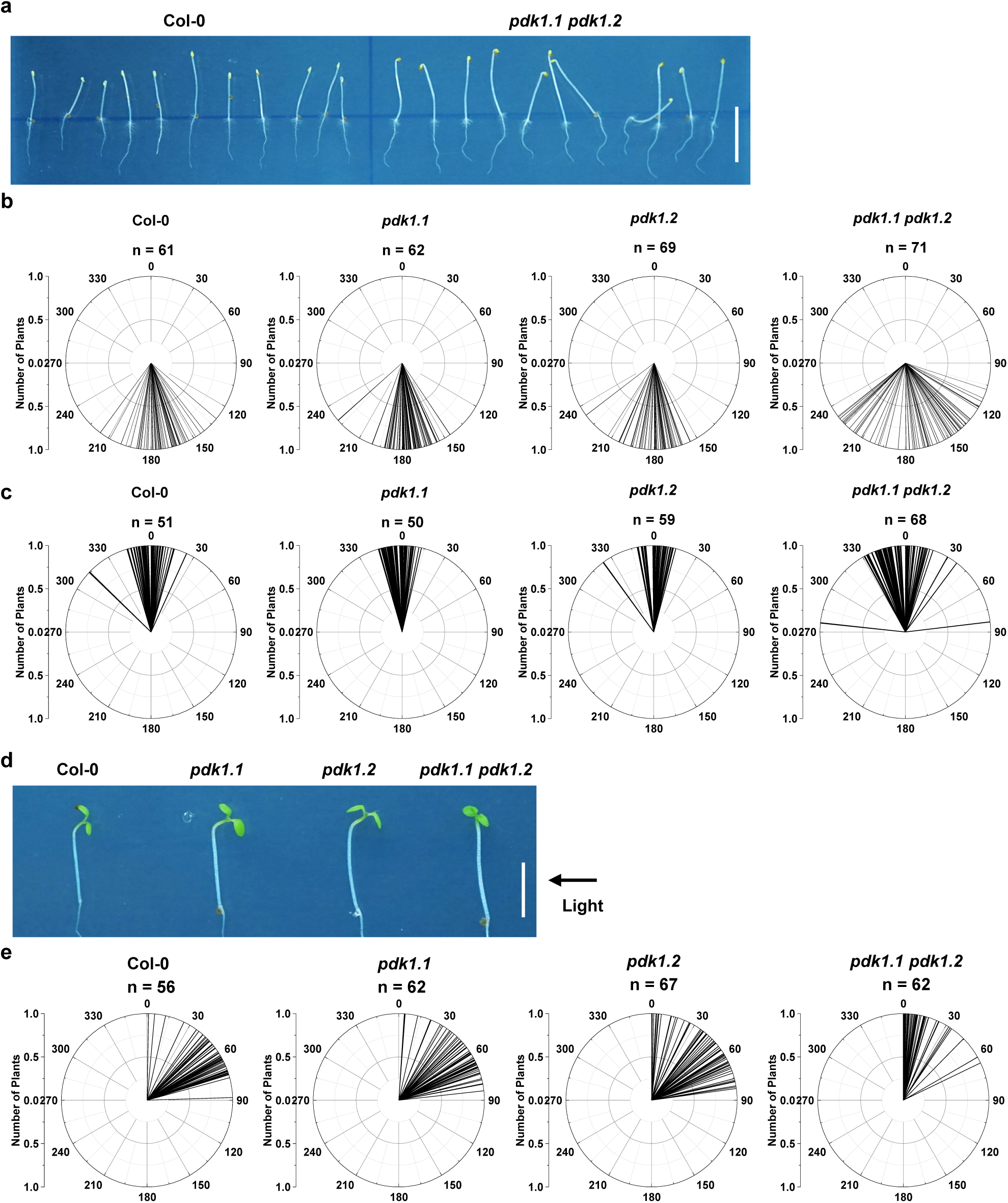
Deficiency of *PDK1.1* and *PDK1.2* impaired the hypocotyl gravitropism under dark and phototropism towards directional light. a. Deficiency of *PDK1.1* and *PDK1.2* impaired root and shoot gravitropic response under dark. Etiolated seedlings of Col-0, *pdk1.1*, *pdk1.2*, and *pdk1.1 pdk1.2* were grown under dark for 90 h and a representative photo was shown. Scale bar, 5 mm. b. Deficiency of *PDK1.1* and *PDK1.2* impaired root gravitropic response under dark. Etiolated seedlings of Col-0, *pdk1.1*, *pdk1.2*, and *pdk1.1 pdk1.2* were grown under dark for 90 h and root tip angles were measured by Image J and data were visualized by polar bar charts. n = 61, 62, 69, and 71, respectively. c. Deficiency of *PDK1.1* and *PDK1.2* led to reduced gravitropic hypocotyls under dark. Etiolated seedlings of Col-0, *pdk1.1*, *pdk1.2*, and *pdk1.1 pdk1.2* were grown under dark for 90 h and hypocotyl angles were measured by Image J and data were visualized by polar bar charts with the Origin program. n = 51, 50, 59, and 68, respectively. d. *pdk1.1 pdk1.2* showed defects in phototropism. Seedlings of Col-0, *pdk1.1*, *pdk1.2,* and *pdk1.1 pdk1.2* were grown under dark for 90 h, exposed to white light for 24 h, and were then subjected to directional white light in a box covered with aluminium foil from the other sides. Scale bar, 5 mm. e. Deficiency of *PDK1.1* and *PDK1.2* impaired hypocotyl phototropic response towards direction light source. As indicated in (c), hypocotyl angles were measured by Image J and data were visualized by polar bar charts. n = 56, 62, 67, and 62, respectively.

**Supplementary Figure 6.**
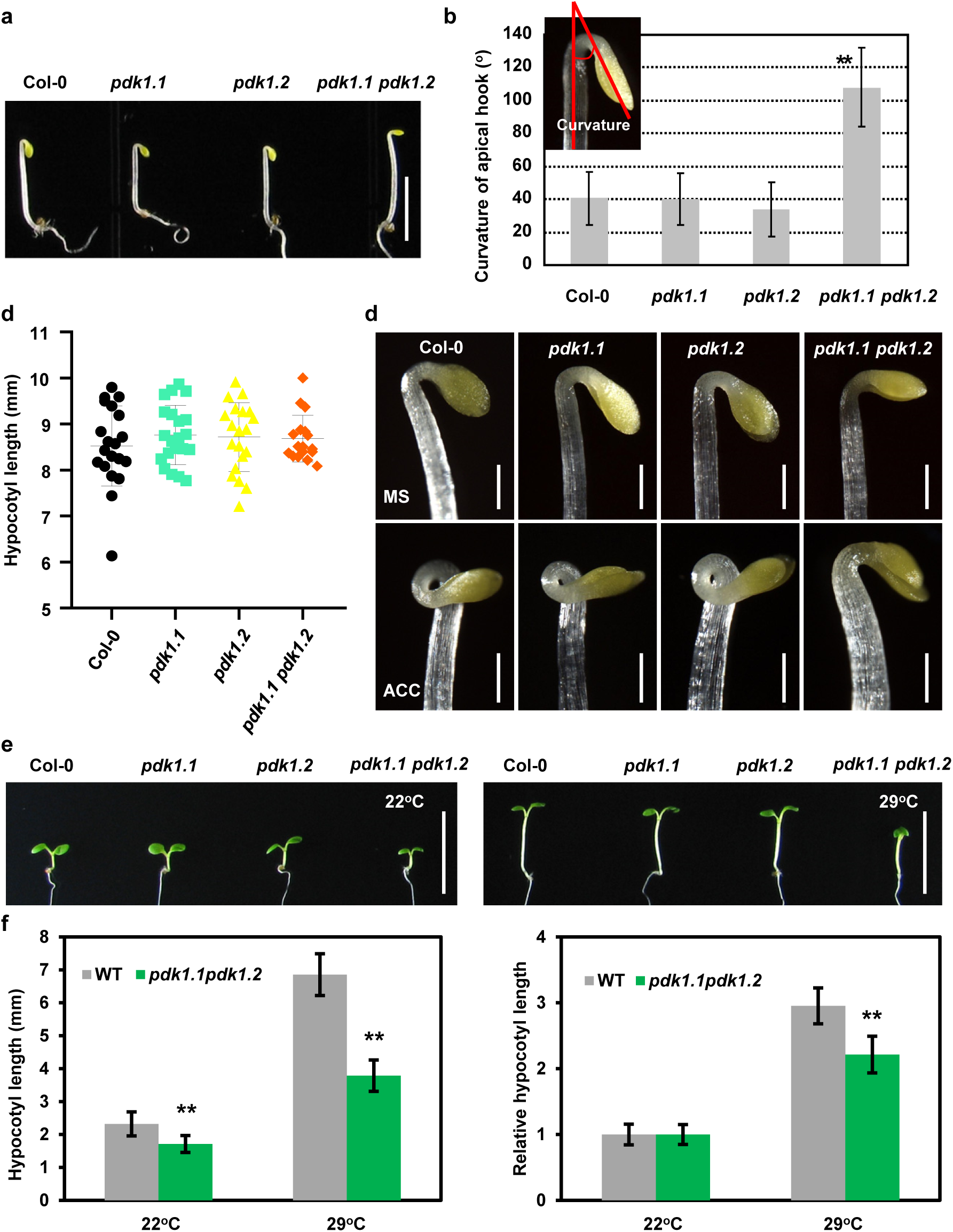
Deficiency of *PDK1.1* and *PDK1.2* impaired the normal development of the apical hook and high temperature-induced hypocotyl elongation. a-b. Observation (a, scale bar, 5 mm) and quantification (b) showed that etiolated seedlings of *pdk1.1 pdk1.2* exhibited less tight apical hooks. Col-0, *pdk1.1*, *pdk1.2,* and *pdk1.1 pdk1.2* seedlings were grown under dark for 90 h. Angles were measured by Image J. **, p<0.01, Student’s t-test. c. Etiolated seedlings of *pdk1.1 pdk1.2* exhibited comparably long hypocotyls to Col-0. Seedlings were grown under dark for 90 h. No significant difference was revealed by detected a Student’s t-test. d. Etiolated seedlings of *pdk1.1 pdk1.2* did not form apical hooks in the presence of ACC. Seedlings of Col-0, *pdk1.1*, *pdk1.2*, and *pdk1.1 pdk1.2* were grown under dark for 90 h in the absence or presence of ACC (10 µM). Scale bars, 500 µm. e-f. *pdk1.1 pdk1.2* showed defects in high temperature-induced hypocotyl elongation. Seedlings of Col-0, *pdk1.1*, *pdk1.2* and *pdk1.1 pdk1.2* were grown under light at 22°C or 29°C for 5 days, and hypocotyl elongation was observed (e, scale bar, 1 cm) and quantified (e). Hypocotyl length was measured by the Image J program and shown as mean ± s.d. (left, n = 20) or relative length by setting the hypocotyl length of Col-0 and *pdk1.1 pdk1.2* at 22°C as “1”, respectively (right, n = 20). **, p<0.01, Student’s t-test.

**Supplementary Figure 7.**
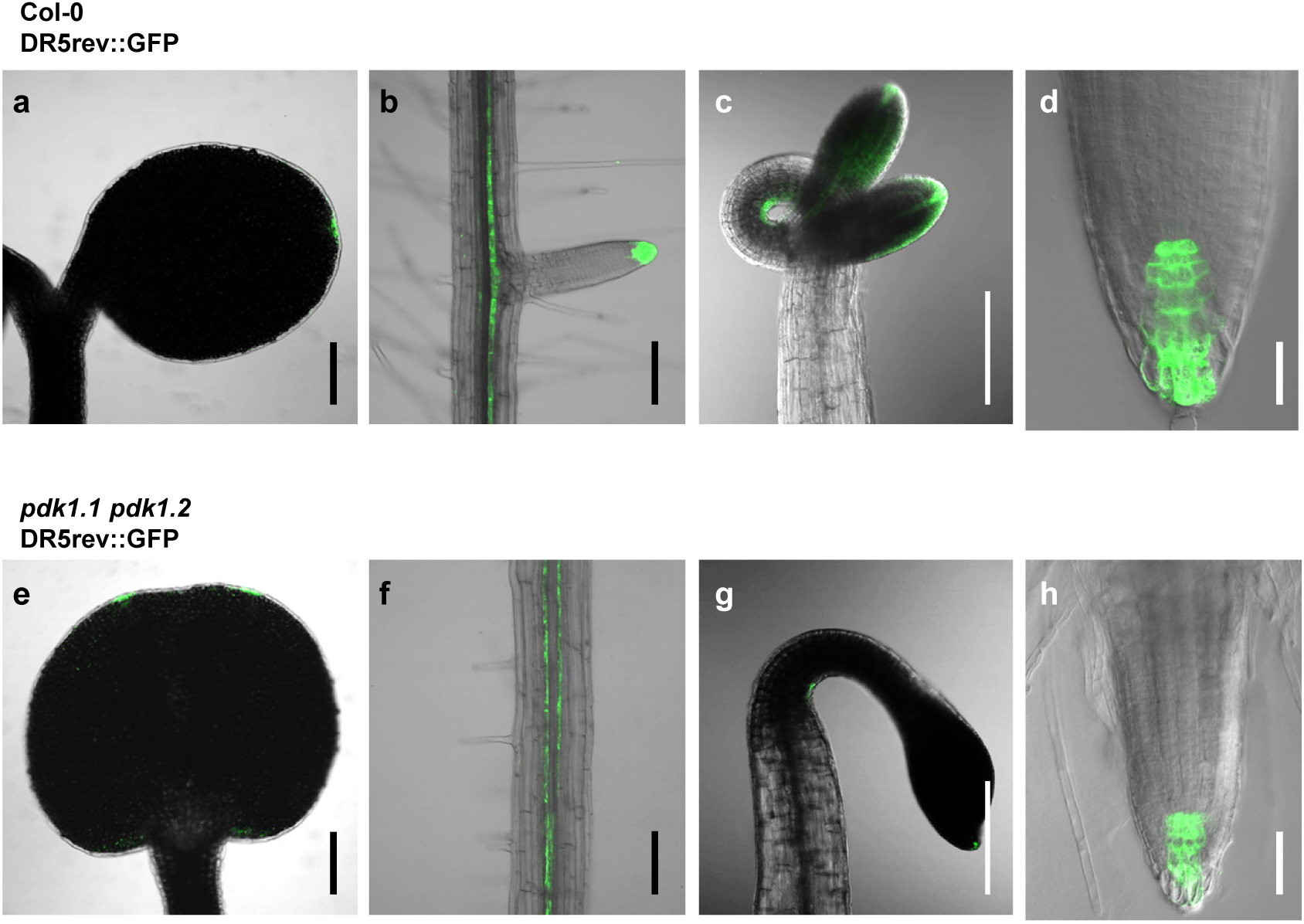
Loss of function of *PDK1.1* and *PDK1.2* impaired auxin distribution. Observation of the auxin responsive reporter DR5rev::GFP by CLSM indicated a dramatic decrease of the auxin maxima in *pdk1.1 pdk1.2* (e-h) compared with Col-0 (a-d). Fused cotyledon exhibiting two sites of auxin maxima in light-grown 7-day-old seedlings of *pdk1.1 pdk1.2* (e) compared with one of Col-0 (a); roots of light-grown 10-day-old seedlings (b and f); apical hooks of 10 µM ACC treated 4-day-old etiolated seedlings (c and g); roots of 10 µM ACC treated 4-day-old etiolated seedlings (d and h). Scale bars, 200 µm.

**Supplementary Figure 8.**
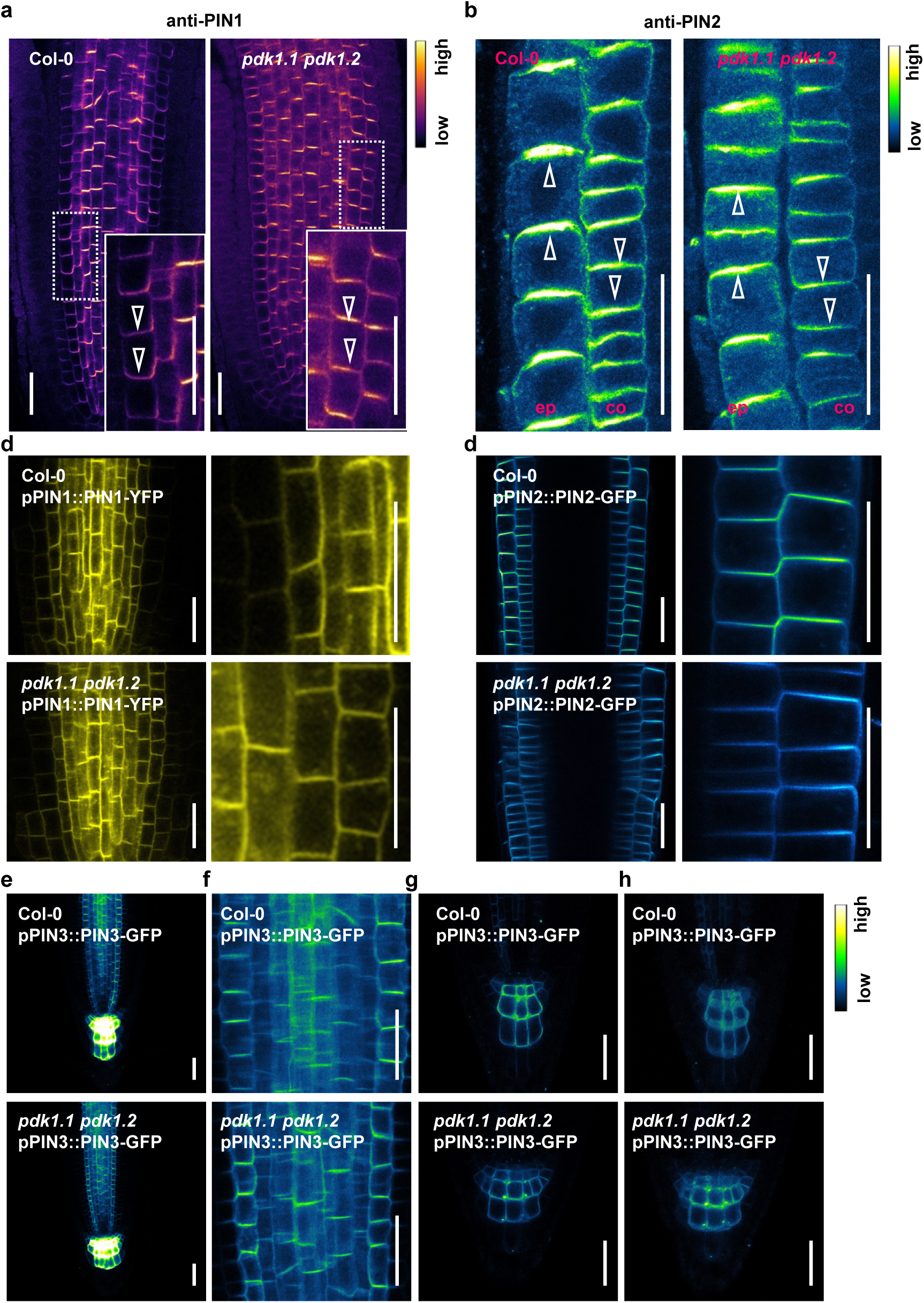
Deficiency of *PDK1.1* and *PDK1.2* did not affect the polarity of PIN proteins. a. Deficiency of *PDK1.1* and *PDK1.2* did not change the polarity of PIN1 in roots. Four-day-old seedlings of Col-0 and *pdk1.1 pdk1.2* were used for immunofluorescence with a rabbit anti-PIN1 antibody (1: 500), and then imaged by CLSM. The “mpl-inferno” LUT was used for photo visualization based on fluorescence intensity by the FIJI program. Arrowheads indicated basal localization. Scale bars, 20 µm. b. Deficiency of *PDK1.1* and *PDK1.2* did not change the polarity of PIN2 in roots. Four-day-old seedlings of Col-0 and *pdk1.1 pdk1.2* were used for immunofluorescence with a rabbit anti-PIN2 antibody (1: 500), and then imaged by CLSM. The “Green Fire Blue” LUT was used for photo visualization based on fluorescence intensity by the FIJI program. Arrowheads indicated apical localization in epidermis (“ep”) and the basal localization in cortex (“co”) respectively. Scale bars, 20 µm. c. Deficiency of *PDK1.1* and *PDK1.2* did not change the polarity of PIN1-YFP. Four-day-old seedlings of *pPIN1::PIN1-YFP* in Col-0 and *pPIN1::PIN1-YF*P in *pdk1.1 pdk1.2* were imaged by CLSM. Scale bars, 20 µm. d. Deficiency of *PDK1.1* and *PDK1.2* did not change the polarity of PIN2-GFP. Four-day-old seedlings of *pPIN2::PIN2-GFP* in Col-0 and *pPIN2::PIN2-GFP* in *pdk1.1 pdk1.2* were imaged by CLSM. Scale bars, 20 µm. e-h. Deficiency of *PDK1.1* and *PDK1.2* did not change the polarity of PIN3-GFP. Four-day-old seedlings of *pPIN3::PIN3-GFP* in Col-0 and *pPIN3::PIN3-GFP* in *pdk1.1 pdk1.2* were imaged by CLSM. (e) subcellular localization of PIN3-GFP; (f) an amplified view of PIN3-GFP in the root stele; (g) a close view of PIN3-GFP in root columella cells; (h) a 3D-projection of PIN3-GFP localization in root columella cells. The “Green Fire Blue” LUT was used for photo visualization based on fluorescence intensity by the FIJI program. Scale bars, 20 µm.

**Supplementary Figure 9.**
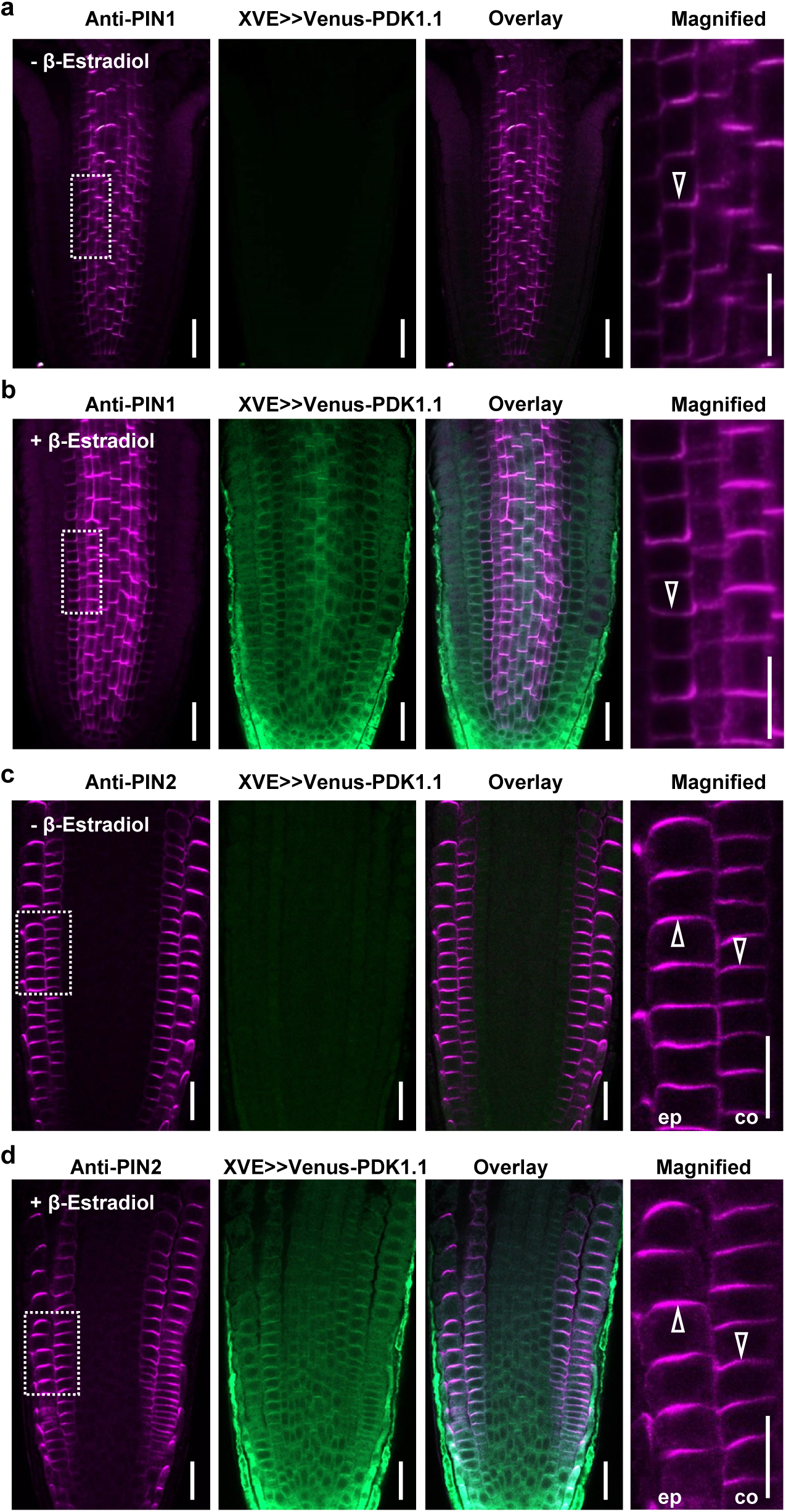
Overexpression of *PDK1.1* did not affect the polarity of PIN1. a-b. Induced overexpression of *PDK1.1* by β-Estradiol did not change the polar localization of PIN1. Four-day-old *XVE>>Venus-PDK1* (in Col-0) seedlings were transferred to MS plates without (a) or with (b) 5 µM β-Estradiol for 12 h, then used for immunofluorescence with a rabbit anti-PIN1 antibody (1: 500) and imaged by CLSM. Open arrowheads indicated the basal localization of PIN1. Scale bar, 20 µm. c-d. Induced overexpression of *PDK1.1* did not change the polar localization of PIN2. Same set of samples as in (a-b) were conducted for immunofluorescence with a rabbit anti-PIN2 antibody (1: 500) and imaged by CLSM. Open arrowheads indicated the apical or the basal localization of PIN2 in epidermis (short as “ep”) or cortex (“co”) respectively. Scale bar, 20 µm.

**Supplementary Figure 10.**
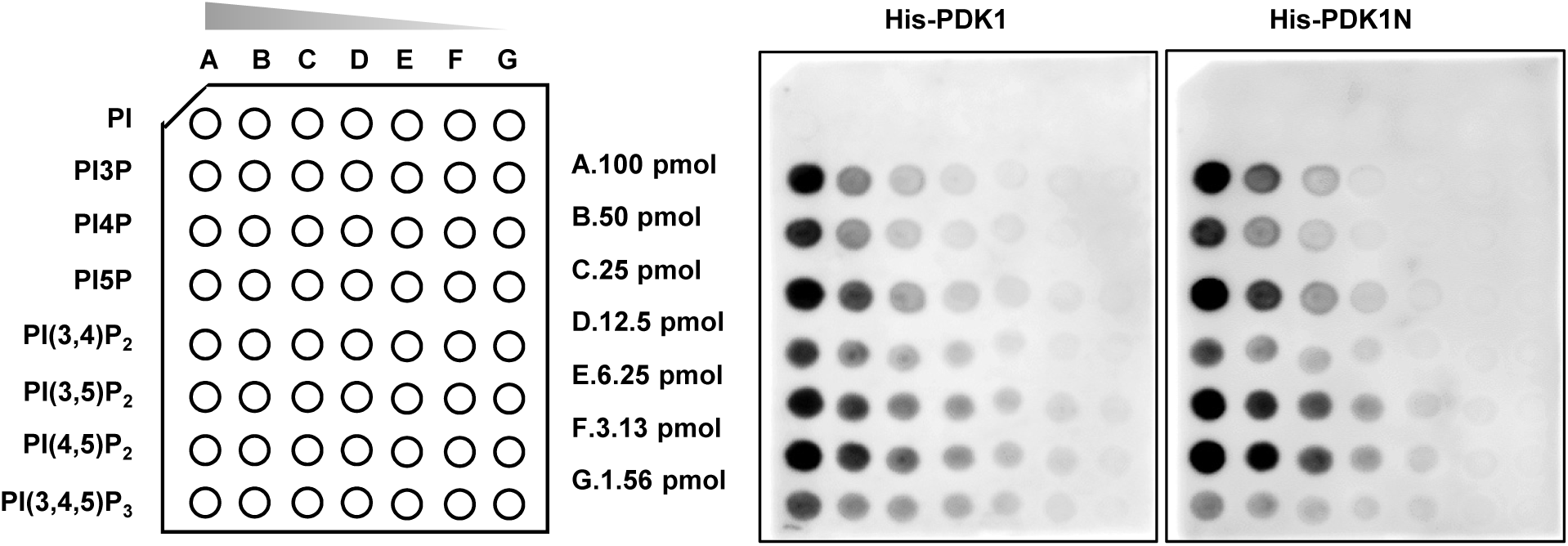
His-PDK1.1 and His-PDK1N bound to a similar spectrum of phospholipids. Lipid-protein blot overlay assays with PIP arrays revealed that recombinant His-PDK1.1 and His-PDK1.1N bound to similar spectrum of phospholipids. PIP arrays (from left to right: 100 pmol, 50 pmol, 25 pmol, 12.5 pmol, 6.25 pmol, 3.13 pmol, and 1.56 pmol, respectively) were incubated with His-tagged recombinant PDK1.1 or PDK1.1N proteins, and detected by a His antibody.

**Supplementary Figure 11.**
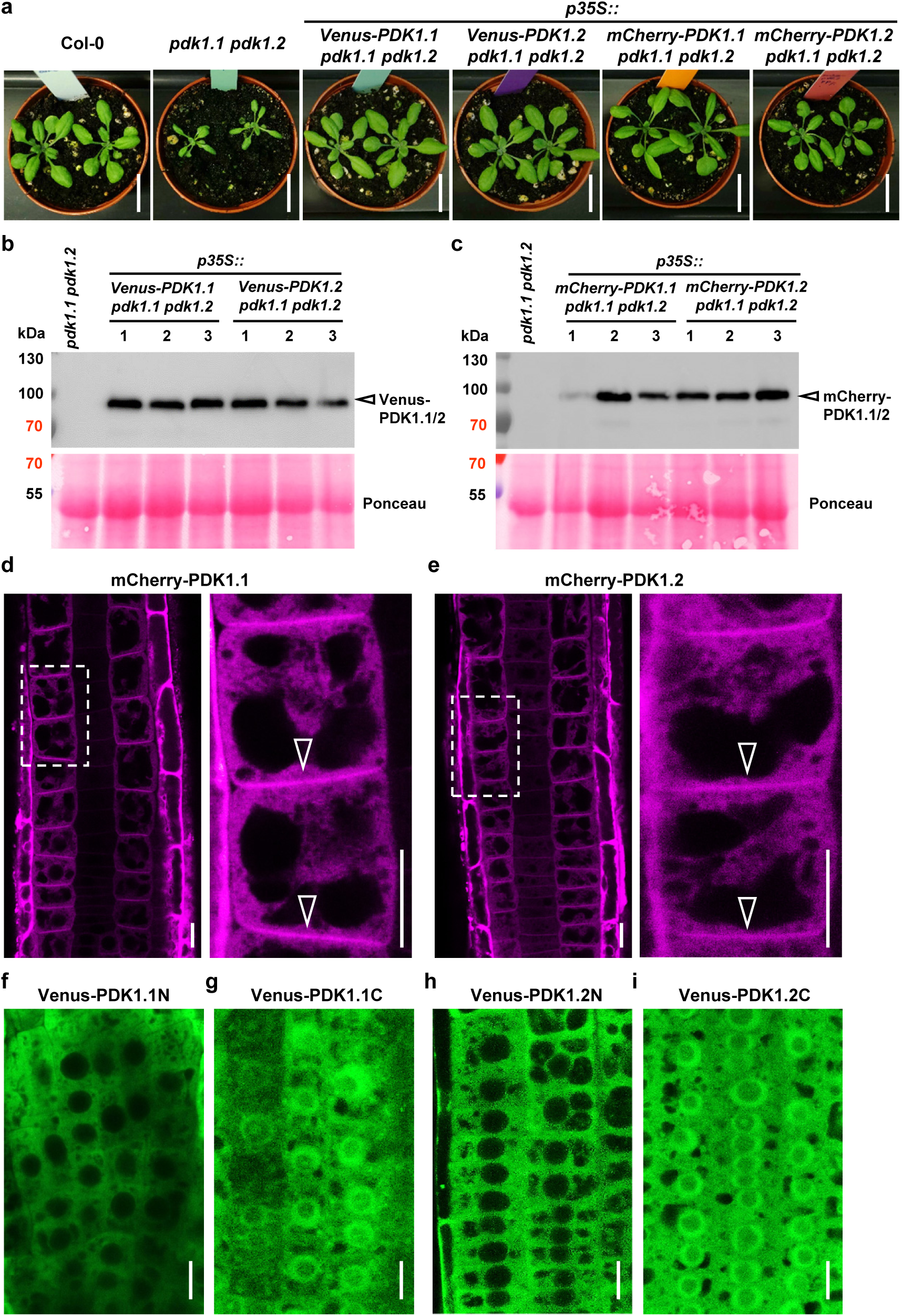
Analysis of *PDK1* transgenic lines. a. Expression of *PDK1.1* or *PDK1.2* (*p35S::Venus-PDK1.1, p35S::Venus-PDK1.2, p35S::mCherry-PDK1.1* and *p35S:: mCherry-PDK1.2*) rescued the growth defects of *pdk1.1 pdk1.2*. Adult plants (25-day-old) were observed and representative photos were shown. Scale bar, 2 cm. b. Western blot analysis verified the PDK1.1 or PDK1.2 expression (*35S::Venus-PDK1.1* and *35S::Venus-PDK1.2*) in *pdk1.1 pdk1.2*, respectively. Seven-day-old T_3_ homozygous seedlings were used for protein extraction and subjected to analysis. Upper panel, anti-GFP (1:2000); lower panel, Ponceau staining. c. Western blot verified the PDK1.1 or PDK1.2 expression (*35S::mCherry-PDK1.1* and *35S::mCherry-PDK1.2*) in *pdk1.1 pdk1.2*, respectively. Seven-day-old T_3_ homozygous seedlings were used for protein extraction and subjected to analysis. Upper panel, anti-RFP (1:2000); lower panel, Ponceau staining. d-e. mCherry-fused PDK1.1 (d) and PDK1.2 (e) localized to both cytoplasm and the basal side of PM. Four-day-old seedlings of *p35S::mCherry-PDK1.1* and *p35S::mCherry-PDK1.2* were imaged by CLSM. Open arrowheads indicated the basal polar localization. Scale bar, 10 µm. f-i. Subcellular localization of Venus-fused PDK1.1N (f, cytoplasm), Venus fused PDK1.1C (g, both cytoplasm and nucleus), Venus fused PDK1.2N (h, cytoplasm), and Venus fused PDK1.2C (i, both cytoplasm and nucleus). Four-day-old seedlings expressing corresponding fusion proteins were imaged by CLSM. Scale bar, 10 µm.

**Supplementary Figure 12.**
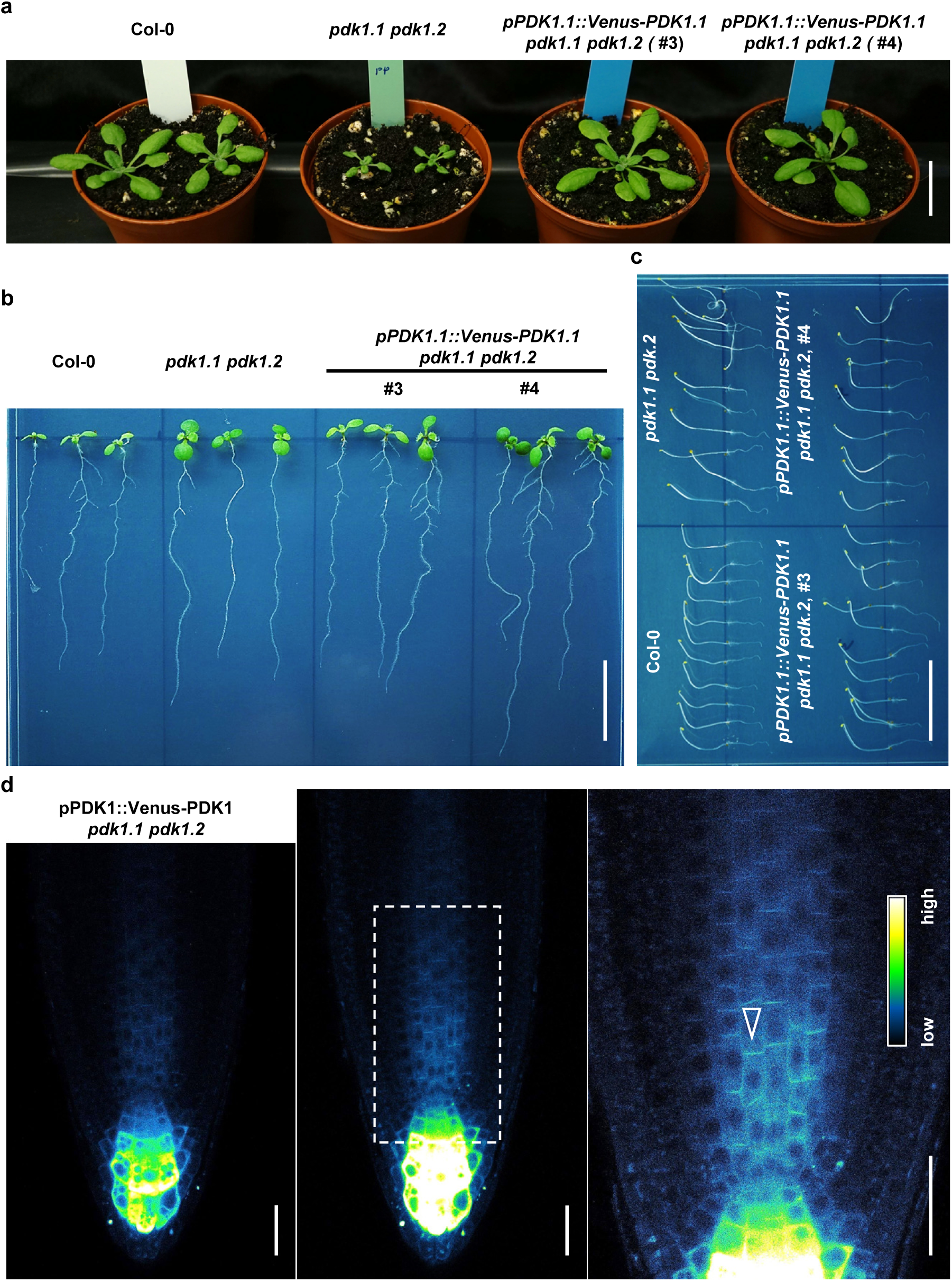
Functional *pPDK1.1::Venus-PDK1.1* localized at both cytoplasm and the basal side of PM. a. *pPDK1.1::Venus-PDK1.1* rescued the growth defects of *pdk1.1 pdk1.2*. 25-day-old adult plants were observed and representative photos are shown. Scale bar, 2 cm. b. *pPDK1.1::Venus-PDK1.1* rescued the lateral root defects of *pdk1.1 pdk1.2*. 10-day-old seedlings were observed and representative photos are shown. Scale bar, 2 cm. c. *pPDK1.1::Venus-PDK1.1* rescued the defects of *pdk1.1 pdk1.2* in the hypocotyl gravitropic response. 4-day-old etiolated seedlings were observed and representative photos are shown. Scale bar, 2 cm. d. Venus-fused PDK1.1 localized to both cytoplasm and PM, especially with a predominant presence at the basal side of PM. Four-day-old seedlings of *pPDK1.1::Venus-PDK1.1* were imaged by CLSM. Open arrowheads indicate the basal polar localization. The “Green Fire Blue” LUT was used for photo visualization based on fluorescence intensity by the FIJI program. Scale bar, 20 µm.

**Supplementary Figure 13.**
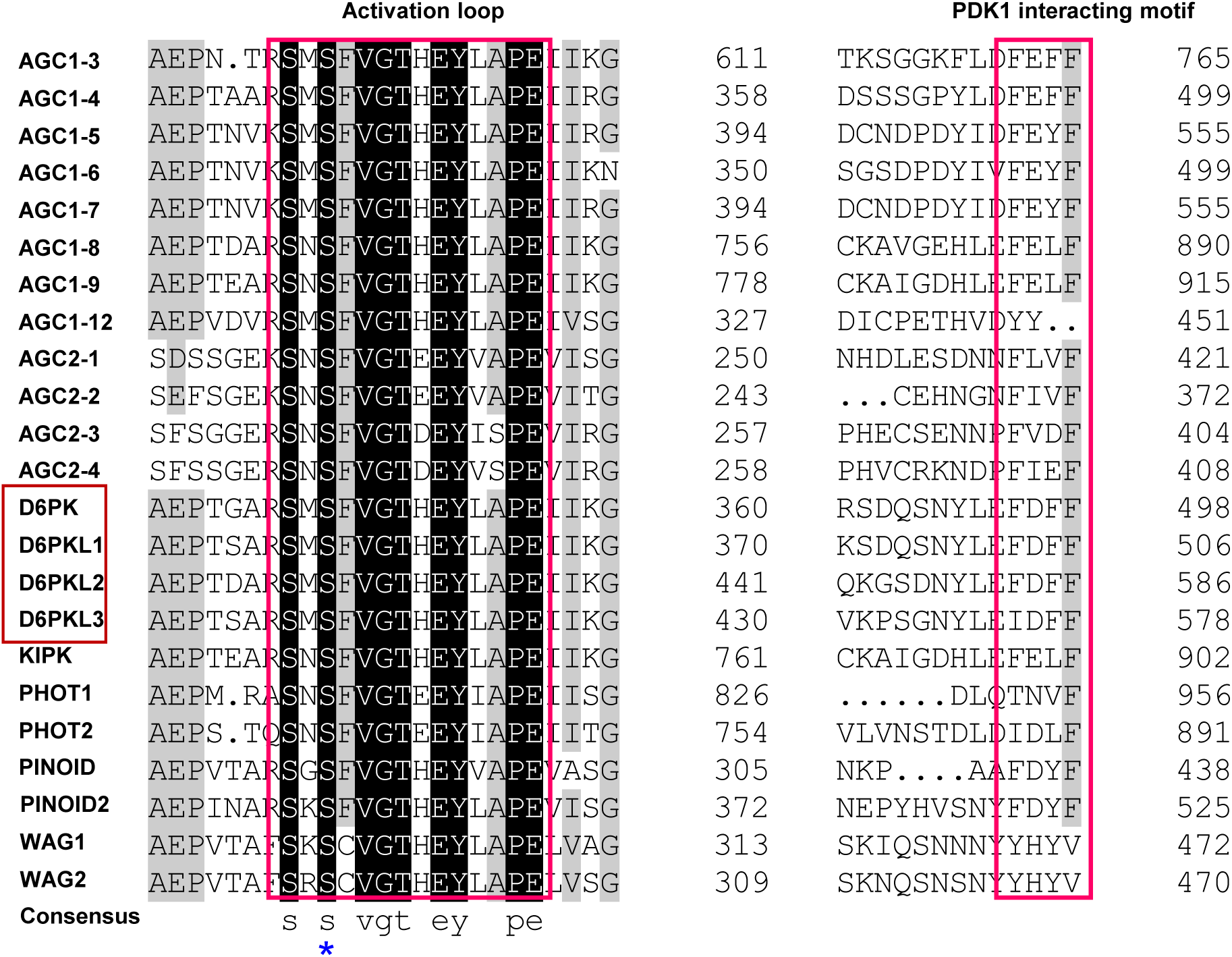
A conserved activation loop and a PDK1 interaction pocket (PIF) motif of AGC protein kinases. Alignment of AGC protein kinases was performed with the DNAMAN program, and the similarity was shown in different colours: back, 100% identity; grey, ≥75%. Asterisk indicated a conserved phosphosite (Ser345 in D6PK).

**Supplementary Figure 14.**
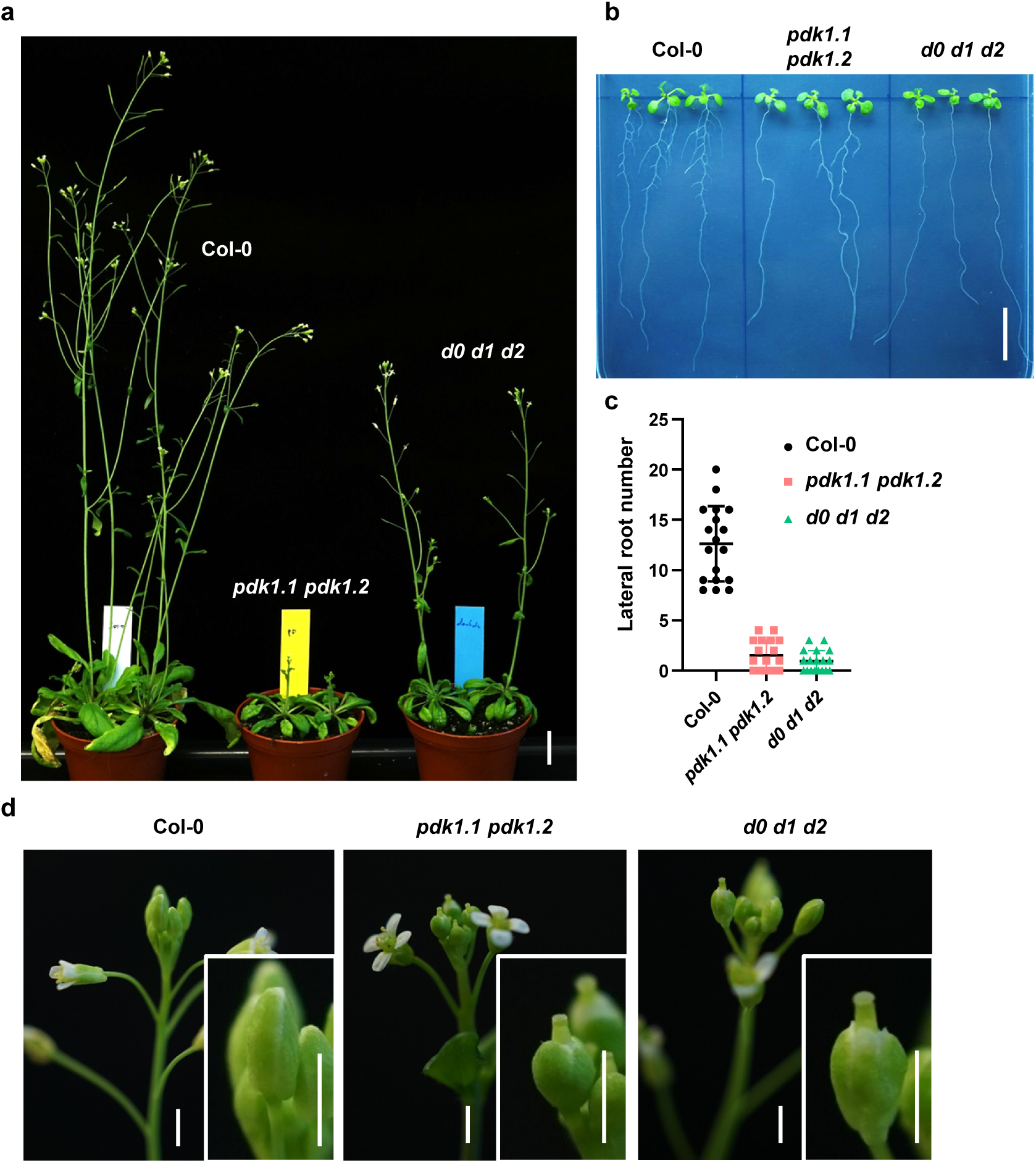
The *d0 d1 d2* triple mutant phenocopied *pdk1.1 pdk1.2*. a. Representative photos of 35-day-old Col-0, *d0 d1 d2* and *pdk1.1 pdk1.2* plants. The delayed flowering of *pdk1.1 pdk1.2* was not observed in *d0 d1 d2*. Scale bar, 2 cm. b-c. *d0 d1 d2* plants showed reduced lateral root number, similar to that of *pdk1.1 pdk1.2*. Twelve-day-old seedlings were observed (b, Scale bar, 1 cm) and number of lateral roots were measured (c). Data are shown as mean ± s.d. (n = 18). d. Observations showed similar abnormal flowers of *d0 d1 d2* and *pdk1.1 pdk1.2*, with the stigma protruding out of the flower buds before opening. Scale bar, 2 mm.

**Supplementary Figure 15.**
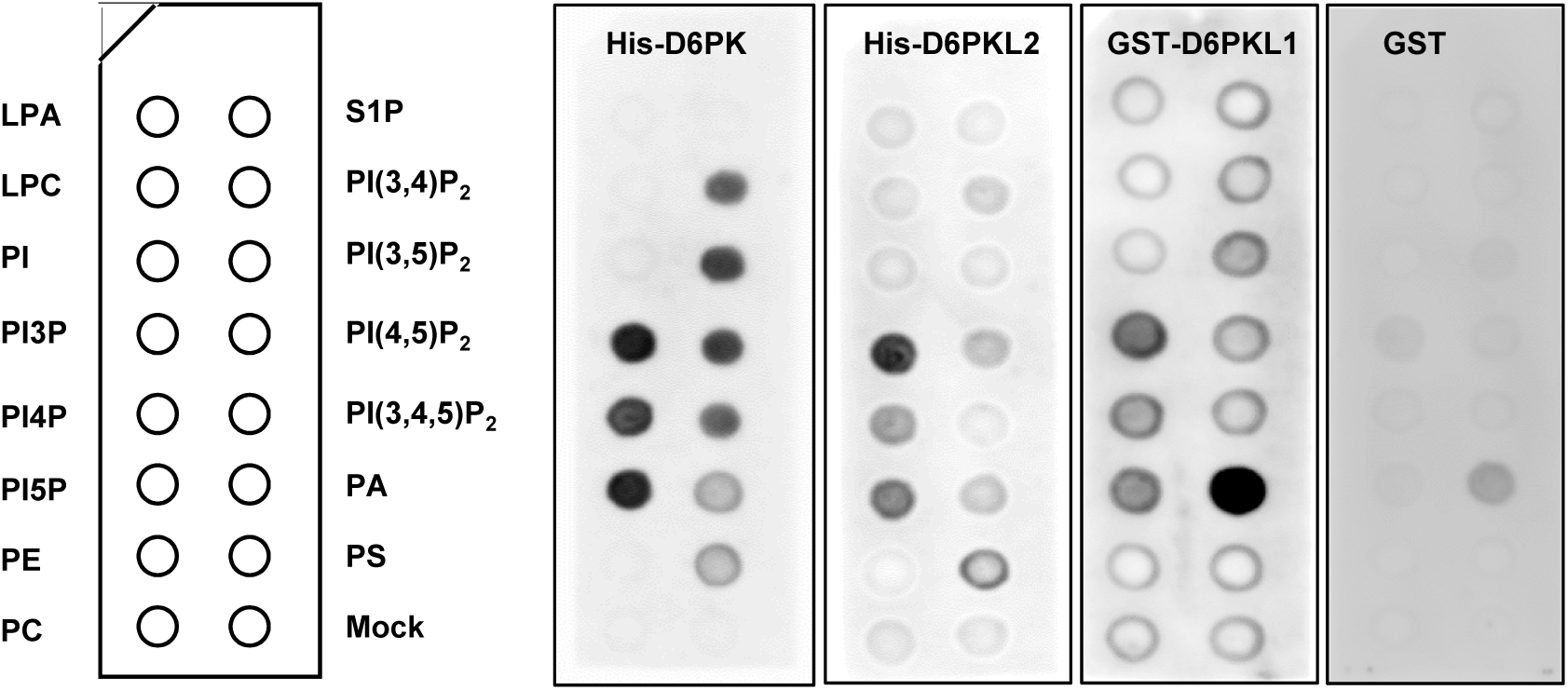
D6PK and D6PKL1/2 (D6PKs) exhibited a similar lipid binding preference as PDK1.1/2. Lipid-protein blot overlay assays revealed that D6PKs bound to phospholipids. PIP strips were incubated with recombinant His-D6PK, His-D6PKL2, GST-D6PKL1, and GST (as control) proteins, and detected by a GST or a His antibody accordingly.

**Supplementary Figure 16.**
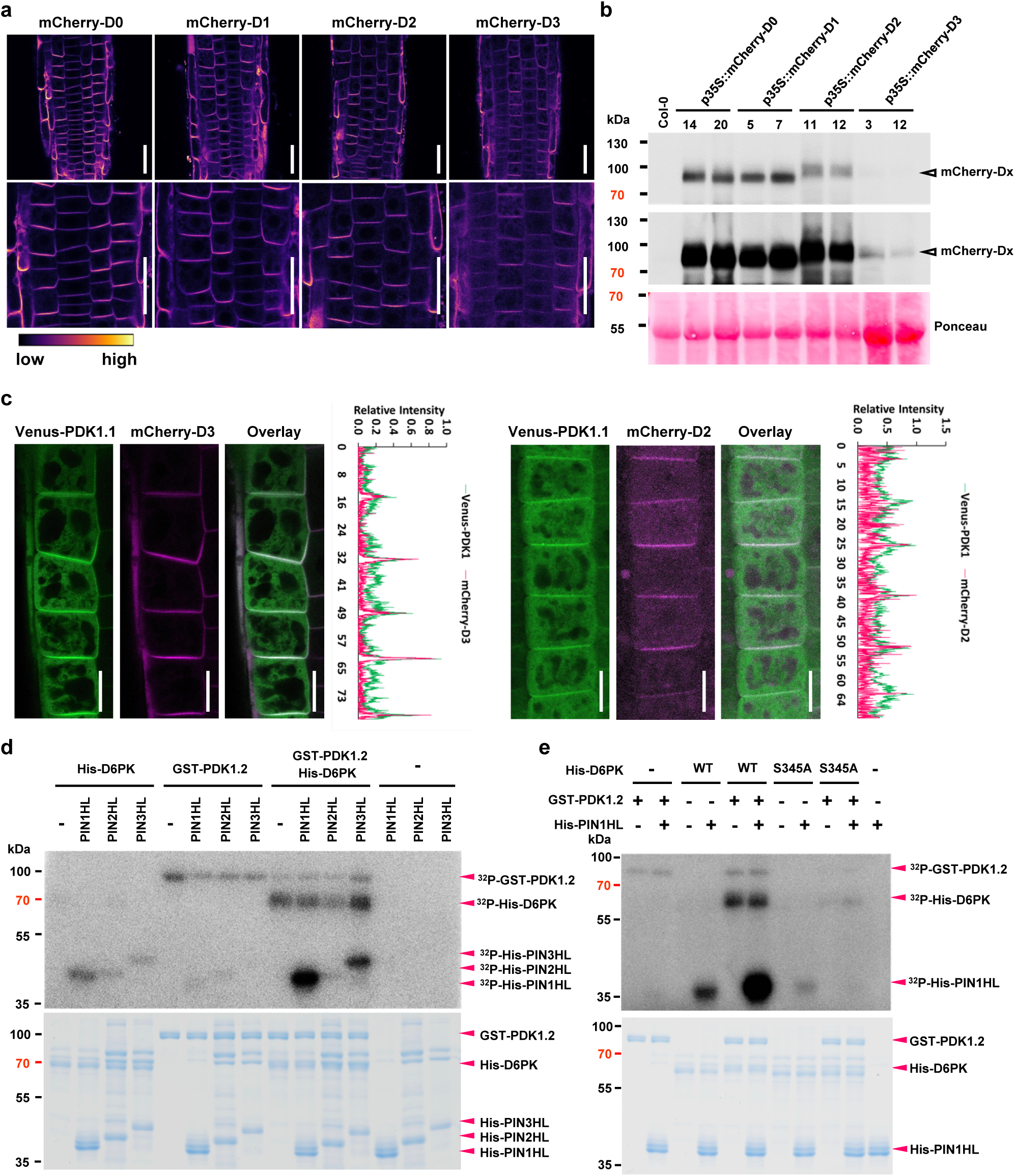
mCherry-D6PKs co-localized with Venus-PDK1.1 at the basal side of PM. a. mCherry-fused D6PKs localized to the basal side of PM. Four-day-old seedlings of *p35S::mCherry-D6PK/D6PKLs* (short as *D0 to D3*) were imaged by CLSM. The “mpl-inferno” LUT was used for photo visualization based on fluorescence intensity by the FIJI program. Scale bars, 10 µm. b. Western blot verified the mCherry-D0∼D3 protein level (*35S::mCherry-D0∼D3*) in Col-0, respectively. Seven-day-old T_3_ homozygous seedlings were used for protein extraction and subjected to Western blot analysis. Upper panel, anti-RFP (1:1000), short exposure (0.5 seconds); medium panel, anti-RFP (1:1000), long exposure (5 seconds, for low expression of mCherry-D3); lower panel, Ponceau staining. c. Venus-PDK1.1 co-localized with mCherry-D6PKL2 and mCherry-D6PKL3 at the basal side of PM. Four-day-old seedlings of *p35S::mCherry-D6PKL2 / p35S::Venus-PDK1.1* and *p35S::mCherry-D6PKL3 / p35S::Venus-PDK1.1* were imaged by CLSM. Scale bar, 10 µm. d. *In vitro* kinase assay with [^32^P]-ATP revealed that GST-PDK1.2-conducted phosphorylation of D6PK facilitates its activity towards PIN-HL phosphorylation. Upper panel, autoradiography; lower panel, CBB staining. e. *In vitro* kinase assay with [^32^P]-ATP revealed that GST-PDK1.2-conducted full phosphorylation and activation of D6PK, towards His-PIN1-HL phosphorylation, required the phosphorylation at S345 for D6PK. Upper panel, autoradiography of ^32^P; lower panel, CBB staining.

**Supplementary Figure 17.**
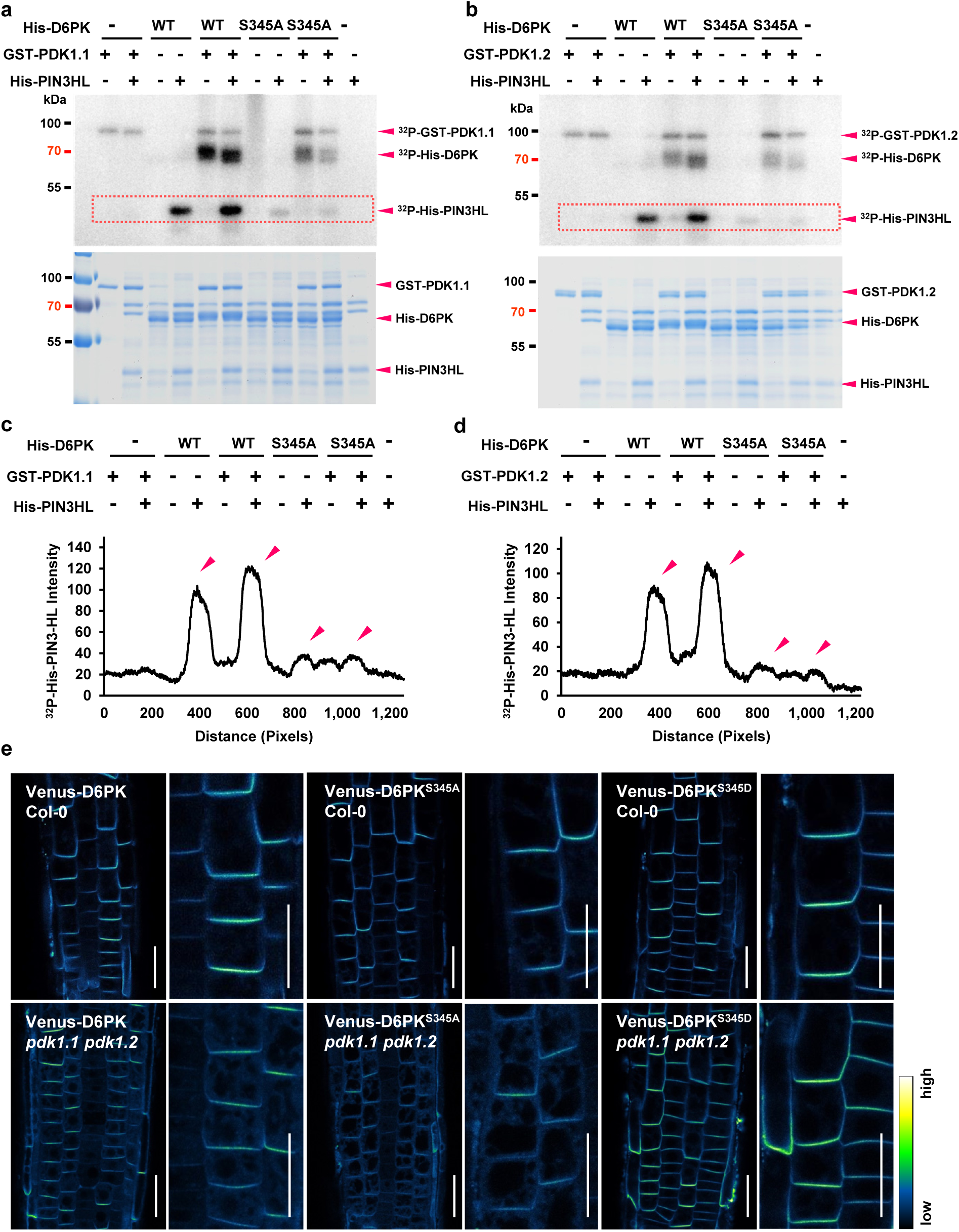
PDK1.1 activated D6PK through phosphorylating Ser345 (S345). a-b. *In vitro* kinase assay with [^32^P]-ATP revealed that GST-PDK1.1- (a) and GST-PDK1.2 (b)-conducted full phosphorylation and activation of D6PK, towards His-PIN3-HL phosphorylation, required the phosphorylation at S345 for D6PK. Upper panel, autoradiography of ^32^P; lower panel, CBB staining. c-d. Quantification of the band intensities of D6PK variants’ phosphorylation toward PIN3HL by the Image J program, as squared in Supplementary Fig. 17A or B. e. The subcellular localizations of Venus-D6PK, Venus-D6PK^S345A^, and Venus-D6PK^S345D^ do not show difference in the basal polarity. Four-day-old seedlings of *p35S::Venus-D6PK*, *p35S::Venus-D6PK^S345A^*, and *p35S::Venus-D6PK^S345D^*, in Col-0 and *pdk1.1 pdk1.2* respectively, were imaged by CLSM. The “Green Fire Blue” LUT was used for photo visualization based on fluorescence intensity by the FIJI program. Representative photos of at least three independent transgenic lines were shown. Scale bars, 10 µm.

**Supplementary Figure 18.**
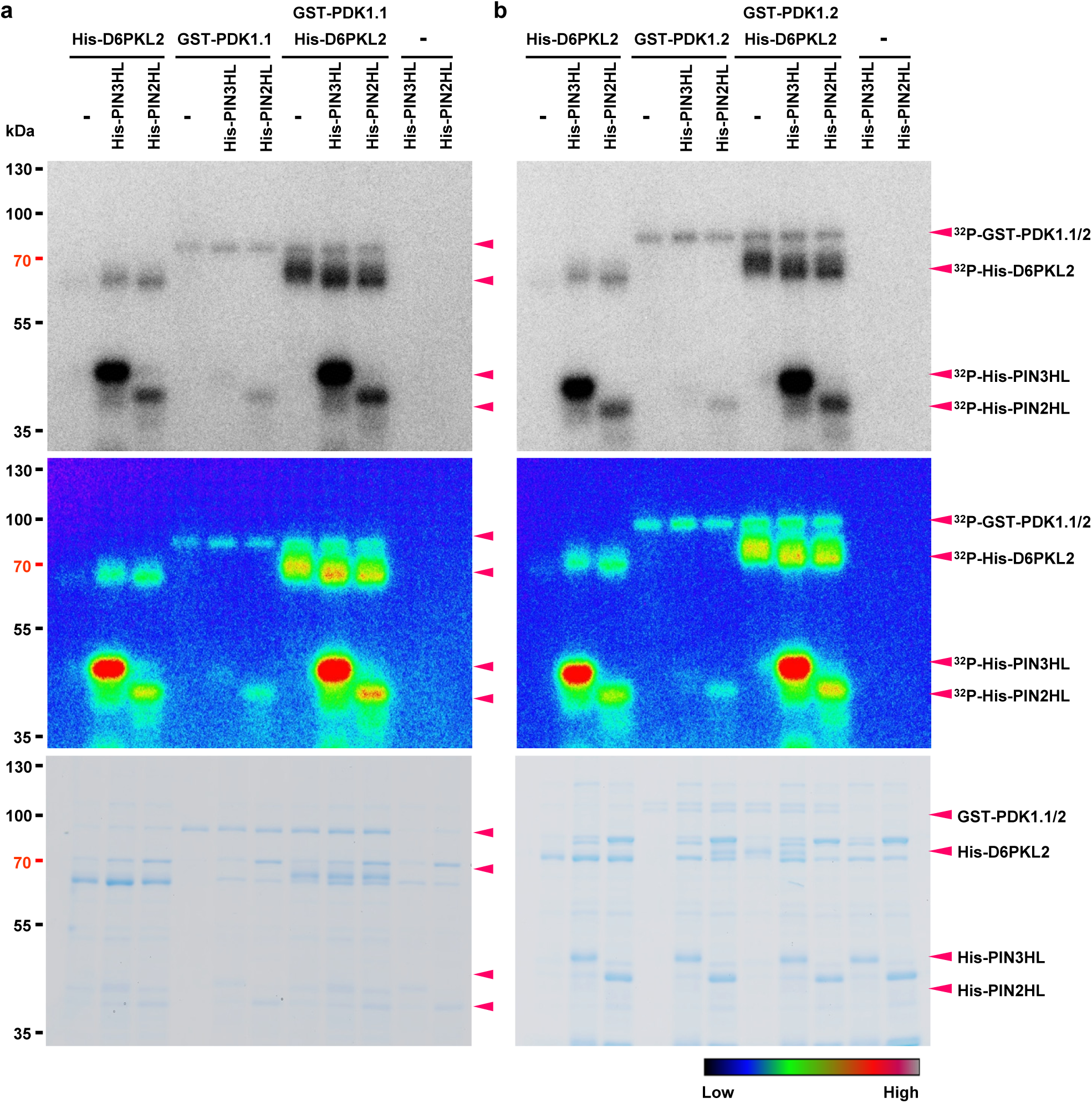
PDK1 phosphorylated D6PKL2 to increase the phosphorylation of PIN3/2-HLs. *In vitro* kinase assay with [^32^P]-ATP revealed that GST-PDK1.1 (a) and GST-PDK1.2 (b)-conducted phosphorylation of D6PK facilitated its activity towards PIN3/2-HL phosphorylation. Upper panels, autoradiography of ^32^P; Middle panels, the “Rainbow RGB” LUT was used for data visualization based on fluorescence intensity by the FIJI program; lower panels, CBB staining.

**Supplementary Table 1.**
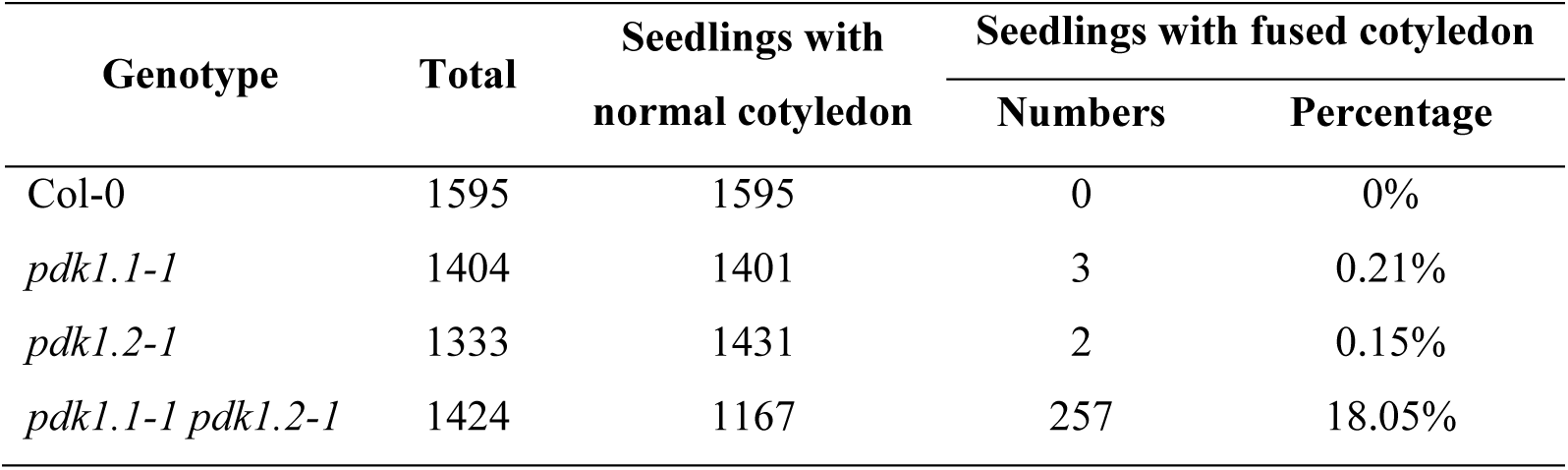
Loss-of-function of *PDK1.1* and *PDK1.2* leads to defective cotyledon development. Seven-day-old seedlings were observed and numbers of seedlings with defective cotyledon development were counted.

**Supplementary Table 2.**
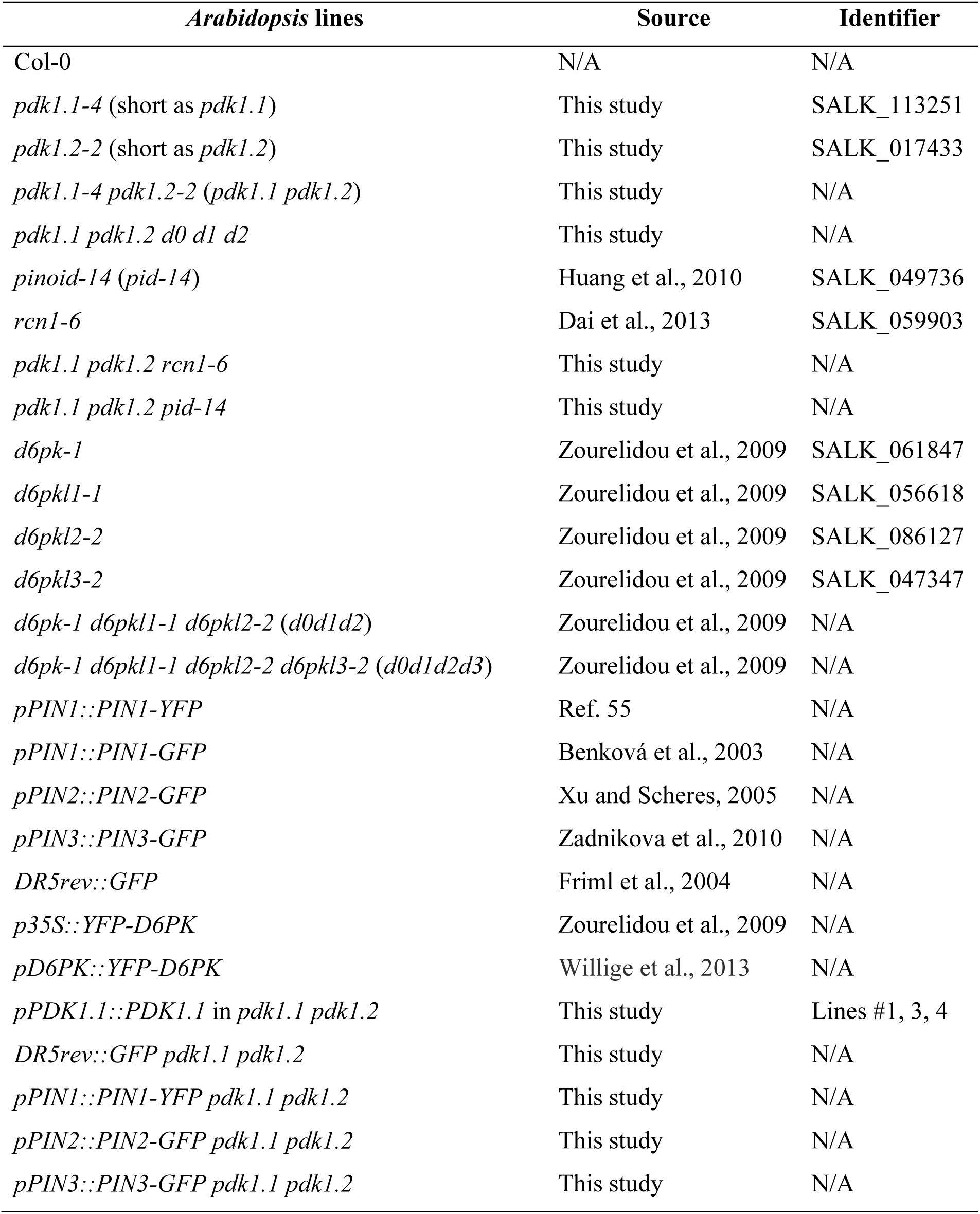
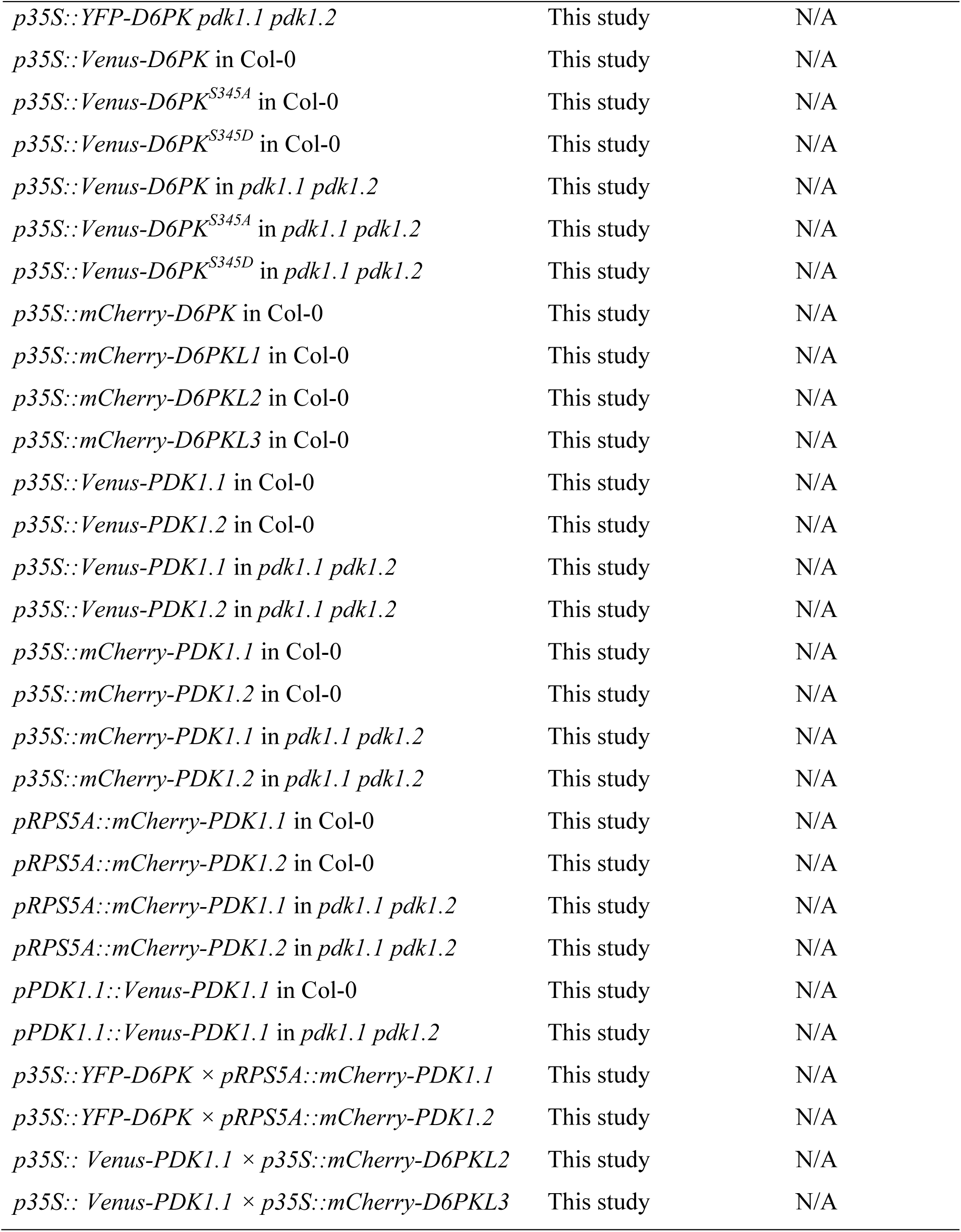
List of plant lines, including mutants and maker lines, used in this study.

**Supplementary Table 3.**
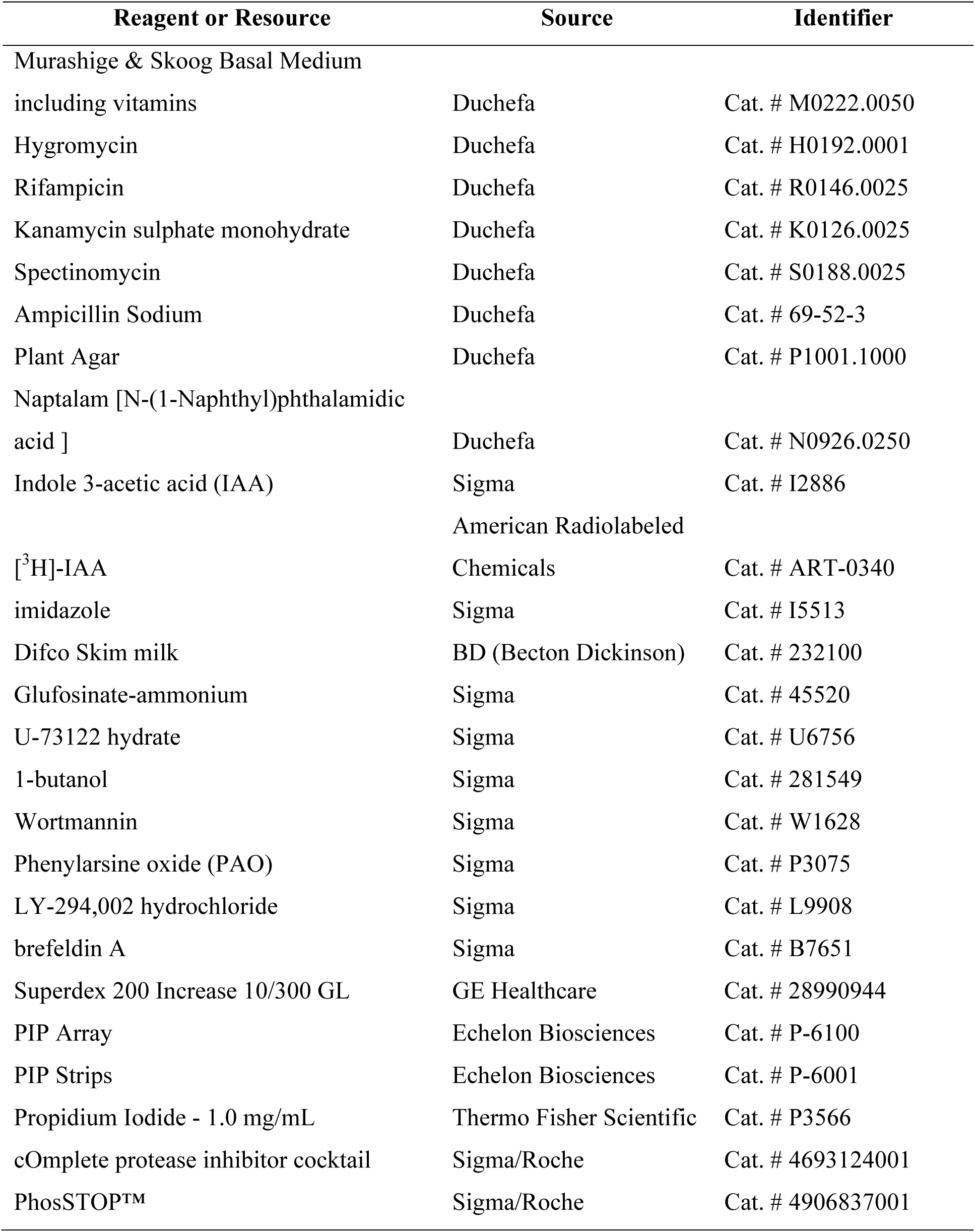
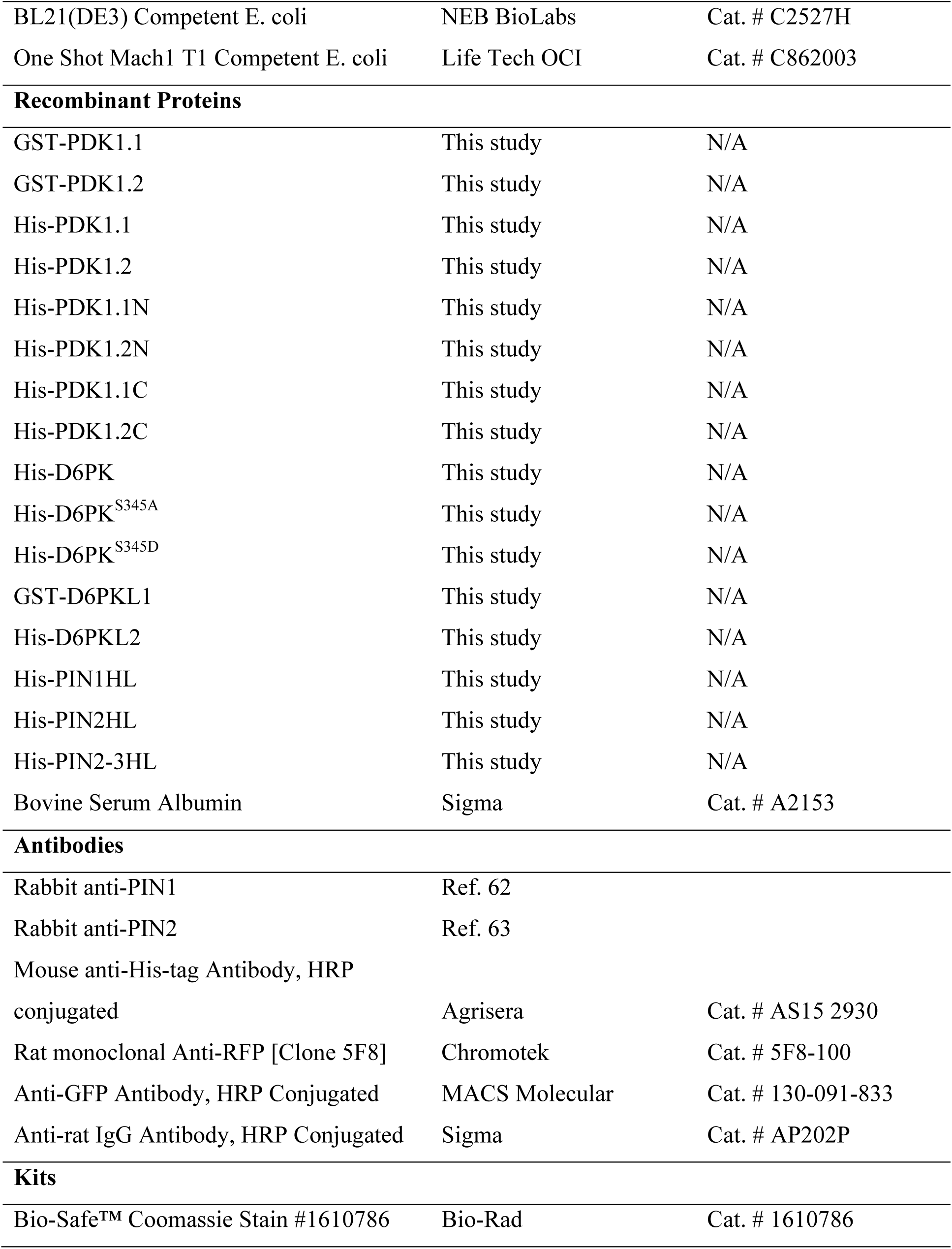
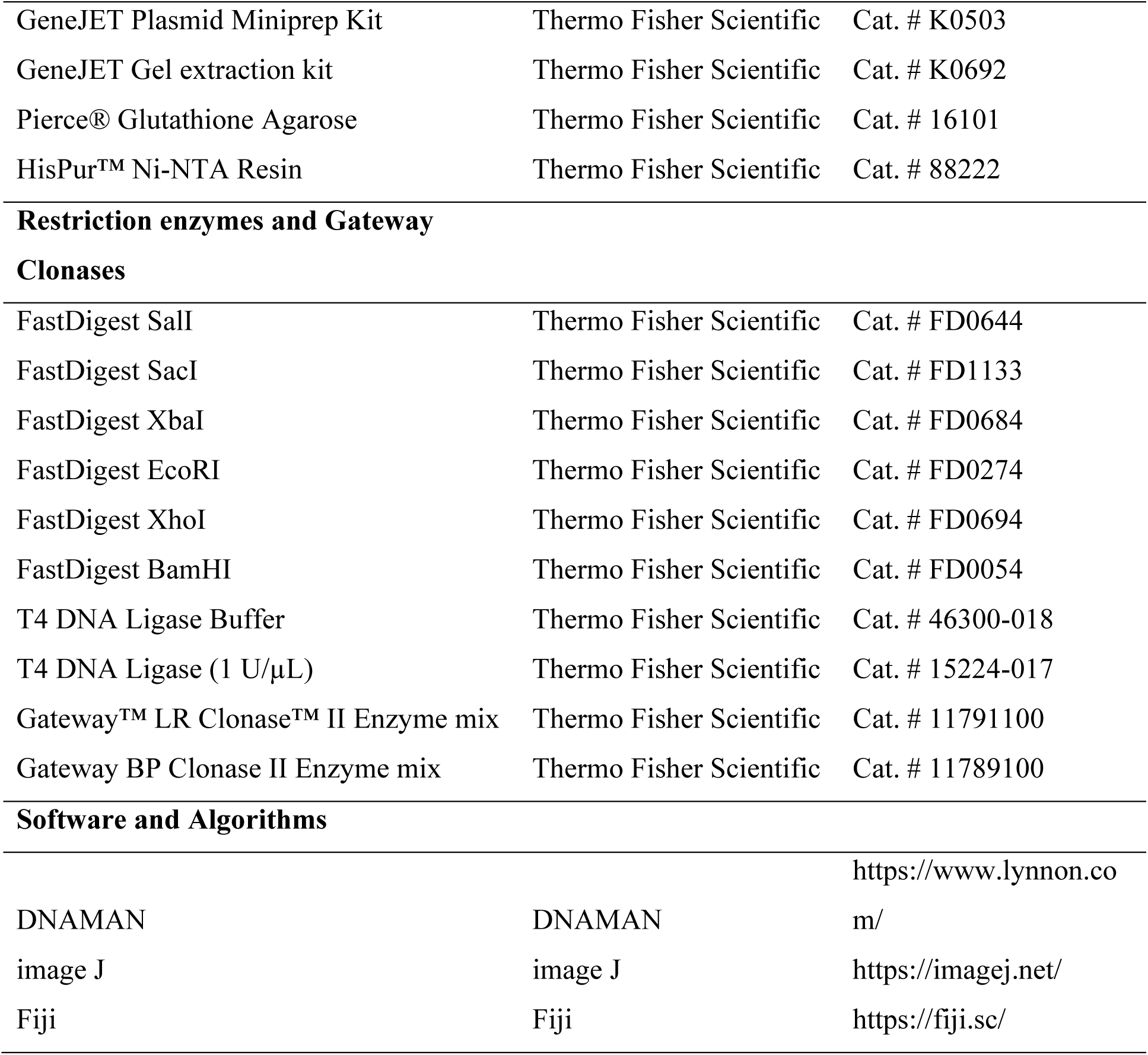
List of reagents used in this study.

**Supplementary Table 4.**
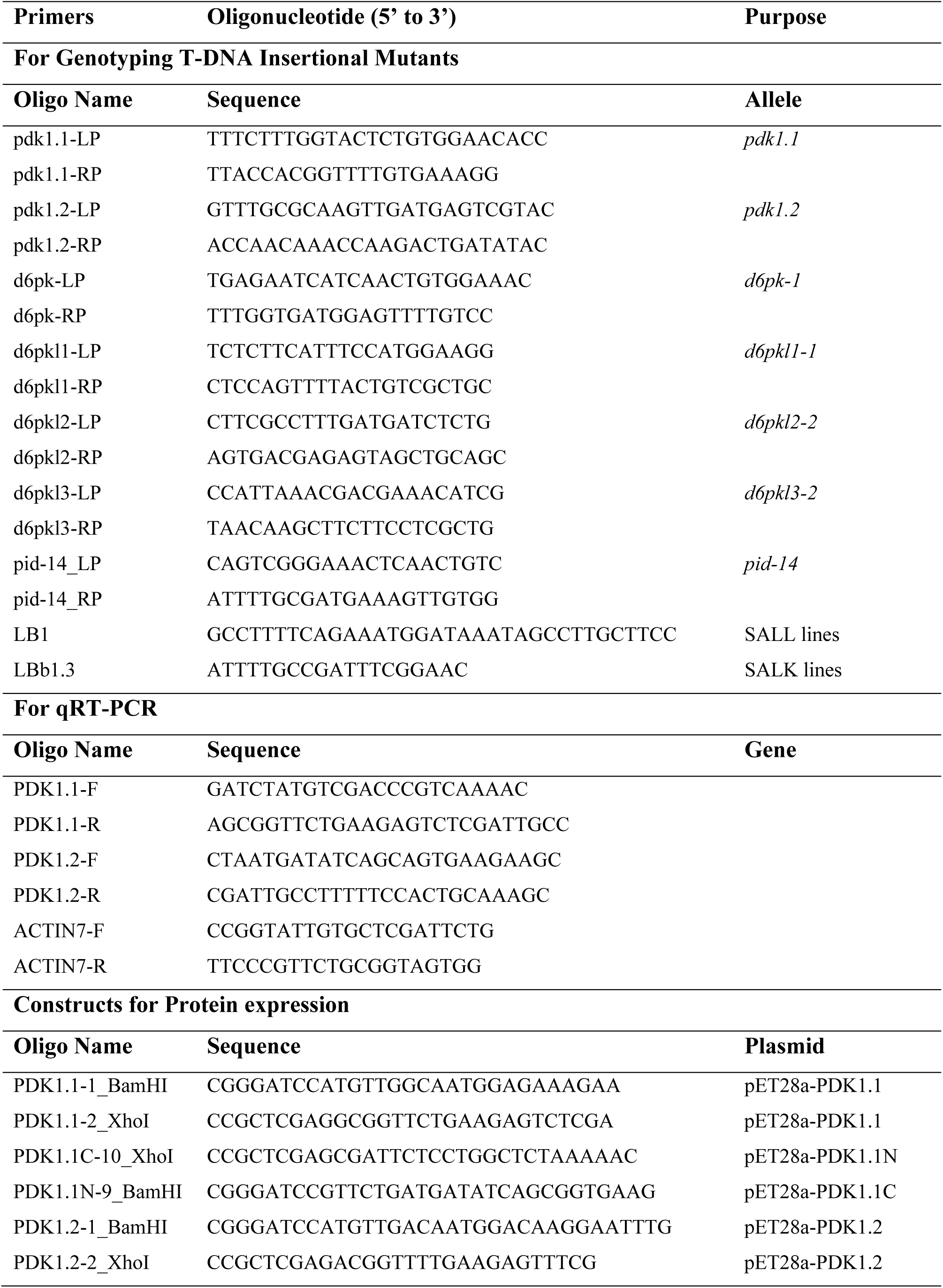
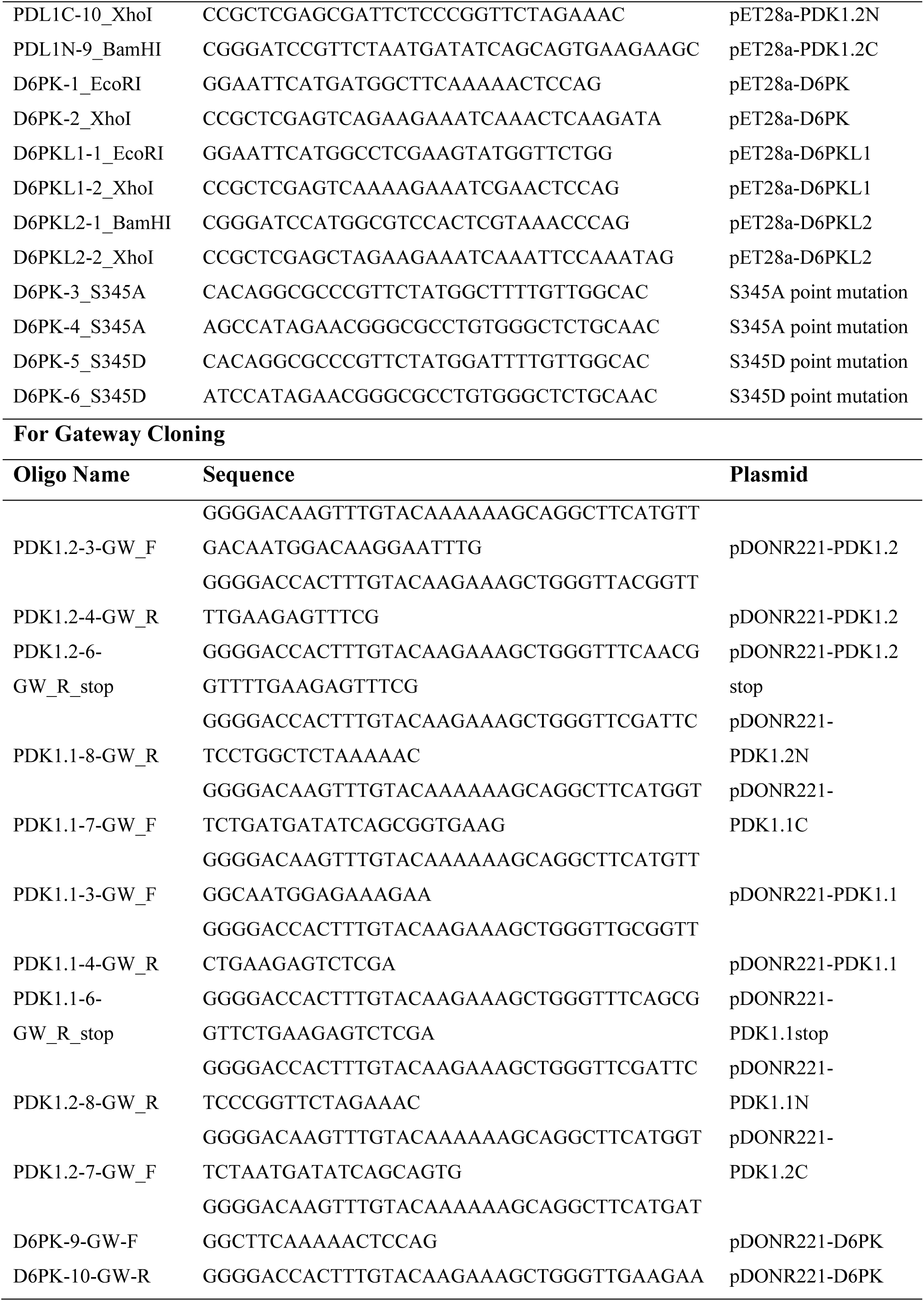
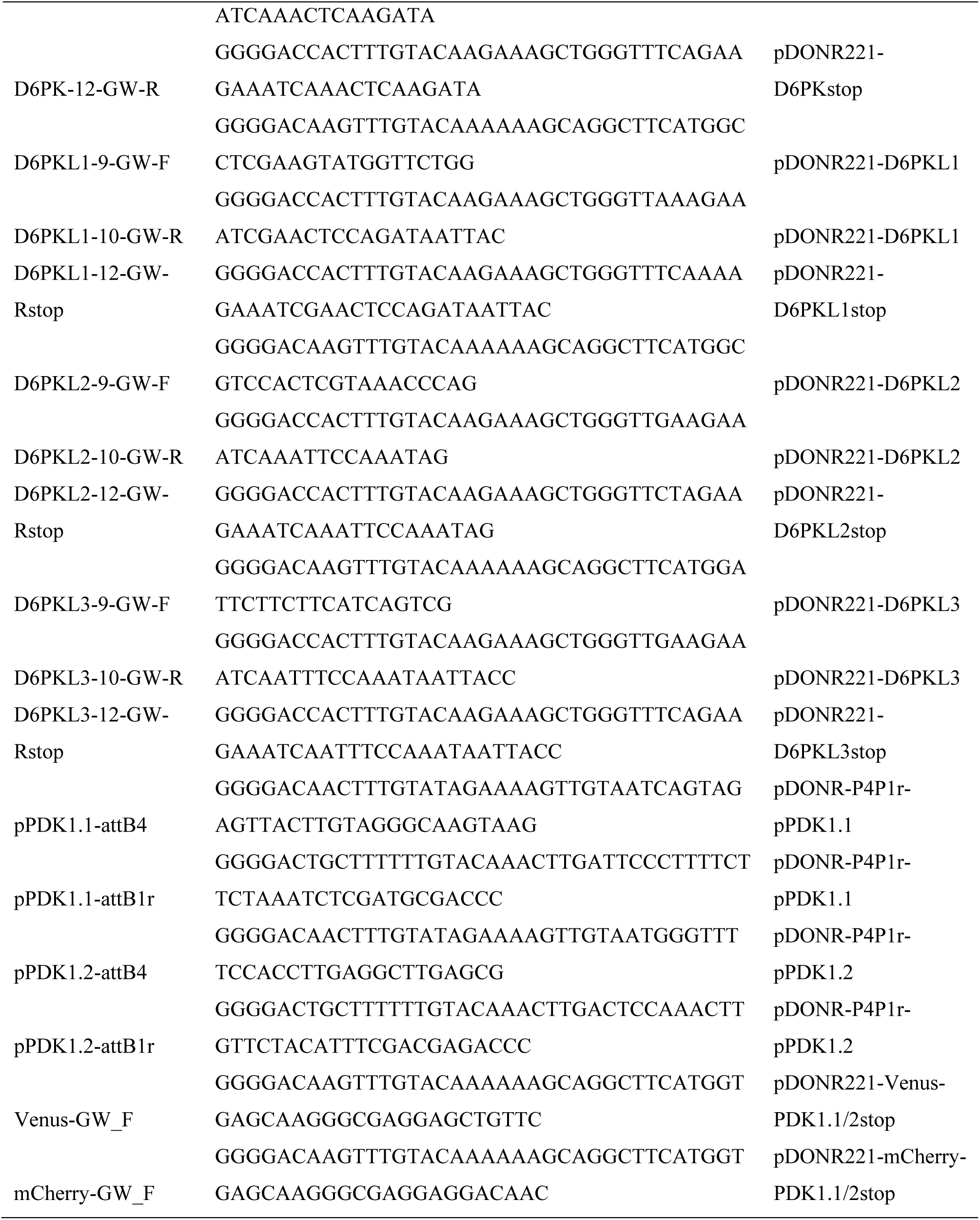
List of primers used in this study.

**Supplementary Table 5.**
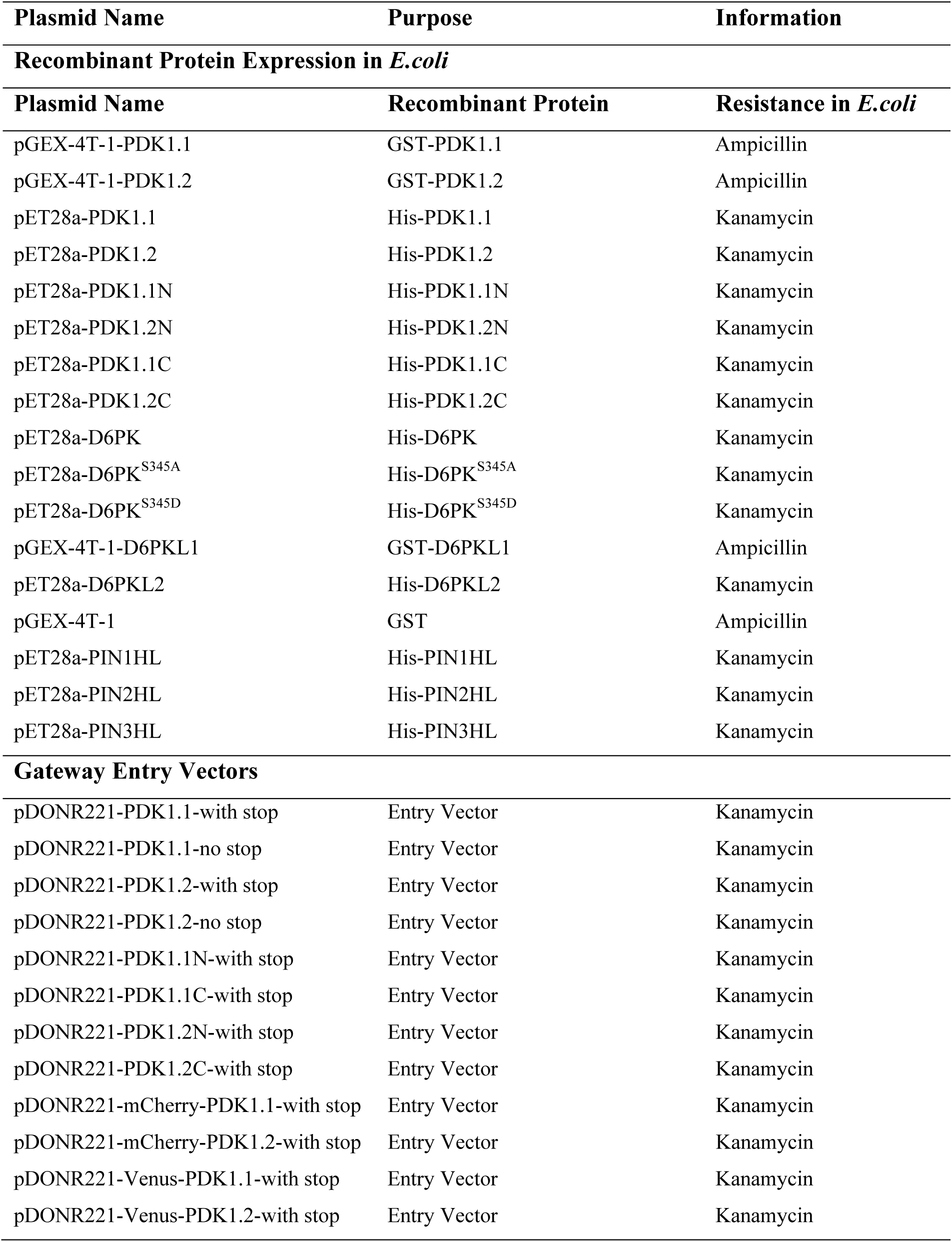
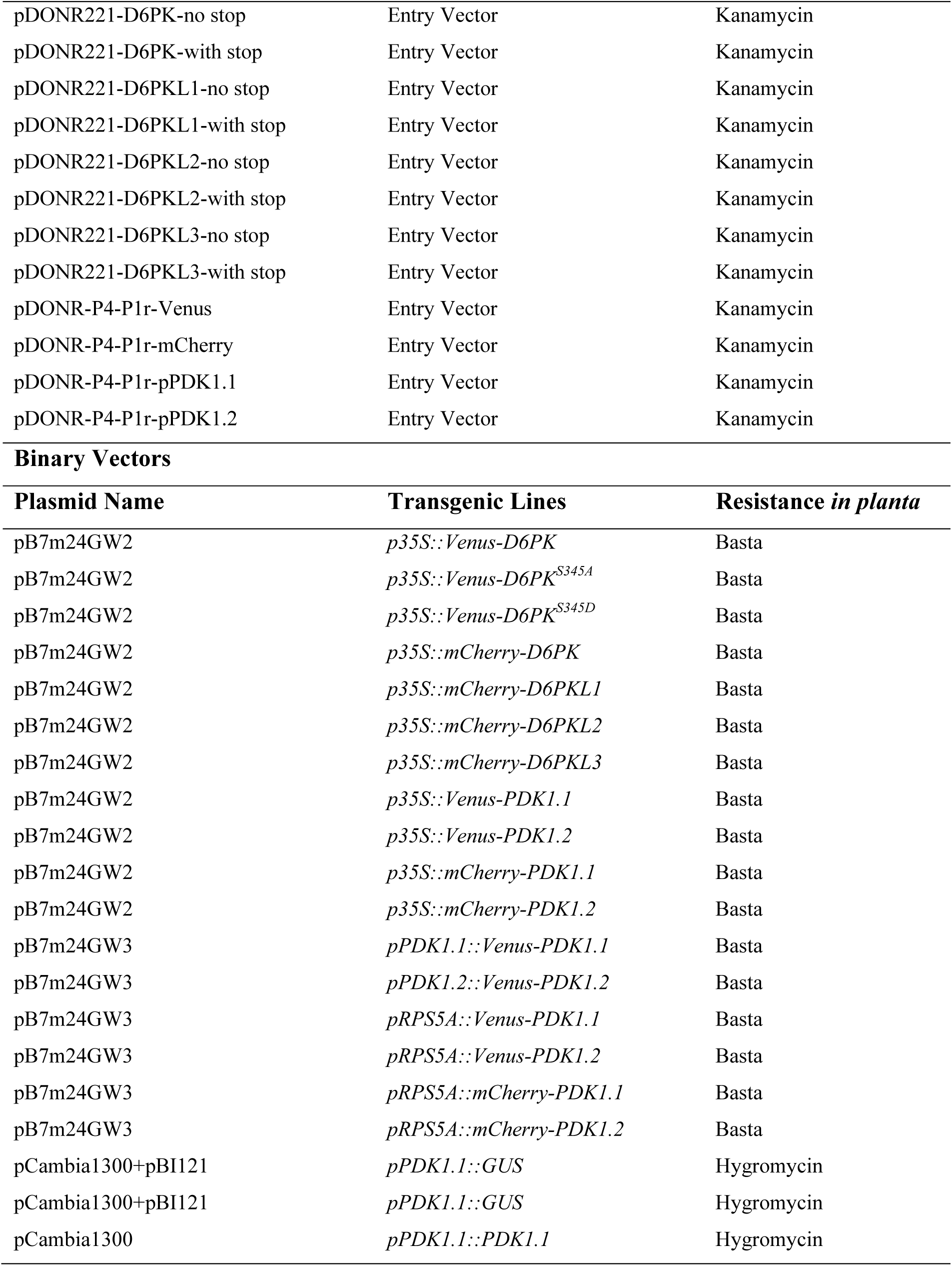
List of plasmids used in this study.

