## Supplemental Figures and Tables for "A lipid code-dependent phosphoswitch directs PIN-mediated auxin efflux in *Arabidopsis* development"

#### Supplemental Figure 1-Figure 1 related

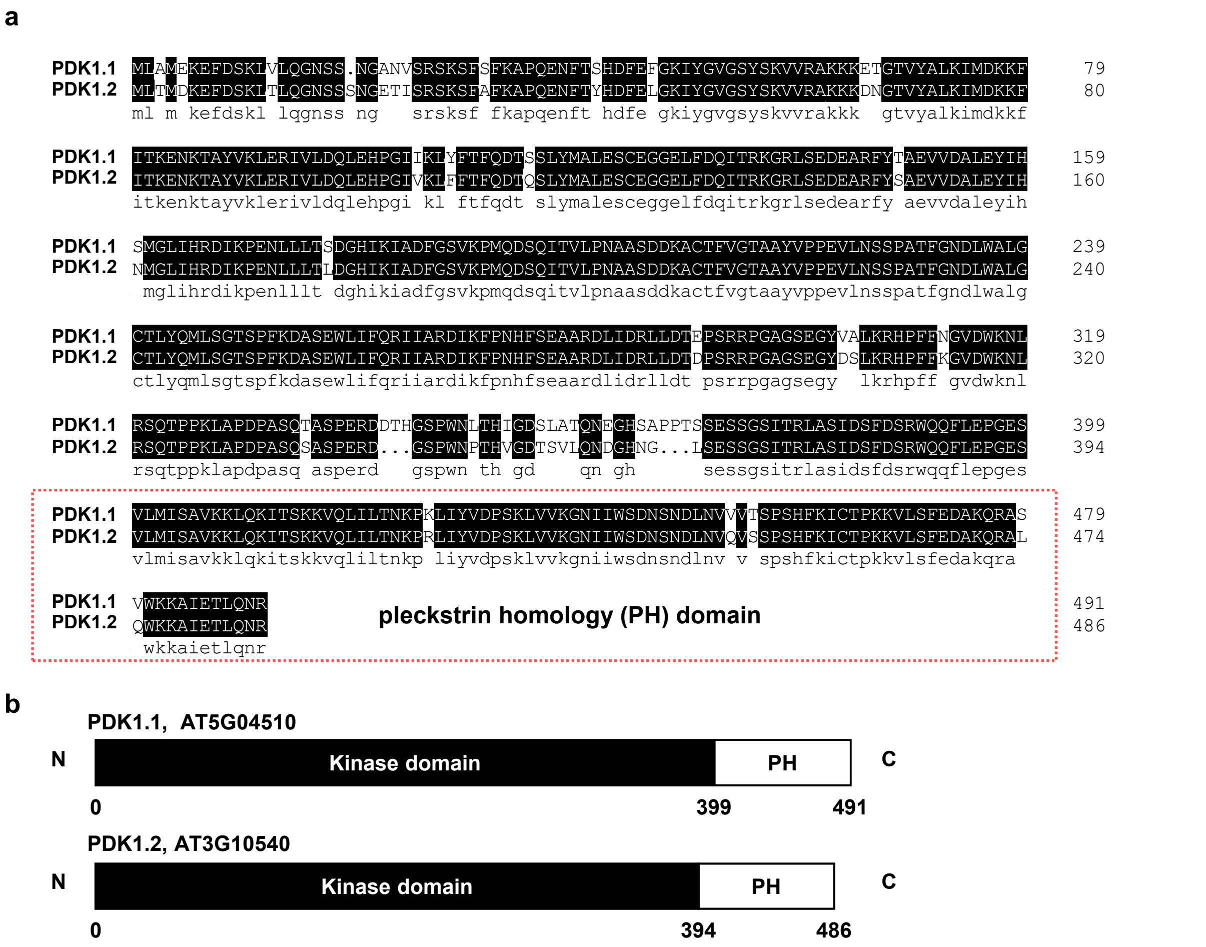


**Supplementary Figure 1. Alignment of the amino acid sequences and schematic graphs of PDK1.1 and PDK1.2.**

a. Alignment of PDK1.1 and PDK1.2 by the DNAMAN program. The two paralogues show 91.06% identity. The predicted pleckstrin homology (PH) domain is indicated by a red rectangle.

b. Schematic graphs of the domain structure of PDK1.1 and PDK1.2 proteins.

#### Supplemental Figure 2-Figure 1 related

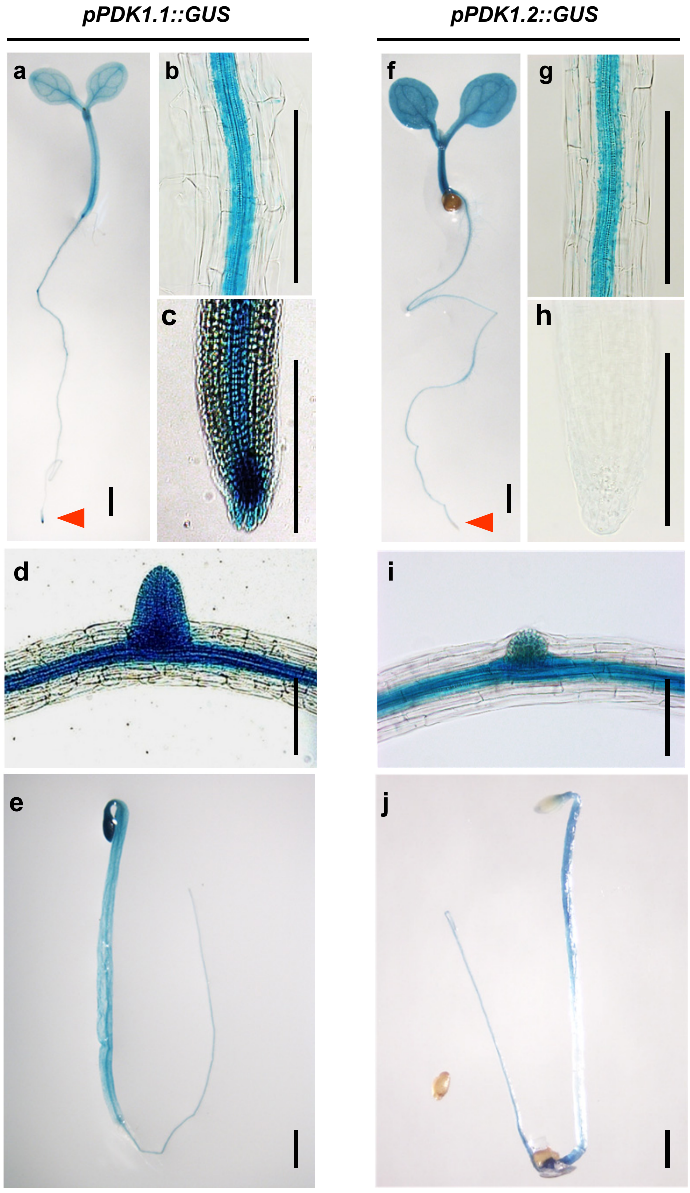


**Supplementary Figure 2. Expression pattern of *PDK1.1* and *PDK1.2*.**

GUS staining of *pPDK1.1::GUS* and *pPDK1.2::GUS* lines indicated that *PDK1.1* and *PDK1.2* were expressed in the vascular tissues in both roots and shoots at various developmental stages, including young seedlings (a, f; 7 days), root stele (b, g; 7 days), columella cells (c, h, only with expression detected for *PDK1.1*; 7 days), lateral root primordia (d, i; 12 days), and dark-grown seedlings (e, j; 4 days). Representative images of three independent homozygous lines were shown. Scale bars, 1 mm.

### Supplemental Figure 3-Figure 1 related

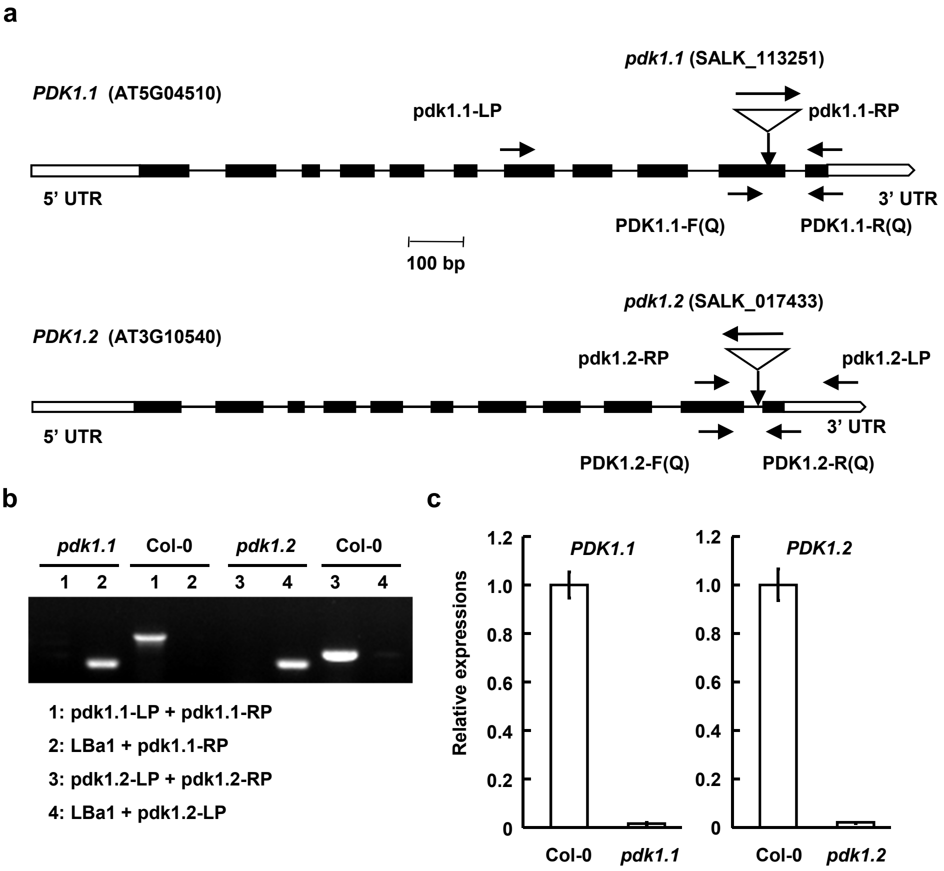


**Supplementary Figure 3. Identification of *Arabidopsis pdk1.1* and *pdk1.2* T-DNA insertional mutants.**

a. Schematic representation of *PDK1.1* and *PDK1.2* genes and positions of T-DNA insertions for *pdk1.1* and *pdk1.2*. Introns, exons, and non-coding regions are indicated by lines, black, or blank boxes respectively. Positions of primers are indicated.

b. Identification of homozygous *pdk1.1* and *pdk1.2* mutants. Genomic DNA of *pdk1.1* and *pdk1.2* mutants was used as templates for PCR amplification. Homozygous lines have a single amplified DNA fragment when using LBa1/pdk1.1-RP or LBa1/pdk1.2-RP primers.

c. qRT-PCR analysis confirmed the deficient expression of *PDK1.1* and *PDK1.2* genes in *pdk1.1* and *pdk1.2* mutants, respectively. Total RNA of 7-day-old WT, *pdk1.1* and *pdk1.2* seedlings was extracted, reversely transcribed, and then used for analysis. *ACTIN7* was amplified and used as an internal reference to normalize the expression of *PDK1.1* and *PDK1.2*. The experiments were biologically repeated for 3 times and results were presented as mean ± s.d..

#### Supplemental Figure 4-Figure 1 related

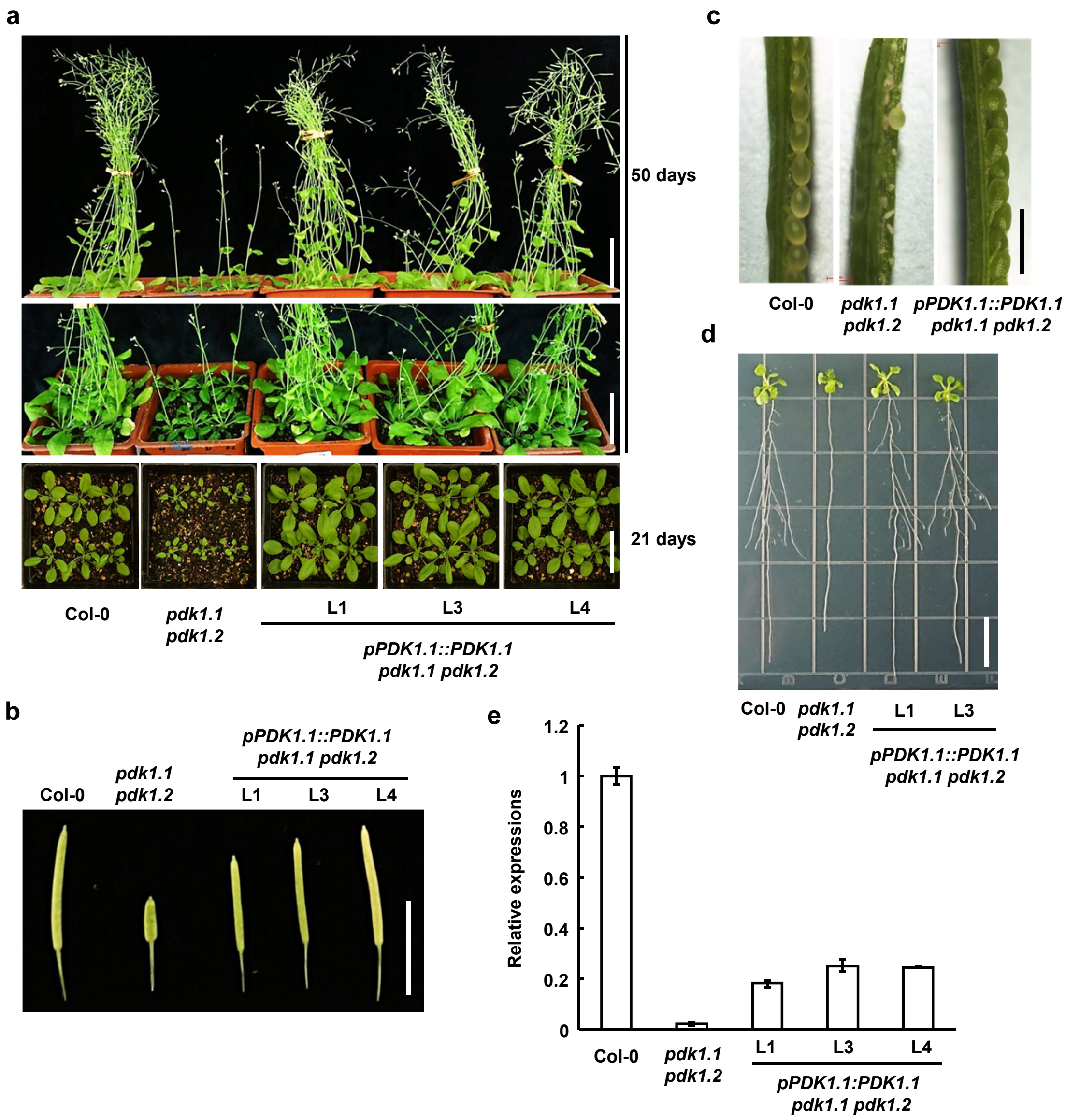


**Supplementary Figure 4. Expression of *PDK1.1* driven by native promoter (*pPDK1.1::PDK1.1*) rescued the defective growth of *pdk1.1 pdk1.2* double mutant.**

a-d. Complementary expression of *PDK1.1* recovered the reduced growth (a, scale bar, 5 cm), silique shortness (b; scale bar, 1 cm), low fertility (c; scale bar, 1 mm), and defective root elongation and lateral root formation (d, representative images of 2-week-old seedlings were shown; scale bar, 1 cm) of *pdk1.1 pdk1.2*.

e. qRT-PCR analysis showed the complementary expression of *PDK1.1* in *pdk1.1 pdk1.2* plants. Total RNA of 7-day-old WT, *pdk1.1 pdk1.2* and transgenic *pdk1.1 pdk1.2* seedlings harbouring *pPDK1.1::PDK1.1* was extracted and used for analysis. *ACTIN7* was amplified and used as an internal reference. Experiments were biologically repeated and data were presented as mean ± s.d. (n = 3).

#### Supplemental Figure 5-Figure 1 related

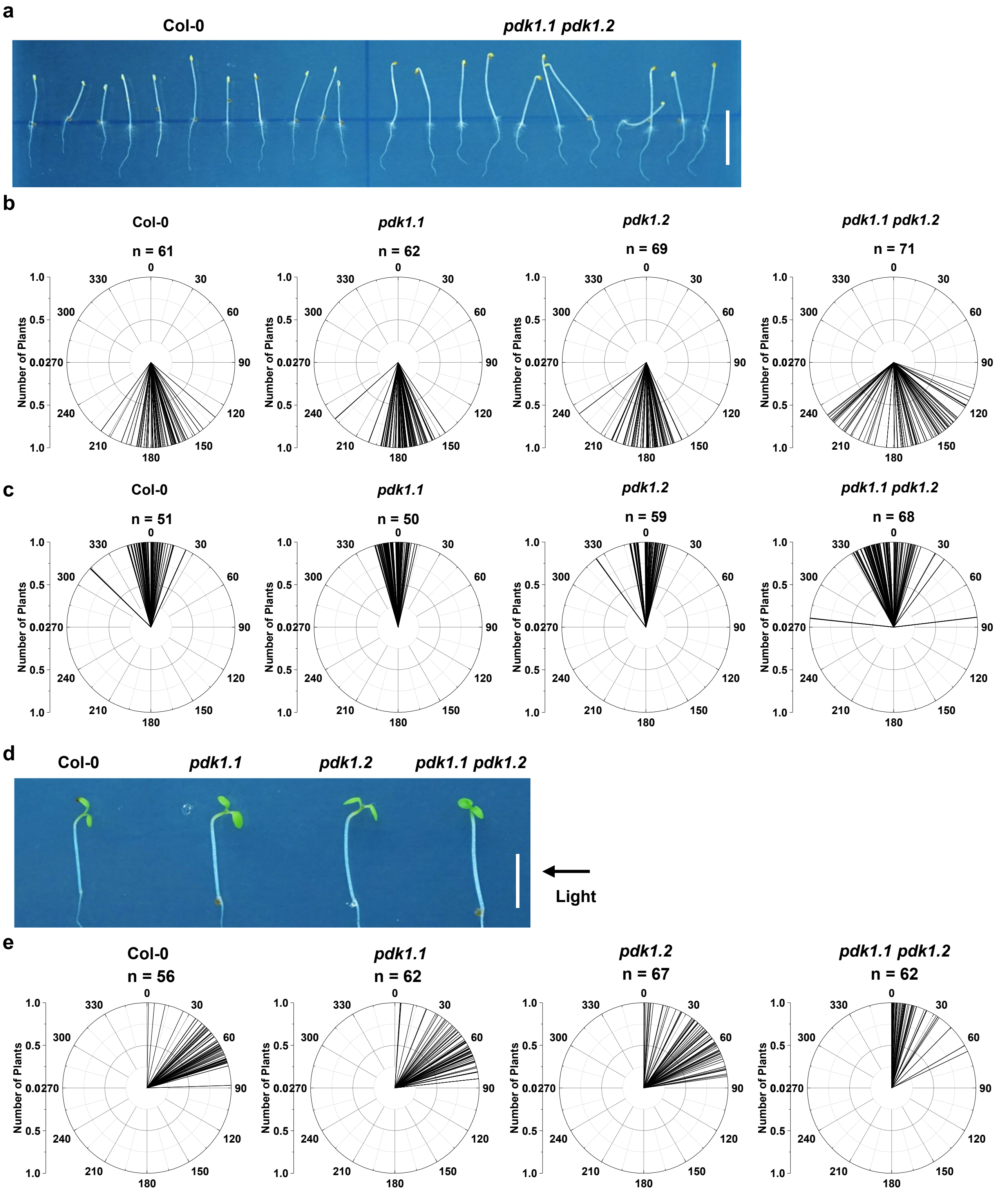


**Supplementary Figure 5. Deficiency of *PDK1.1* and *PDK1.2* impaired the hypocotyl gravitropism under dark and phototropism towards directional light.**

a. Deficiency of *PDK1.1* and *PDK1.2* impaired root and shoot gravitropic response under dark. Etiolated seedlings of Col-0, *pdk1.1*, *pdk1.2*, and *pdk1.1 pdk1.2* were grown under dark for 90 h and a representative photo was shown. Scale bar, 5 mm.

b. Deficiency of *PDK1.1* and *PDK1.2* impaired root gravitropic response under dark. Etiolated seedlings of Col-0, *pdk1.1*, *pdk1.2*, and *pdk1.1 pdk1.2* were grown under dark for 90 h and root tip angles were measured by Image J and data were visualized by polar bar charts. n = 61, 62, 69, and 71, respectively.

c. Deficiency of *PDK1.1* and *PDK1.2* led to reduced gravitropic hypocotyls under dark. Etiolated seedlings of Col-0, *pdk1.1*, *pdk1.2*, and *pdk1.1 pdk1.2* were grown under dark for 90 h and hypocotyl angles were measured by Image J and data were visualized by polar bar charts with the Origin program. n = 51, 50, 59, and 68, respectively.

d. *pdk1.1 pdk1.2* showed defects in phototropism. Seedlings of Col-0, *pdk1.1*, *pdk1.2,* and *pdk1.1 pdk1.2* were grown under dark for 90 h, exposed to white light for 24 h, and were then subjected to directional white light in a box covered with aluminium foil from the other sides. Scale bar, 5 mm.

e. Deficiency of *PDK1.1* and *PDK1.2* impaired hypocotyl phototropic response towards direction light source. As indicated in (c), hypocotyl angles were measured by Image J and data were visualized by polar bar charts. n = 56, 62, 67, and 62, respectively.

### Supplemental Figure 6-Figure 1 related

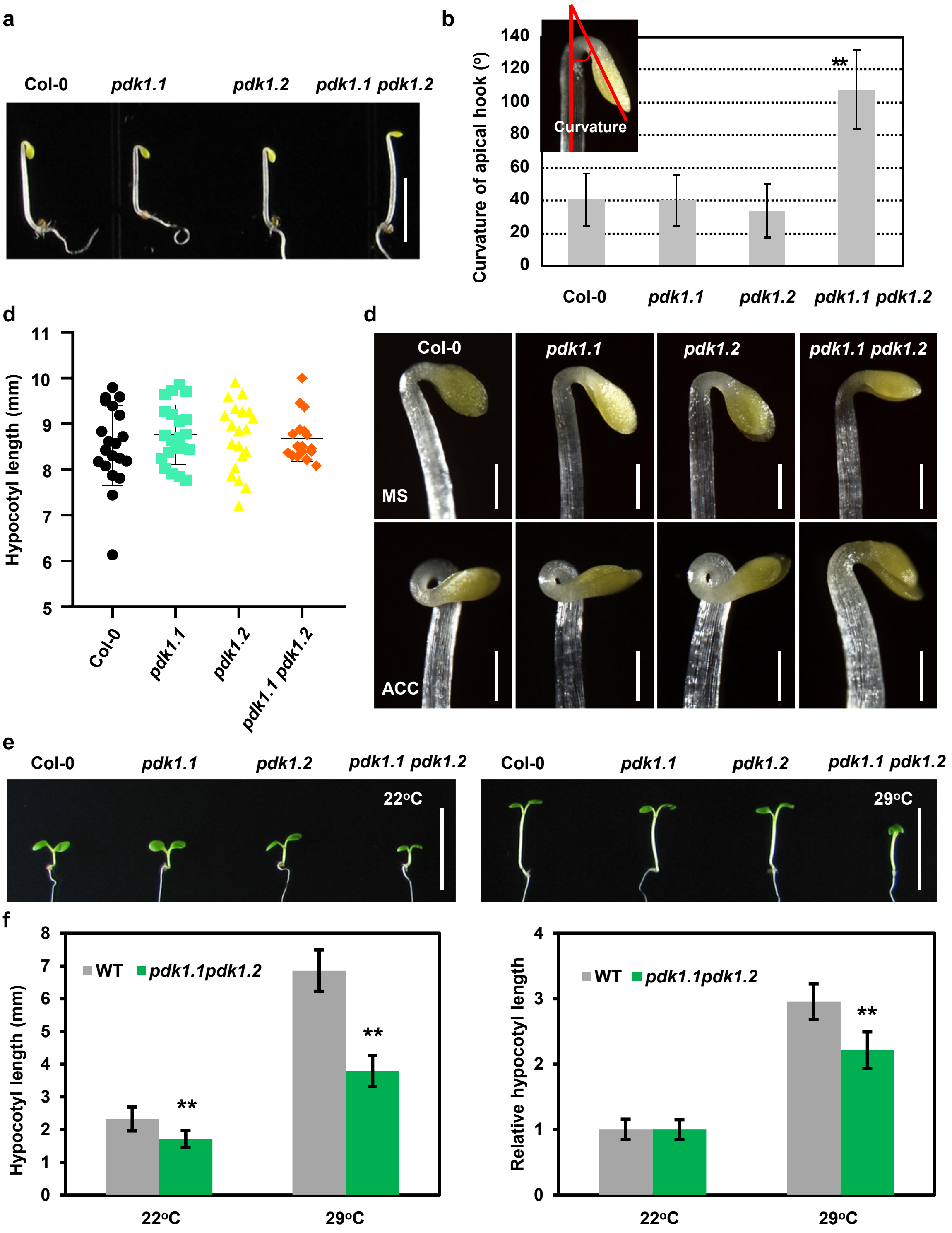


**Supplementary Figure 6. Deficiency of *PDK1.1* and *PDK1.2* impaired the normal development of the apical hook and high temperature-induced hypocotyl elongation.**

a-b. Observation (a, scale bar, 5 mm) and quantification (b) showed that etiolated seedlings of *pdk1.1 pdk1.2* exhibited less tight apical hooks. Col-0, *pdk1.1*, *pdk1.2,* and *pdk1.1 pdk1.2* seedlings were grown under dark for 90 h. Angles were measured by Image J. **, p<0.01, Student's t-test.

c. Etiolated seedlings of *pdk1.1 pdk1.2* exhibited comparably long hypocotyls to Col-0. Seedlings were grown under dark for 90 h. No significant difference was revealed by detected a Student's t-test.

d. Etiolated seedlings of *pdk1.1 pdk1.2* did not form apical hooks in the presence of ACC. Seedlings of Col-0, *pdk1.1*, *pdk1.2*, and *pdk1.1 pdk1.2* were grown under dark for 90 h in the absence or presence of ACC (10 µM). Scale bars, 500 µm.

e-f. *pdk1.1 pdk1.2* showed defects in high temperature-induced hypocotyl elongation. Seedlings of Col-0, *pdk1.1*, *pdk1.2* and *pdk1.1 pdk1.2* were grown under light at 22° C or 29° C for 5 days, and hypocotyl elongation was observed (e, scale bar, 1 cm) and quantified (e). Hypocotyl length was measured by the Image J program and shown as mean ± s.d. (left, n = 20) or relative length by setting the hypocotyl length of Col-0 and *pdk1.1 pdk1.2* at 22° C as "1", respectively (right, n = 20). **, p<0.01, Student's t-test.

### Supplemental Figure 7-Figure 2 related

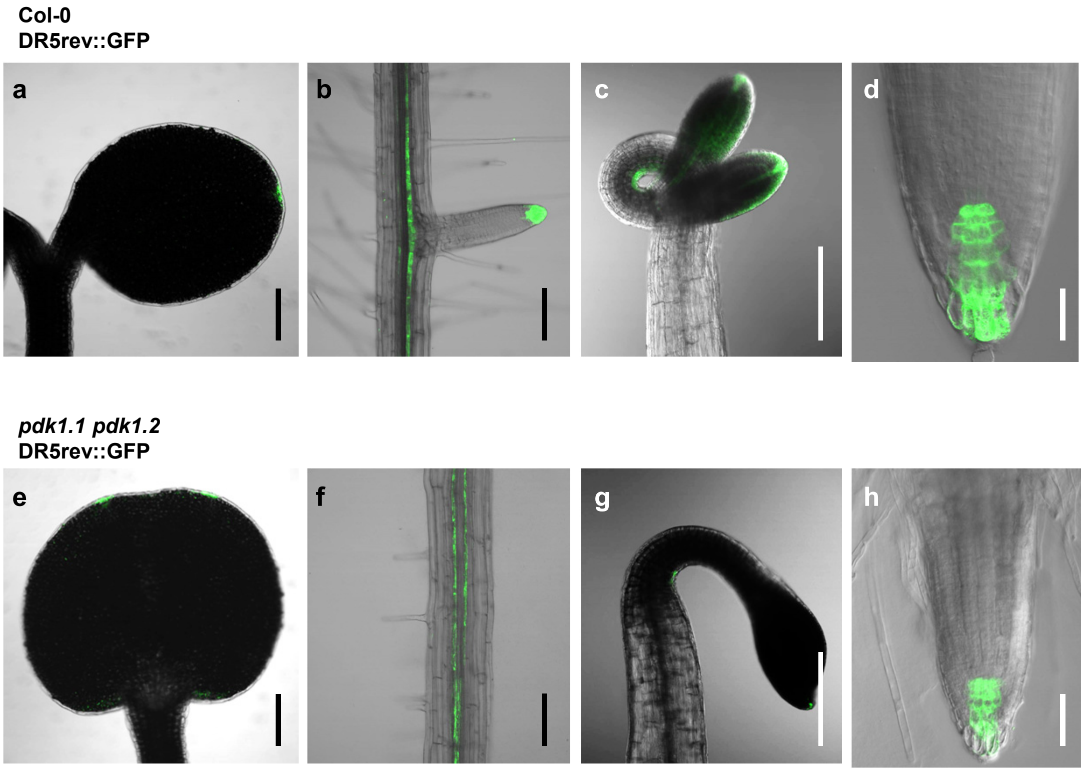


**Supplementary Figure 7. Loss of function of *PDK1.1* and *PDK1.2* impaired auxin distribution.**

Observation of the auxin responsive reporter DR5rev::GFP by CLSM indicated a dramatic decrease of the auxin maxima in *pdk1.1 pdk1.2* (e-h) compared with Col-0 (a-d). Fused cotyledon exhibiting two sites of auxin maxima in light-grown 7-day-old seedlings of *pdk1.1 pdk1.2* (e) compared with one of Col-0 (a); roots of light-grown 10-day-old seedlings (b and f); apical hooks of 10 μM ACC treated 4-day-old etiolated seedlings (c and g); roots of 10 μM ACC treated 4-day-old etiolated seedlings (d and h). Scale bars, 200 μm.

#### Supplemental Figure 8-Figure 2 related

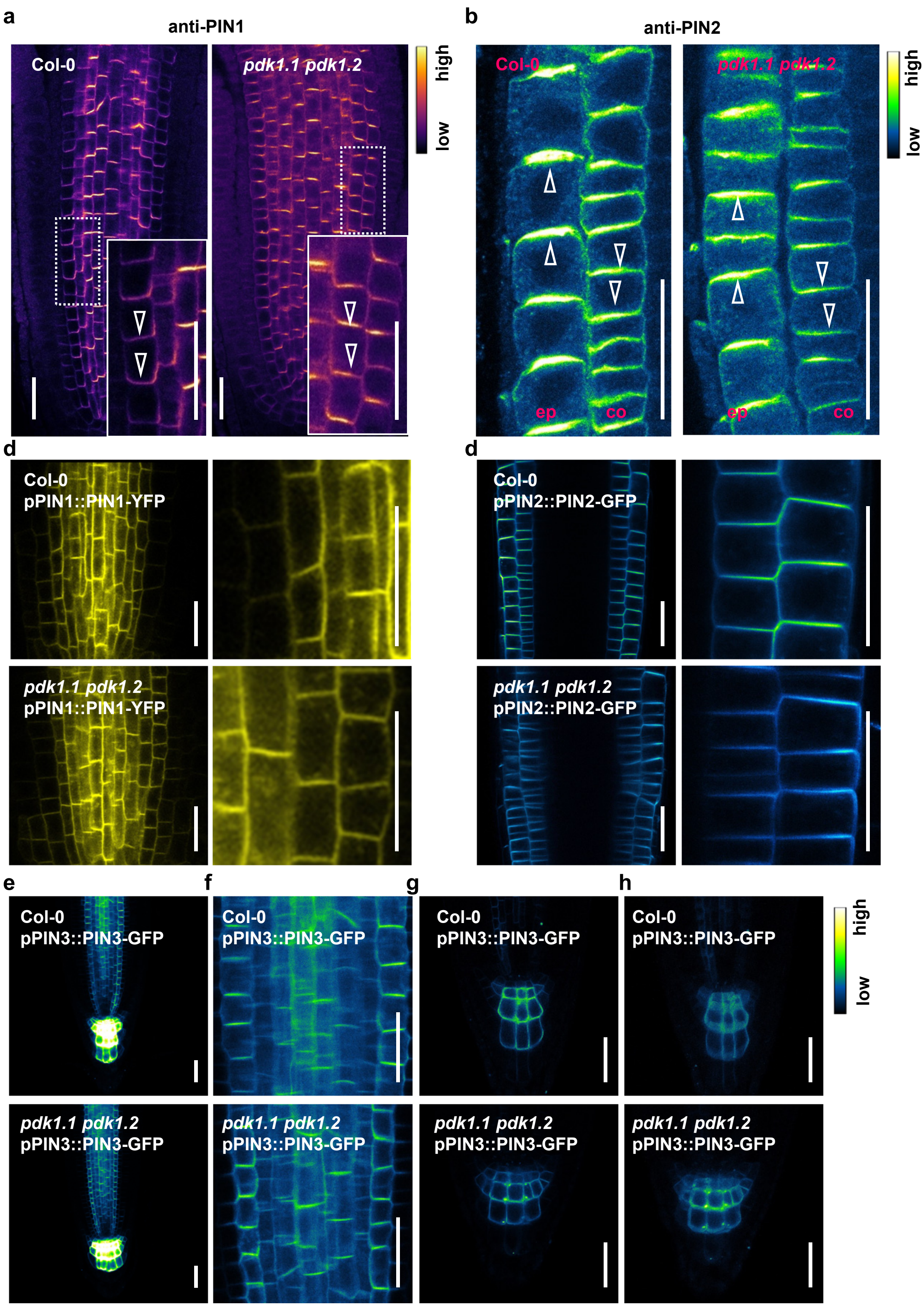


**Supplementary Figure 8. Deficiency of *PDK1.1* and *PDK1.2* did not affect the polarity of PIN proteins.**

a. Deficiency of *PDK1.1* and *PDK1.2* did not change the polarity of PIN1 in roots. Four-day-old seedlings of Col-0 and *pdk1.1 pdk1.2* were used for immunofluorescence with a rabbit anti-PIN1 antibody (1: 500), and then imaged by CLSM. The "mpl-inferno" LUT was used for photo visualization based on fluorescence intensity by the FIJI program. Arrowheads indicated basal localization. Scale bars, 20 μm.

b. Deficiency of *PDK1.1* and *PDK1.2* did not change the polarity of PIN2 in roots. Four-day-old seedlings of Col-0 and *pdk1.1 pdk1.2* were used for immunofluorescence with a rabbit anti-PIN2 antibody (1: 500), and then imaged by CLSM. The "Green Fire Blue" LUT was used for photo visualization based on fluorescence intensity by the FIJI program. Arrowheads indicated apical localization in epidermis ("ep") and the basal localization in cortex ("co") respectively. Scale bars, 20 μm.

c. Deficiency of *PDK1.1* and *PDK1.2* did not change the polarity of PIN1-YFP. Four-day-old seedlings of *pPIN1::PIN1-YFP* in Col-0 and *pPIN1::PIN1-YF*P in *pdk1.1 pdk1.2* were imaged by CLSM. Scale bars, 20 μm.

d. Deficiency of *PDK1.1* and *PDK1.2* did not change the polarity of PIN2-GFP. Four-day-old seedlings of *pPIN2::PIN2-GFP* in Col-0 and *pPIN2::PIN2-GFP* in *pdk1.1 pdk1.2* were imaged by CLSM. Scale bars, 20 μm.

e-h. Deficiency of *PDK1.1* and *PDK1.2* did not change the polarity of PIN3-GFP. Four-day-old seedlings of *pPIN3::PIN3-GFP* in Col-0 and *pPIN3::PIN3-GFP* in *pdk1.1 pdk1.2* were imaged by CLSM. (e) subcellular localization of PIN3-GFP; (f) an amplified view of PIN3-GFP in the root stele; (g) a close view of PIN3-GFP in root columella cells; (h) a 3D-projection of PIN3-GFP localization in root columella cells. The "Green Fire Blue" LUT was used for photo visualization based on fluorescence intensity by the FIJI program. Scale bars, 20 μm.

#### Supplemental Figure 9-Figure 2 related

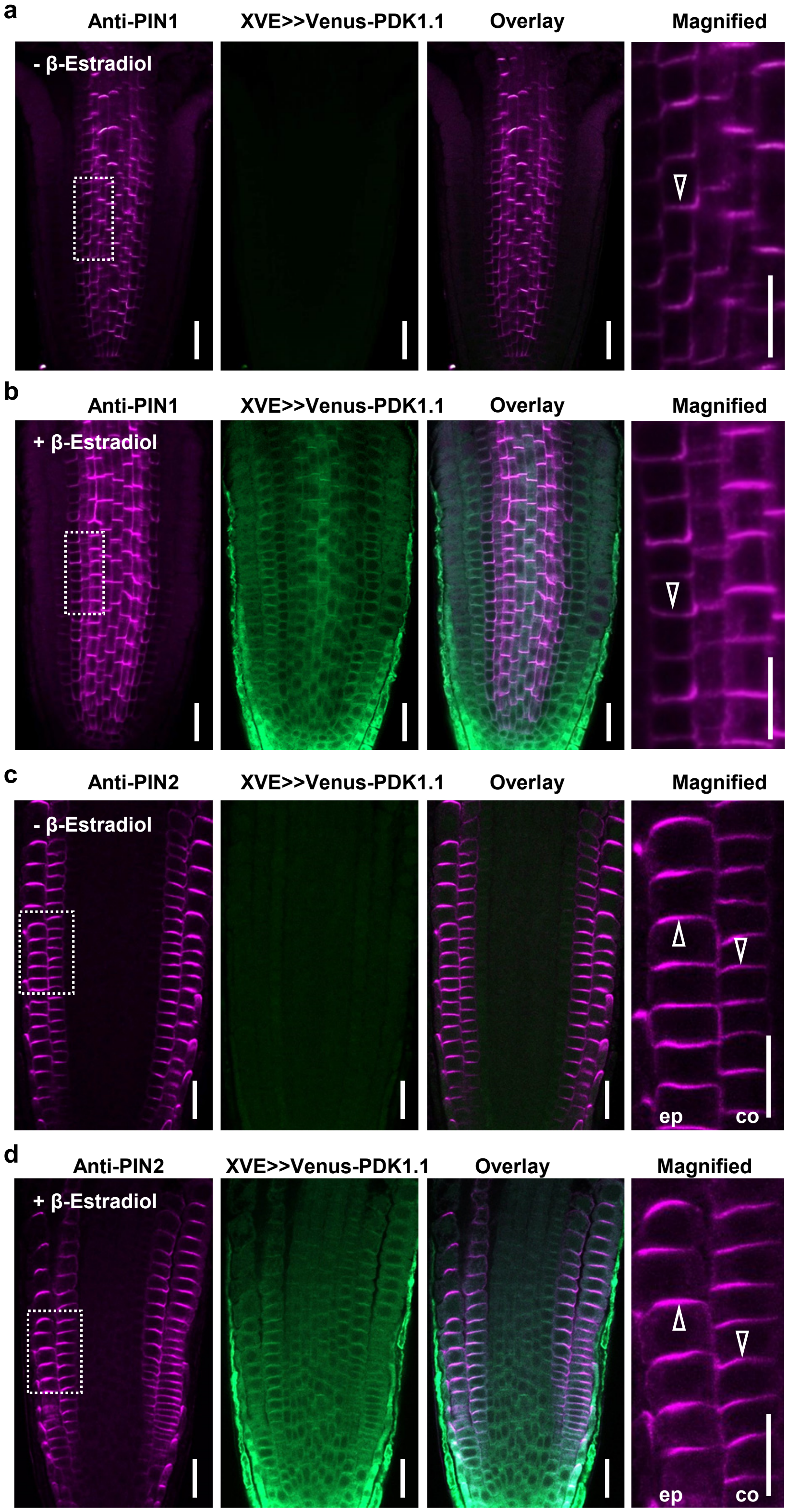


**Supplementary Figure 9. Overexpression of *PDK1.1* did not affect the polarity of PIN1.**

a-b. Induced overexpression of *PDK1.1* by β-Estradiol did not change the polar localization of PIN1. Four-day-old *XVE>>Venus-PDK1* (in Col-0) seedlings were transferred to MS plates without (a) or with (b) 5 µM β-Estradiol for 12 h, then used for immunofluorescence with a rabbit anti-PIN1 antibody (1: 500) and imaged by CLSM. Open arrowheads indicated the basal localization of PIN1. Scale bar, 20 µm.

c-d. Induced overexpression of *PDK1.1* did not change the polar localization of PIN2. Same set of samples as in (a-b) were conducted for immunofluorescence with a rabbit anti-PIN2 antibody (1: 500) and imaged by CLSM. Open arrowheads indicated the apical or the basal localization of PIN2 in epidermis (short as “ep”) or cortex (“co”) respectively. Scale bar, 20 µm.

#### Supplemental Figure 10-Figure 3 related

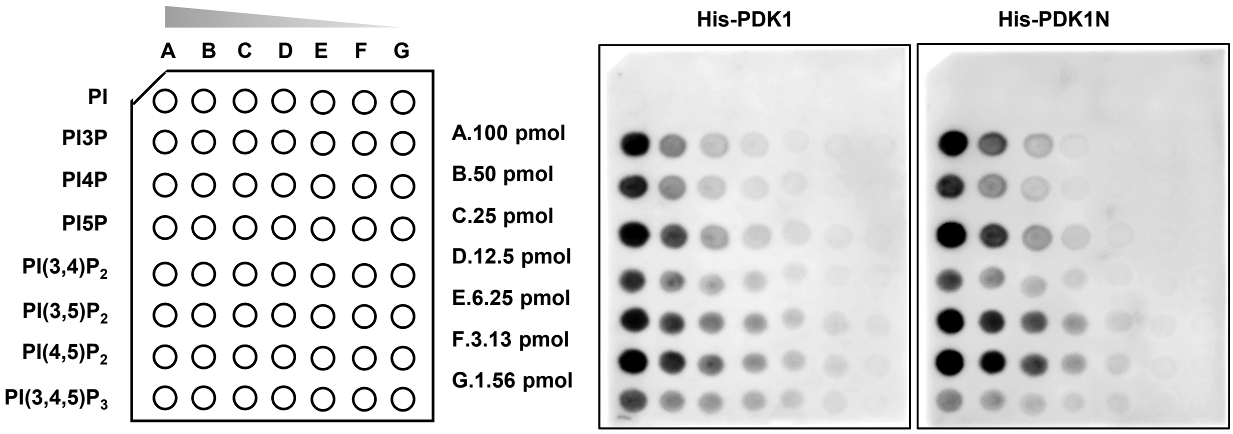


**Supplementary Figure 10. His-PDK1.1 and His-PDK1N bound to a similar spectrum of phospholipids**

Lipid-protein blot overlay assays with PIP arrays revealed that recombinant His-PDK1.1 and His-PDK1.1N bound to similar spectrum of phospholipids. PIP arrays (from left to right: 100 pmol, 50 pmol, 25 pmol, 12.5 pmol, 6.25 pmol, 3.13 pmol, and 1.56 pmol, respectively) were incubated with His-tagged recombinant PDK1.1 or PDK1.1N proteins, and detected by a His antibody.

#### Supplemental Figure 11-Figure 3 related

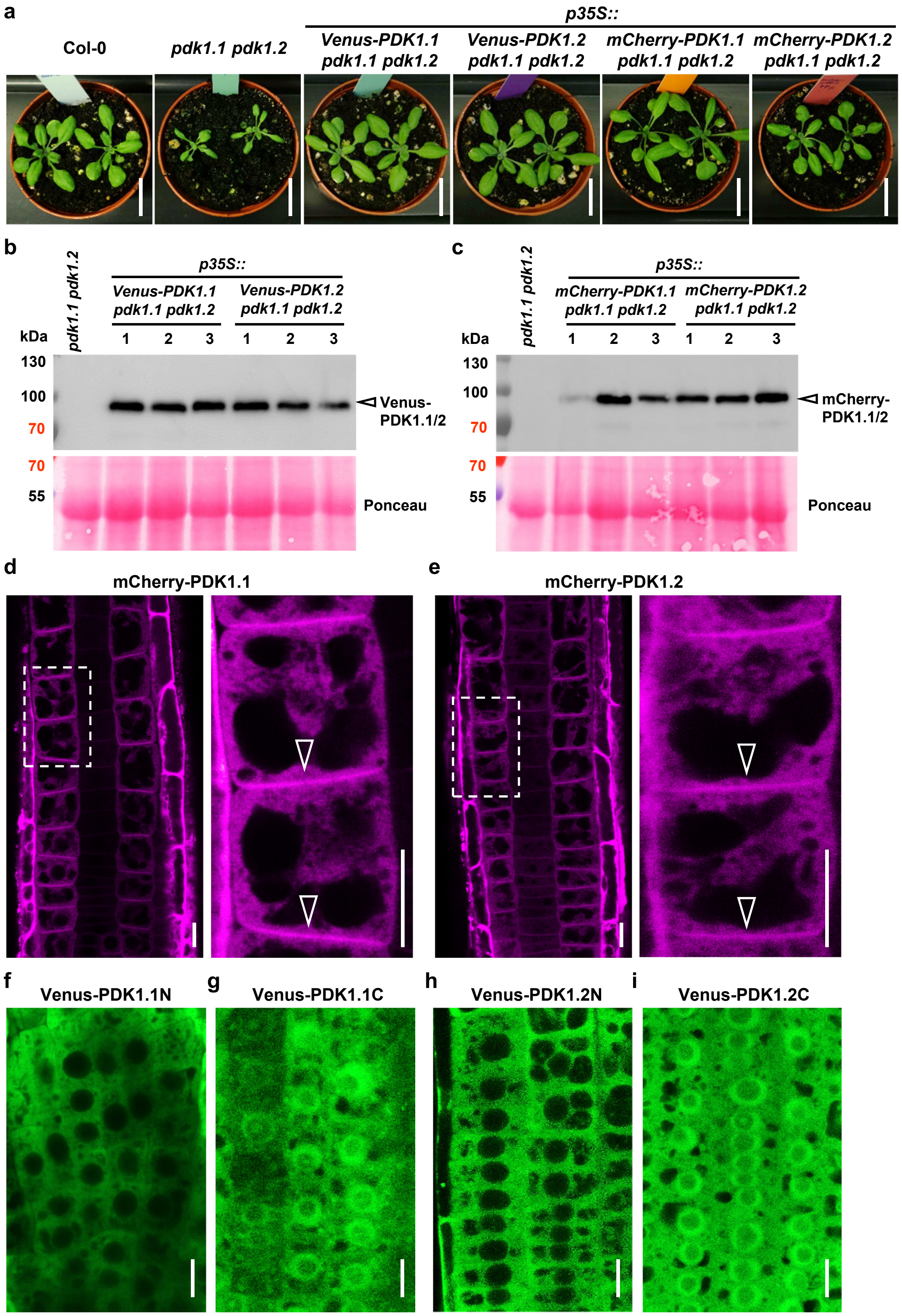


**Supplementary Figure 11. Analysis of *PDK1* transgenic lines.**

a. Expression of *PDK1.1* or *PDK1.2* (*p35S::Venus-PDK1.1, p35S::Venus-PDK1.2, p35S::mCherry-PDK1.1* and *p35S:: mCherry-PDK1.2*) rescued the growth defects of *pdk1.1 pdk1.2*. Adult plants (25-day-old) were observed and representative photos were shown. Scale bar, 2 cm.

b. Western blot analysis verified the PDK1.1 or PDK1.2 expression (*35S::Venus-PDK1.1* and *35S::Venus-PDK1.2*) in *pdk1.1 pdk1.2*, respectively. Seven-day-old $T_3$ homozygous seedlings were used for protein extraction and subjected to analysis. Upper panel, anti-GFP (1:2000); lower panel, Ponceau staining.

c. Western blot verified the PDK1.1 or PDK1.2 expression (*35S::mCherry-PDK1.1* and *35S::mCherry-PDK1.2*) in *pdk1.1 pdk1.2*, respectively. Seven-day-old $T_3$ homozygous seedlings were used for protein extraction and subjected to analysis. Upper panel, anti-RFP (1:2000); lower panel, Ponceau staining.

d-e. mCherry-fused PDK1.1 (d) and PDK1.2 (e) localized to both cytoplasm and the basal side of PM. Four-day-old seedlings of *p35S::mCherry-PDK1.1* and *p35S::mCherry-PDK1.2* were imaged by CLSM. Open arrowheads indicated the basal polar localization. Scale bar, 10 µm.

f-i. Subcellular localization of Venus-fused PDK1.1N (f, cytoplasm), Venus fused PDK1.1C (g, both cytoplasm and nucleus), Venus fused PDK1.2N (h, cytoplasm), and Venus fused PDK1.2C (i, both cytoplasm and nucleus). Four-day-old seedlings expressing corresponding fusion proteins were imaged by CLSM. Scale bar, 10 µm.

#### Supplemental Figure 12-Figure 3 related

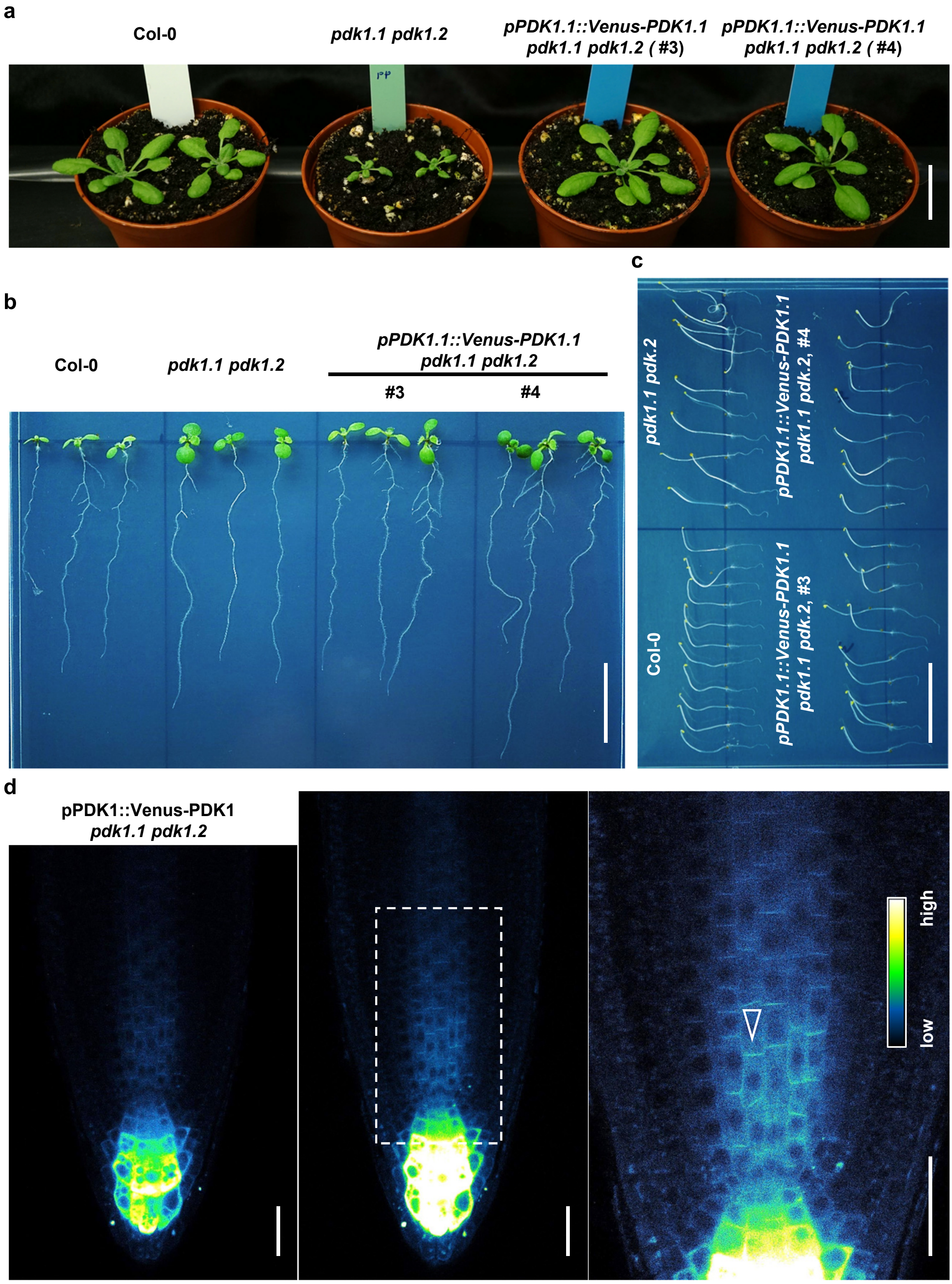


**Supplementary Figure 12. Functional *pPDK1.1::Venus-PDK1.1* localized at both cytoplasm and the basal side of PM.**

a. *pPDK1.1::Venus-PDK1.1* rescued the growth defects of *pdk1.1 pdk1.2*. 25-day-old adult plants were observed and representative photos are shown. Scale bar, 2 cm.

b. *pPDK1.1::Venus-PDK1.1* rescued the lateral root defects of *pdk1.1 pdk1.2*. 10-day-old seedlings were observed and representative photos are shown. Scale bar, 2 cm.

c. *pPDK1.1::Venus-PDK1.1* rescued the defects of *pdk1.1 pdk1.2* in the hypocotyl gravitropic response. 4-day-old etiolated seedlings were observed and representative photos are shown. Scale bar, 2 cm.

d. Venus-fused PDK1.1 localized to both cytoplasm and PM, especially with a predominant presence at the basal side of PM. Four-day-old seedlings of *pPDK1.1::Venus-PDK1.1* were imaged by CLSM. Open arrowheads indicate the basal polar localization. The “Green Fire Blue” LUT was used for photo visualization based on fluorescence intensity by the FIJI program. Scale bar, 20 µm.

### Supplemental Figure 13-Figure 4 related

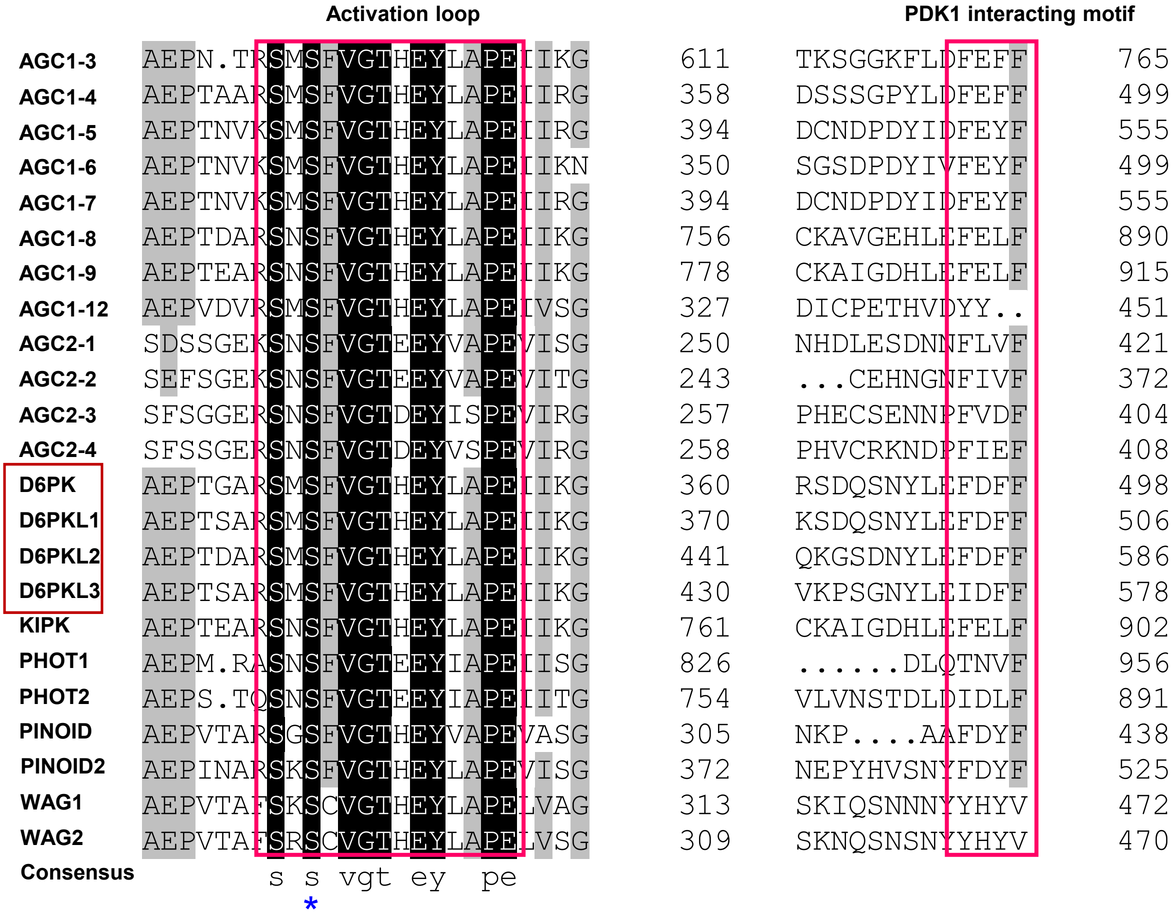


**Supplementary Figure 13. A conserved activation loop and a PDK1 interaction pocket (PIF) motif of AGC protein kinases.**

Alignment of AGC protein kinases was performed with the DNAMAN program, and the similarity was shown in different colours: back, 100% identity; grey, ≥75%. Asterisk indicated a conserved phosphosite (Ser345 in D6PK).

#### Supplemental Figure 14-Figure 4 related

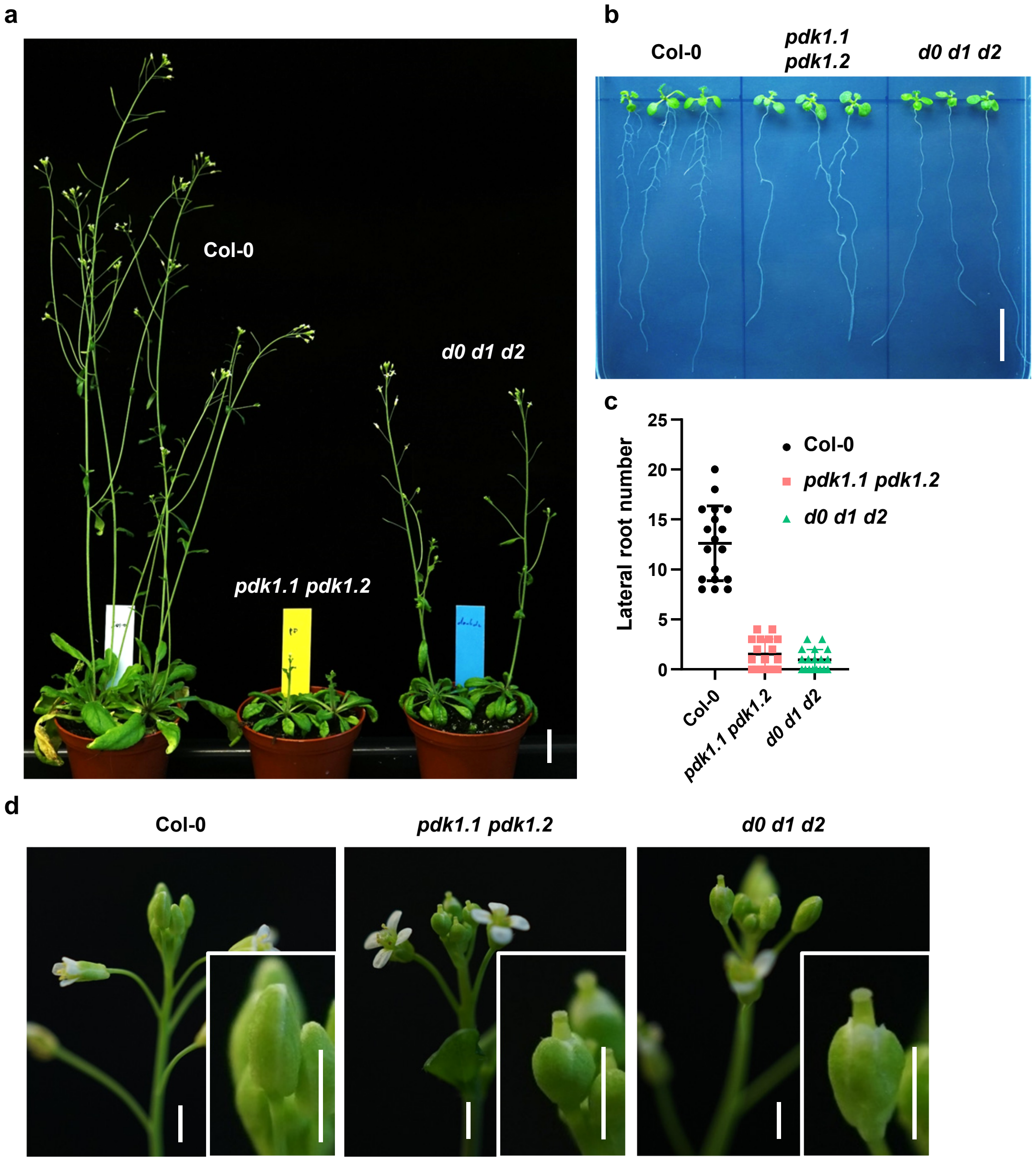


**Supplementary Figure 14. The *d0 d1 d2* triple mutant phenocopied *pdk1.1 pdk1.2*.**

a. Representative photos of 35-day-old Col-0, *d0 d1 d2* and *pdk1.1 pdk1.2* plants. The delayed flowering of *pdk1.1 pdk1.2* was not observed in *d0 d1 d2*. Scale bar, 2 cm.

b-c. *d0 d1 d2* plants showed reduced lateral root number, similar to that of *pdk1.1 pdk1.2*. Twelve-day-old seedlings were observed (b, Scale bar, 1 cm) and number of lateral roots were measured (c). Data are shown as mean ± s.d. (n = 18).

d. Observations showed similar abnormal flowers of *d0 d1 d2* and *pdk1.1 pdk1.2*, with the stigma protruding out of the flower buds before opening. Scale bar, 2 mm.

#### Supplemental Figure 15-Figure 4 related

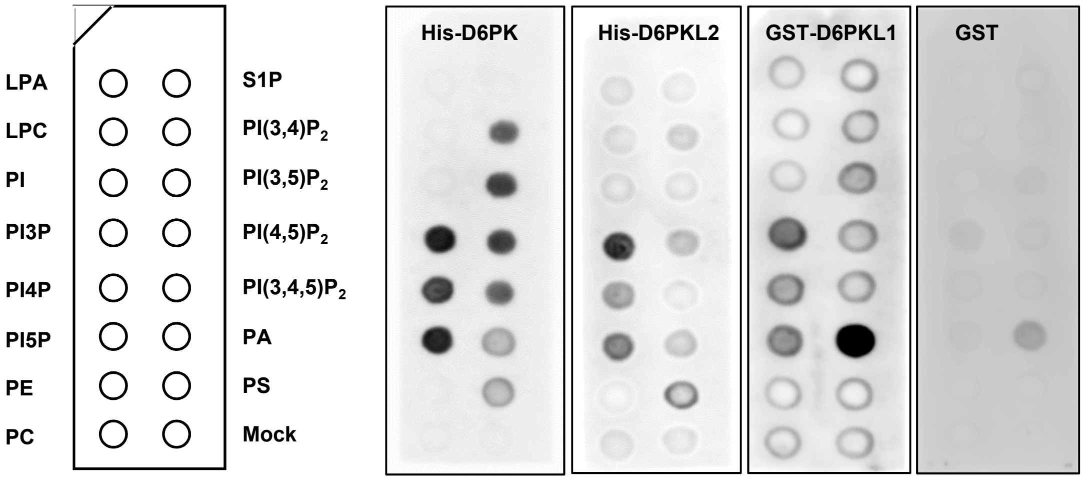


**Supplementary Figure 15. D6PK and D6PKL1/2 (D6PKs) exhibited a similar lipid binding preference as PDK1.1/2.**

Lipid-protein blot overlay assays revealed that D6PKs bound to phospholipids. PIP strips were incubated with recombinant His-D6PK, His-D6PKL2, GST-D6PKL1, and GST (as control) proteins, and detected by a GST or a His antibody accordingly.

### Supplemental Figure 16-Figure 4 related

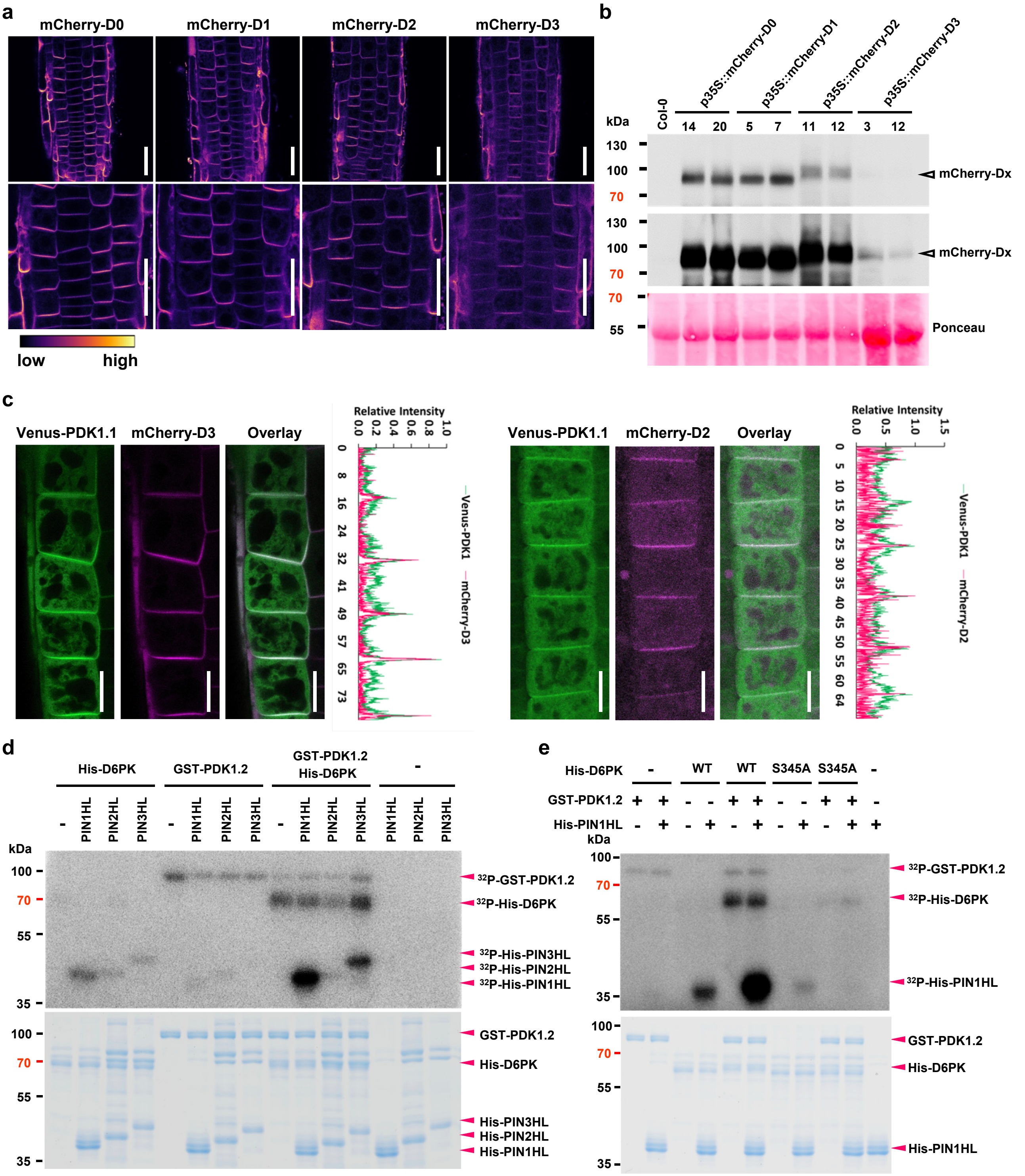


**Supplementary Figure 16. mCherry-D6PKs co-localized with Venus-PDK1.1 at the basal side of PM.**

a. mCherry-fused D6PKs localized to the basal side of PM. Four-day-old seedlings of *p35S::mCherry-D6PK/D6PKLs* (short as *D0 to D3*) were imaged by CLSM. The “mpl-inferno” LUT was used for photo visualization based on fluorescence intensity by the FIJI program. Scale bars, 10 µm.

b. Western blot verified the mCherry-D0~D3 protein level (*35S::mCherry-D0~D3*) in Col-0, respectively. Seven-day-old $T_3$ homozygous seedlings were used for protein extraction and subjected to Western blot analysis. Upper panel, anti-RFP (1:1000), short exposure (0.5 seconds); medium panel, anti-RFP (1:1000), long exposure (5 seconds, for low expression of mCherry-D3); lower panel, Ponceau staining.

c. Venus-PDK1.1 co-localized with mCherry-D6PKL2 and mCherry-D6PKL3 at the basal side of PM. Four-day-old seedlings of *p35S::mCherry-D6PKL2 / p35S::Venus-PDK1.1* and *p35S::mCherry-D6PKL3 / p35S::Venus-PDK1.1* were imaged by CLSM. Scale bar, 10 µm.

d. *In vitro* kinase assay with [$^{32}$P]-ATP revealed that GST-PDK1.2-conducted phosphorylation of D6PK facilitates its activity towards PIN-HL phosphorylation. Upper panel, autoradiography; lower panel, CBB staining.

e. *In vitro* kinase assay with [$^{32}$P]-ATP revealed that GST-PDK1.2-conducted full phosphorylation and activation of D6PK, towards His-PIN1-HL phosphorylation, required the phosphorylation at S345 for D6PK. Upper panel, autoradiography of $^{32}$P; lower panel, CBB staining.

### Supplemental Figure 17-Figure 4 related

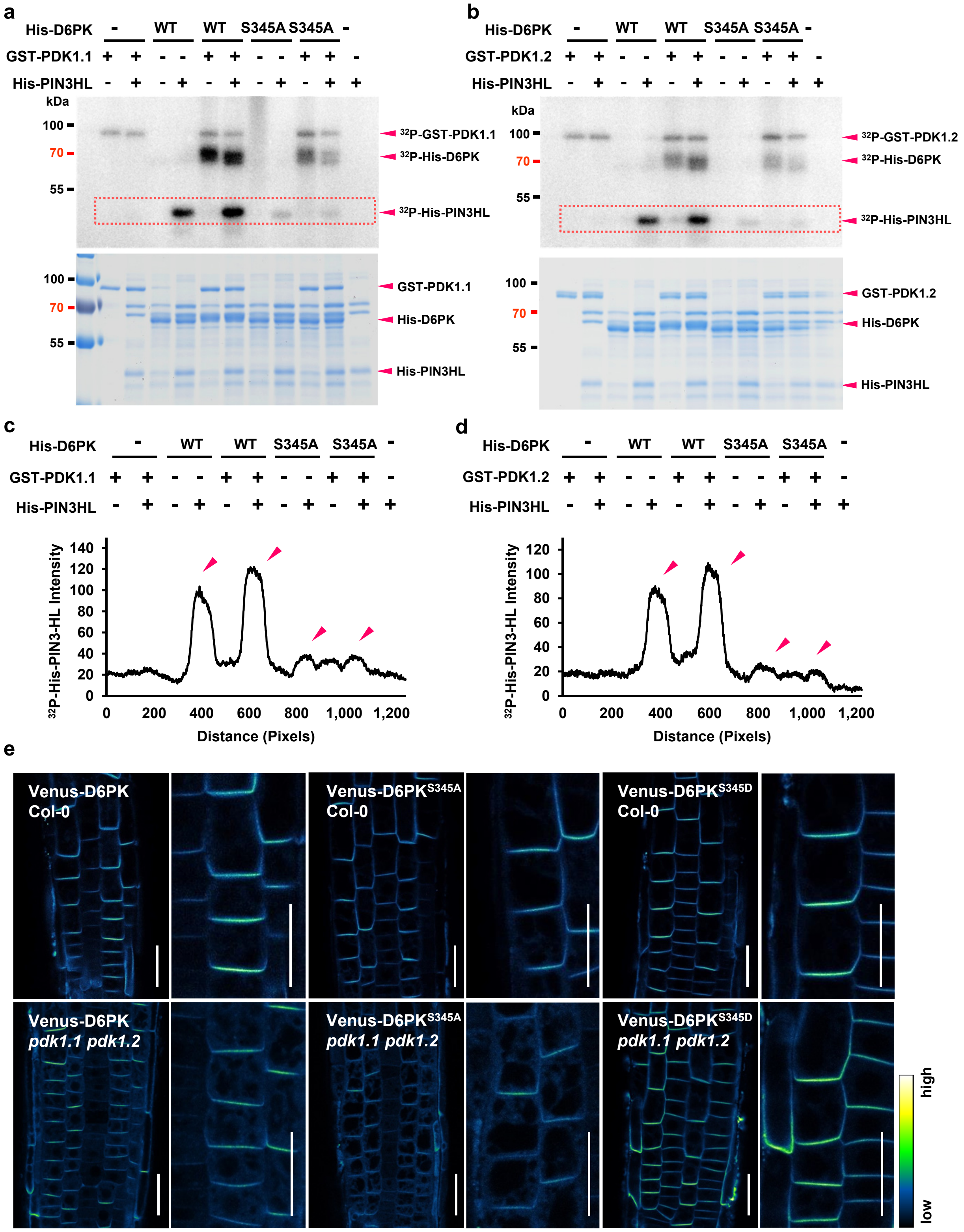


**Supplementary Figure 17. PDK1.1 activated D6PK through phosphorylating Ser345 (S345).**

a-b. *In vitro* kinase assay with [$^{32}$P]-ATP revealed that GST-PDK1.1- (a) and GST-PDK1.2 (b)-conducted full phosphorylation and activation of D6PK, towards His-PIN3-HL phosphorylation, required the phosphorylation at S345 for D6PK. Upper panel, autoradiography of $^{32}$P; lower panel, CBB staining.

c-d. Quantification of the band intensities of D6PK variants' phosphorylation toward PIN3HL by the Image J program, as squared in Supplementary Fig. 17A or B.

e. The subcellular localizations of Venus-D6PK, Venus-D6PK$^{S345A}$, and Venus-D6PK$^{S345D}$ do not show difference in the basal polarity. Four-day-old seedlings of p35S::Venus-D6PK, p35S::Venus-D6PK$^{S345A}$, and p35S::Venus-D6PK$^{S345D}$, in Col-0 and *pdk1.1 pdk1.2* respectively, were imaged by CLSM. The "Green Fire Blue" LUT was used for photo visualization based on fluorescence intensity by the FIJI program. Representative photos of at least three independent transgenic lines were shown. Scale bars, 10 µm.

#### Supplemental Figure 18-Figure 4 related

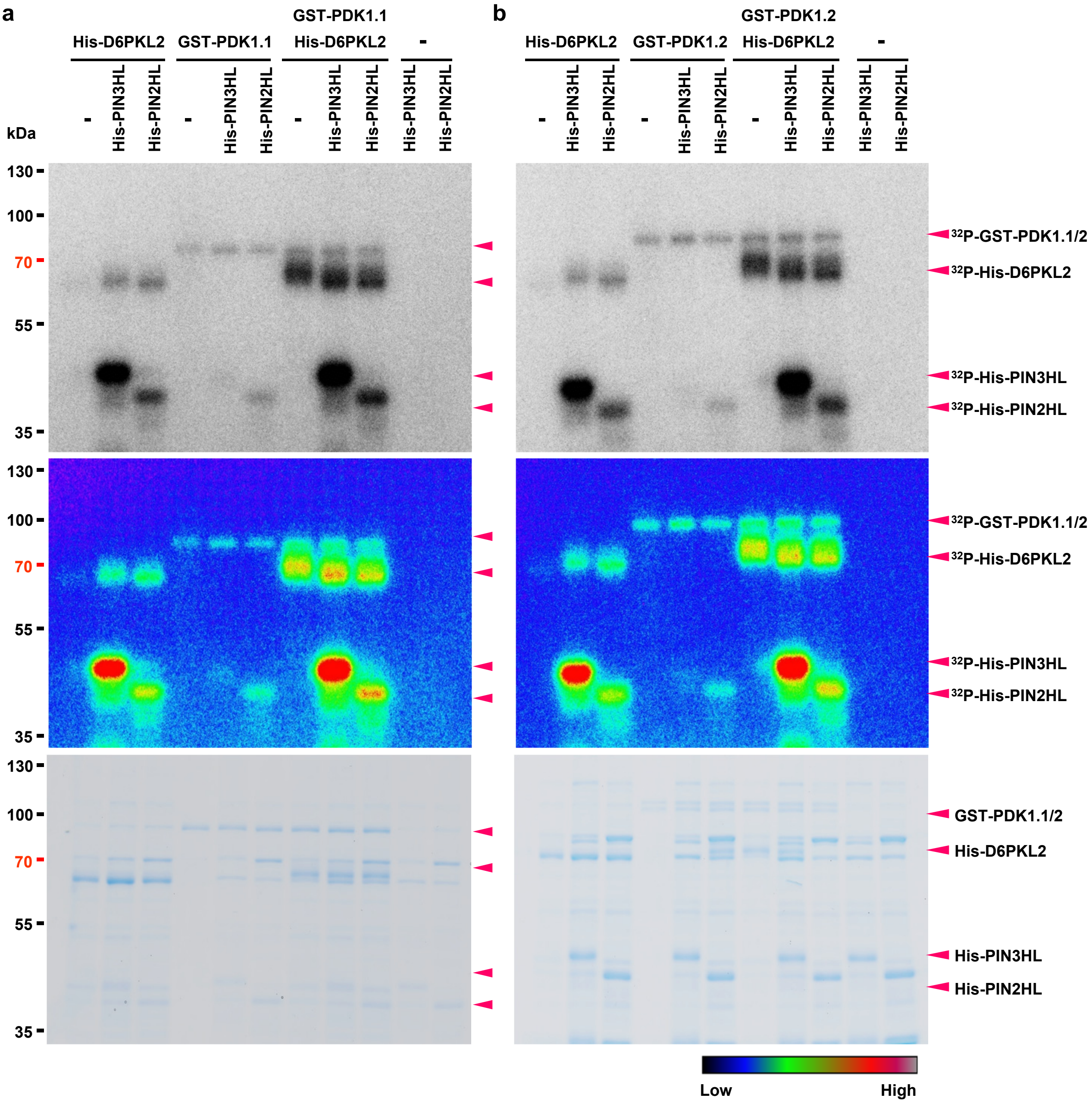


**Supplementary Figure 18. PDK1 phosphorylated D6PKL2 to increase the phosphorylation of PIN3/2-HLs.**

*In vitro* kinase assay with [$^{32}$P]-ATP revealed that GST-PDK1.1 (a) and GST-PDK1.2 (b)-conducted phosphorylation of D6PK facilitated its activity towards PIN3/2-HL phosphorylation. Upper panels, autoradiography of $^{32}$P; Middle panels, the "Rainbow RGB" LUT was used for data visualization based on fluorescence intensity by the FIJI program; lower panels, CBB staining.

**Supplementary Table 1. Loss-of-function of *PDK1.1* and *PDK1.2* leads to defective cotyledon development.** Seven-day-old seedlings were observed and numbers of seedlings with defective cotyledon development were counted.

| Genotype | Total | Seedlings with normal cotyledon | Seedlings with fused cotyledon | |
|---|---|---|---|---|
| | | | Numbers | Percentage |
| Col-0 | 1595 | 1595 | 0 | 0% |
| *pdk1.1-1* | 1404 | 1401 | 3 | 0.21% |
| *pdk1.2-1* | 1333 | 1431 | 2 | 0.15% |
| *pdk1.1-1 pdk1.2-1* | 1424 | 1167 | 257 | 18.05% |

**Supplementary Table 2. List of plant lines, including mutants and maker lines, used in this study.**

| *Arabidopsis* lines | Source | Identifier |
|---|---|---|
| Col-0 | N/A | N/A |
| *pdk1.1-4* (short as *pdk1.1*) | This study | SALK_113251 |
| *pdk1.2-2* (short as *pdk1.2*) | This study | SALK_017433 |
| *pdk1.1-4 pdk1.2-2* (*pdk1.1 pdk1.2*) | This study | N/A |
| *pdk1.1 pdk1.2 d0 d1 d2* | This study | N/A |
| *pinoid-14* (*pid-14*) | Huang et al., 2010 | SALK_049736 |
| *rcn1-6* | Dai et al., 2013 | SALK_059903 |
| *pdk1.1 pdk1.2 rcn1-6* | This study | N/A |
| *pdk1.1 pdk1.2 pid-14* | This study | N/A |
| *d6pk-1* | Zourelidou et al., 2009 | SALK_061847 |
| *d6pkl1-1* | Zourelidou et al., 2009 | SALK_056618 |
| *d6pkl2-2* | Zourelidou et al., 2009 | SALK_086127 |
| *d6pkl3-2* | Zourelidou et al., 2009 | SALK_047347 |
| *d6pk-1 d6pkl1-1 d6pkl2-2* (*d0d1d2*) | Zourelidou et al., 2009 | N/A |
| *d6pk-1 d6pkl1-1 d6pkl2-2 d6pkl3-2* (*d0d1d2d3*) | Zourelidou et al., 2009 | N/A |
| *pPIN1::PIN1-YFP* | Ref. 55 | N/A |
| *pPIN1::PIN1-GFP* | Benková et al., 2003 | N/A |
| *pPIN2::PIN2-GFP* | Xu and Scheres, 2005 | N/A |
| *pPIN3::PIN3-GFP* | Zadnikova et al., 2010 | N/A |
| *DR5rev::GFP* | Friml et al., 2004 | N/A |
| *p35S::YFP-D6PK* | Zourelidou et al., 2009 | N/A |
| *pD6PK::YFP-D6PK* | Willige et al., 2013 | N/A |
| *pPDK1.1::PDK1.1* in *pdk1.1 pdk1.2* | This study | Lines #1, 3, 4 |
| *DR5rev::GFP pdk1.1 pdk1.2* | This study | N/A |
| *pPIN1::PIN1-YFP pdk1.1 pdk1.2* | This study | N/A |
| *pPIN2::PIN2-GFP pdk1.1 pdk1.2* | This study | N/A |
| *pPIN3::PIN3-GFP pdk1.1 pdk1.2* | This study | N/A |

| | | |
|---|---|---|
| *p35S::YFP-D6PK pdk1.1 pdk1.2* | This study | N/A |
| *p35S::Venus-D6PK* in Col-0 | This study | N/A |
| *p35S::Venus-D6PK*$^{S345A}$ in Col-0 | This study | N/A |
| *p35S::Venus-D6PK*$^{S345D}$ in Col-0 | This study | N/A |
| *p35S::Venus-D6PK* in *pdk1.1 pdk1.2* | This study | N/A |
| *p35S::Venus-D6PK*$^{S345A}$ in *pdk1.1 pdk1.2* | This study | N/A |
| *p35S::Venus-D6PK*$^{S345D}$ in *pdk1.1 pdk1.2* | This study | N/A |
| *p35S::mCherry-D6PK* in Col-0 | This study | N/A |
| *p35S::mCherry-D6PKL1* in Col-0 | This study | N/A |
| *p35S::mCherry-D6PKL2* in Col-0 | This study | N/A |
| *p35S::mCherry-D6PKL3* in Col-0 | This study | N/A |
| *p35S::Venus-PDK1.1* in Col-0 | This study | N/A |
| *p35S::Venus-PDK1.2* in Col-0 | This study | N/A |
| *p35S::Venus-PDK1.1* in *pdk1.1 pdk1.2* | This study | N/A |
| *p35S::Venus-PDK1.2* in *pdk1.1 pdk1.2* | This study | N/A |
| *p35S::mCherry-PDK1.1* in Col-0 | This study | N/A |
| *p35S::mCherry-PDK1.2* in Col-0 | This study | N/A |
| *p35S::mCherry-PDK1.1* in *pdk1.1 pdk1.2* | This study | N/A |
| *p35S::mCherry-PDK1.2* in *pdk1.1 pdk1.2* | This study | N/A |
| *pRPS5A::mCherry-PDK1.1* in Col-0 | This study | N/A |
| *pRPS5A::mCherry-PDK1.2* in Col-0 | This study | N/A |
| *pRPS5A::mCherry-PDK1.1* in *pdk1.1 pdk1.2* | This study | N/A |
| *pRPS5A::mCherry-PDK1.2* in *pdk1.1 pdk1.2* | This study | N/A |
| *pPDK1.1::Venus-PDK1.1* in Col-0 | This study | N/A |
| *pPDK1.1::Venus-PDK1.1* in *pdk1.1 pdk1.2* | This study | N/A |
| *p35S::YFP-D6PK* × *pRPS5A::mCherry-PDK1.1* | This study | N/A |
| *p35S::YFP-D6PK* × *pRPS5A::mCherry-PDK1.2* | This study | N/A |
| *p35S:: Venus-PDK1.1* × *p35S::mCherry-D6PKL2* | This study | N/A |
| *p35S:: Venus-PDK1.1* × *p35S::mCherry-D6PKL3* | This study | N/A |

**Supplementary Table 3. List of reagents used in this study.**

| Reagent or Resource | Source | Identifier |
|---|---|---|
| Murashige & Skoog Basal Medium including vitamins | Duchefa | Cat. # M0222.0050 |
| Hygromycin | Duchefa | Cat. # H0192.0001 |
| Rifampicin | Duchefa | Cat. # R0146.0025 |
| Kanamycin sulphate monohydrate | Duchefa | Cat. # K0126.0025 |
| Spectinomycin | Duchefa | Cat. # S0188.0025 |
| Ampicillin Sodium | Duchefa | Cat. # 69-52-3 |
| Plant Agar | Duchefa | Cat. # P1001.1000 |
| Naptalam [N-(1-Naphthyl)phthalamidic acid ] | Duchefa | Cat. # N0926.0250 |
| Indole 3-acetic acid (IAA) | Sigma | Cat. # I2886 |
| [$^3$H]-IAA | American Radiolabeled Chemicals | Cat. # ART-0340 |
| imidazole | Sigma | Cat. # I5513 |
| Difco Skim milk | BD (Becton Dickinson) | Cat. # 232100 |
| Glufosinate-ammonium | Sigma | Cat. # 45520 |
| U-73122 hydrate | Sigma | Cat. # U6756 |
| 1-butanol | Sigma | Cat. # 281549 |
| Wortmannin | Sigma | Cat. # W1628 |
| Phenylarsine oxide (PAO) | Sigma | Cat. # P3075 |
| LY-294,002 hydrochloride | Sigma | Cat. # L9908 |
| brefeldin A | Sigma | Cat. # B7651 |
| Superdex 200 Increase 10/300 GL | GE Healthcare | Cat. # 28990944 |
| PIP Array | Echelon Biosciences | Cat. # P-6100 |
| PIP Strips | Echelon Biosciences | Cat. # P-6001 |
| Propidium Iodide - 1.0 mg/mL | Thermo Fisher Scientific | Cat. # P3566 |
| cOmplete protease inhibitor cocktail | Sigma/Roche | Cat. # 4693124001 |
| PhosSTOP™ | Sigma/Roche | Cat. # 4906837001 |

| | | |
|---|---|---|
| BL21(DE3) Competent E. coli | NEB BioLabs | Cat. # C2527H |
| One Shot Mach1 T1 Competent E. coli | Life Tech OCI | Cat. # C862003 |
| **Recombinant Proteins** | | |
| GST-PDK1.1 | This study | N/A |
| GST-PDK1.2 | This study | N/A |
| His-PDK1.1 | This study | N/A |
| His-PDK1.2 | This study | N/A |
| His-PDK1.1N | This study | N/A |
| His-PDK1.2N | This study | N/A |
| His-PDK1.1C | This study | N/A |
| His-PDK1.2C | This study | N/A |
| His-D6PK | This study | N/A |
| His-D6PK$^{S345A}$ | This study | N/A |
| His-D6PK$^{S345D}$ | This study | N/A |
| GST-D6PKL1 | This study | N/A |
| His-D6PKL2 | This study | N/A |
| His-PIN1HL | This study | N/A |
| His-PIN2HL | This study | N/A |
| His-PIN2-3HL | This study | N/A |
| Bovine Serum Albumin | Sigma | Cat. # A2153 |
| **Antibodies** | | |
| Rabbit anti-PIN1 | Ref. 62 | |
| Rabbit anti-PIN2 | Ref. 63 | |
| Mouse anti-His-tag Antibody, HRP conjugated | Agrisera | Cat. # AS15 2930 |
| Rat monoclonal Anti-RFP [Clone 5F8] | Chromotek | Cat. # 5F8-100 |
| Anti-GFP Antibody, HRP Conjugated | MACS Molecular | Cat. # 130-091-833 |
| Anti-rat IgG Antibody, HRP Conjugated | Sigma | Cat. # AP202P |
| **Kits** | | |
| Bio-Safe™ Coomassie Stain #1610786 | Bio-Rad | Cat. # 1610786 |

| | | |
|---|---|---|
| GeneJET Plasmid Miniprep Kit | Thermo Fisher Scientific | Cat. # K0503 |
| GeneJET Gel extraction kit | Thermo Fisher Scientific | Cat. # K0692 |
| Pierce® Glutathione Agarose | Thermo Fisher Scientific | Cat. # 16101 |
| HisPur™ Ni-NTA Resin | Thermo Fisher Scientific | Cat. # 88222 |
| **Restriction enzymes and Gateway Clonases** | | |
| FastDigest SalI | Thermo Fisher Scientific | Cat. # FD0644 |
| FastDigest SacI | Thermo Fisher Scientific | Cat. # FD1133 |
| FastDigest XbaI | Thermo Fisher Scientific | Cat. # FD0684 |
| FastDigest EcoRI | Thermo Fisher Scientific | Cat. # FD0274 |
| FastDigest XhoI | Thermo Fisher Scientific | Cat. # FD0694 |
| FastDigest BamHI | Thermo Fisher Scientific | Cat. # FD0054 |
| T4 DNA Ligase Buffer | Thermo Fisher Scientific | Cat. # 46300-018 |
| T4 DNA Ligase (1 U/µL) | Thermo Fisher Scientific | Cat. # 15224-017 |
| Gateway™ LR Clonase™ II Enzyme mix | Thermo Fisher Scientific | Cat. # 11791100 |
| Gateway BP Clonase II Enzyme mix | Thermo Fisher Scientific | Cat. # 11789100 |
| **Software and Algorithms** | | |
| DNAMAN | DNAMAN | https://www.lynnon.com/ |
| image J | image J | https://imagej.net/ |
| Fiji | Fiji | https://fiji.sc/ |

**Supplementary Table 4. List of primers used in this study.**

| **Primers** | **Oligonucleotide (5' to 3')** | **Purpose** |
|---|---|---|
| **For Genotyping T-DNA Insertional Mutants** | | |
| **Oligo Name** | **Sequence** | **Allele** |
| pdk1.1-LP | TTTCTTTGGTACTCTGTGGAACACC | *pdk1.1* |
| pdk1.1-RP | TTACCACGGTTTTGTGAAAGG | |
| pdk1.2-LP | GTTTGCGCAAGTTGATGAGTCGTAC | *pdk1.2* |
| pdk1.2-RP | ACCAACAAACCAAGACTGATATAC | |
| d6pk-LP | TGAGAATCATCAACTGTGGAAAC | *d6pk-1* |
| d6pk-RP | TTTGGTGATGGAGTTTTGTCC | |
| d6pkl1-LP | TCTCTTCATTTCCATGGAAGG | *d6pkl1-1* |
| d6pkl1-RP | CTCCAGTTTTACTGTCGCTGC | |
| d6pkl2-LP | CTTCGCCTTTGATGATCTCTG | *d6pkl2-2* |
| d6pkl2-RP | AGTGACGAGAGTAGCTGCAGC | |
| d6pkl3-LP | CCATTAAACGACGAAACATCG | *d6pkl3-2* |
| d6pkl3-RP | TAACAAGCTTCTTCCTCGCTG | |
| pid-14_LP | CAGTCGGGAAACTCAACTGTC | *pid-14* |
| pid-14_RP | ATTTTGCGATGAAAGTTGTGG | |
| LB1 | GCCTTTTCAGAAATGGATAAATAGCCTTGCTTCC | SALL lines |
| LBb1.3 | ATTTTGCCGATTTCGGAAC | SALK lines |
| **For qRT-PCR** | | |
| **Oligo Name** | **Sequence** | **Gene** |
| PDK1.1-F | GATCTATGTCGACCCGTCAAAAC | |
| PDK1.1-R | AGCGGTTCTGAAGAGTCTCGATTGCC | |
| PDK1.2-F | CTAATGATATCAGCAGTGAAGAAGC | |
| PDK1.2-R | CGATTGCCTTTTTCCACTGCAAAGC | |
| ACTIN7-F | CCGGTATTGTGCTCGATTCTG | |
| ACTIN7-R | TTCCCGTTCTGCGGTAGTGG | |
| **Constructs for Protein expression** | | |
| **Oligo Name** | **Sequence** | **Plasmid** |
| PDK1.1-1_BamHI | CGGGATCCATGTTGGCAATGGAGAAAGAA | pET28a-PDK1.1 |
| PDK1.1-2_XhoI | CCGCTCGAGGCGGTTCTGAAGAGTCTCGA | pET28a-PDK1.1 |
| PDK1.1C-10_XhoI | CCGCTCGAGCGATTCTCCTGGCTCTAAAAAC | pET28a-PDK1.1N |
| PDK1.1N-9_BamHI | CGGGATCCGTTCTGATGATATCAGCGGTGAAG | pET28a-PDK1.1C |
| PDK1.2-1_BamHI | CGGGATCCATGTTGACAATGGACAAGGAATTTG | pET28a-PDK1.2 |
| PDK1.2-2_XhoI | CCGCTCGAGACGGTTTTGAAGAGTTTCG | pET28a-PDK1.2 |

| | | |
|---|---|---|
| PDL1C-10_XhoI | CCGCTCGAGCGATTCTCCCGGTTCTAGAAAC | pET28a-PDK1.2N |
| PDL1N-9_BamHI | CGGGATCCGTTCTAATGATATCAGCAGTGAAGAAGC | pET28a-PDK1.2C |
| D6PK-1_EcoRI | GGAATTCATGATGGCTTCAAAAACTCCAG | pET28a-D6PK |
| D6PK-2_XhoI | CCGCTCGAGTCAGAAGAAATCAAACTCAAGATA | pET28a-D6PK |
| D6PKL1-1_EcoRI | GGAATTCATGGCCTCGAAGTATGGTTCTGG | pET28a-D6PKL1 |
| D6PKL1-2_XhoI | CCGCTCGAGTCAAAAGAAATCGAACTCCAG | pET28a-D6PKL1 |
| D6PKL2-1_BamHI | CGGGATCCATGGCGTCCACTCGTAAACCCAG | pET28a-D6PKL2 |
| D6PKL2-2_XhoI | CCGCTCGAGCTAGAAGAAATCAAATTCCAAATAG | pET28a-D6PKL2 |
| D6PK-3_S345A | CACAGGCGCCCGTTCTATGGCTTTTGTTGGCAC | S345A point mutation |
| D6PK-4_S345A | AGCCATAGAACGGGCGCCTGTGGGCTCTGCAAC | S345A point mutation |
| D6PK-5_S345D | CACAGGCGCCCGTTCTATGGATTTTGTTGGCAC | S345D point mutation |
| D6PK-6_S345D | ATCCATAGAACGGGCGCCTGTGGGCTCTGCAAC | S345D point mutation |

**For Gateway Cloning**

| Oligo Name | Sequence | Plasmid |
|---|---|---|
| PDK1.2-3-GW_F | GGGGACAAGTTTGTACAAAAAAGCAGGCTTCATGTTGACAATGGACAAGGAATTTG | pDONR221-PDK1.2 |
| PDK1.2-4-GW_R | GGGGACCACTTTGTACAAGAAAGCTGGGTTACGGTTTTGAAGAGTTTCG | pDONR221-PDK1.2 |
| PDK1.2-6-GW_R_stop | GGGGACCACTTTGTACAAGAAAGCTGGGTTTCAACGGTTTTGAAGAGTTTCG | pDONR221-PDK1.2 stop |
| PDK1.1-8-GW_R | GGGGACCACTTTGTACAAGAAAGCTGGGTTCGATTCTCCTGGCTCTAAAAAC | pDONR221-PDK1.2N |
| PDK1.1-7-GW_F | GGGGACAAGTTTGTACAAAAAAGCAGGCTTCATGGTTCTGATGATATCAGCGGTGAAG | pDONR221-PDK1.1C |
| PDK1.1-3-GW_F | GGGGACAAGTTTGTACAAAAAAGCAGGCTTCATGTTGGCAATGGAGAAAGAA | pDONR221-PDK1.1 |
| PDK1.1-4-GW_R | GGGGACCACTTTGTACAAGAAAGCTGGGTTGCGGTTCTGAAGAGTCTCGA | pDONR221-PDK1.1 |
| PDK1.1-6-GW_R_stop | GGGGACCACTTTGTACAAGAAAGCTGGGTTTCAGCGGTTCTGAAGAGTCTCGA | pDONR221-PDK1.1stop |
| PDK1.2-8-GW_R | GGGGACCACTTTGTACAAGAAAGCTGGGTTCGATTCTCCCGGTTCTAGAAAC | pDONR221-PDK1.1N |
| PDK1.2-7-GW_F | GGGGACAAGTTTGTACAAAAAAGCAGGCTTCATGGTTCTAATGATATCAGCAGTG | pDONR221-PDK1.2C |
| D6PK-9-GW-F | GGGGACAAGTTTGTACAAAAAAGCAGGCTTCATGATGGCTTCAAAAACTCCAG | pDONR221-D6PK |
| D6PK-10-GW-R | GGGGACCACTTTGTACAAGAAAGCTGGGTTGAAGAA | pDONR221-D6PK |

| | | |
|---|---|---|
| | ATCAAACTCAAGATA | |
| D6PK-12-GW-R | GGGGACCACTTTGTACAAGAAAGCTGGGTTTCAGAAGAAATCAAACTCAAGATA | pDONR221-D6PKstop |
| D6PKL1-9-GW-F | GGGGACAAGTTTGTACAAAAAAGCAGGCTTCATGGCCTCGAAGTATGGTTCTGG | pDONR221-D6PKL1 |
| D6PKL1-10-GW-R | GGGGACCACTTTGTACAAGAAAGCTGGGTTAAAGAAATCGAACTCCAGATAATTAC | pDONR221-D6PKL1 |
| D6PKL1-12-GW-Rstop | GGGGACCACTTTGTACAAGAAAGCTGGGTTTCAAAAGAAATCGAACTCCAGATAATTAC | pDONR221-D6PKL1stop |
| D6PKL2-9-GW-F | GGGGACAAGTTTGTACAAAAAAGCAGGCTTCATGGCGTCCACTCGTAAACCCAG | pDONR221-D6PKL2 |
| D6PKL2-10-GW-R | GGGGACCACTTTGTACAAGAAAGCTGGGTTGAAGAAATCAAATTCCAAATAG | pDONR221-D6PKL2 |
| D6PKL2-12-GW-Rstop | GGGGACCACTTTGTACAAGAAAGCTGGGTTCTAGAAGAAATCAAATTCCAAATAG | pDONR221-D6PKL2stop |
| D6PKL3-9-GW-F | GGGGACAAGTTTGTACAAAAAAGCAGGCTTCATGGATTCTTCTTCATCAGTCG | pDONR221-D6PKL3 |
| D6PKL3-10-GW-R | GGGGACCACTTTGTACAAGAAAGCTGGGTTGAAGAAATCAATTTCCAAATAATTACC | pDONR221-D6PKL3 |
| D6PKL3-12-GW-Rstop | GGGGACCACTTTGTACAAGAAAGCTGGGTTTCAGAAGAAATCAATTTCCAAATAATTACC | pDONR221-D6PKL3stop |
| pPDK1.1-attB4 | GGGGACAACTTTGTATAGAAAAGTTGTAATCAGTAGAGTTACTTGTAGGGCAAGTAAG | pDONR-P4P1r-pPDK1.1 |
| pPDK1.1-attB1r | GGGGACTGCTTTTTTGTACAAACTTGATTCCCTTTTCTTCTAAATCTCGATGCGACCC | pDONR-P4P1r-pPDK1.1 |
| pPDK1.2-attB4 | GGGGACAACTTTGTATAGAAAAGTTGTAATGGGTTTTCCACCTTGAGGCTTGAGCG | pDONR-P4P1r-pPDK1.2 |
| pPDK1.2-attB1r | GGGGACTGCTTTTTTGTACAAACTTGACTCCAAACTTGTTCTACATTTCGACGAGACCC | pDONR-P4P1r-pPDK1.2 |
| Venus-GW_F | GGGGACAAGTTTGTACAAAAAAGCAGGCTTCATGGTGAGCAAGGGCGAGGAGCTGTTC | pDONR221-Venus-PDK1.1/2stop |
| mCherry-GW_F | GGGGACAAGTTTGTACAAAAAAGCAGGCTTCATGGTGAGCAAGGGCGAGGAGGACAAC | pDONR221-mCherry-PDK1.1/2stop |

**Supplementary Table 5. List of plasmids used in this study.**

| **Plasmid Name** | **Purpose** | **Information** |
|---|---|---|
| **Recombinant Protein Expression in *E.coli*** | | |
| **Plasmid Name** | **Recombinant Protein** | **Resistance in *E.coli*** |
| pGEX-4T-1-PDK1.1 | GST-PDK1.1 | Ampicillin |
| pGEX-4T-1-PDK1.2 | GST-PDK1.2 | Ampicillin |
| pET28a-PDK1.1 | His-PDK1.1 | Kanamycin |
| pET28a-PDK1.2 | His-PDK1.2 | Kanamycin |
| pET28a-PDK1.1N | His-PDK1.1N | Kanamycin |
| pET28a-PDK1.2N | His-PDK1.2N | Kanamycin |
| pET28a-PDK1.1C | His-PDK1.1C | Kanamycin |
| pET28a-PDK1.2C | His-PDK1.2C | Kanamycin |
| pET28a-D6PK | His-D6PK | Kanamycin |
| pET28a-D6PK$^{S345A}$ | His-D6PK$^{S345A}$ | Kanamycin |
| pET28a-D6PK$^{S345D}$ | His-D6PK$^{S345D}$ | Kanamycin |
| pGEX-4T-1-D6PKL1 | GST-D6PKL1 | Ampicillin |
| pET28a-D6PKL2 | His-D6PKL2 | Kanamycin |
| pGEX-4T-1 | GST | Ampicillin |
| pET28a-PIN1HL | His-PIN1HL | Kanamycin |
| pET28a-PIN2HL | His-PIN2HL | Kanamycin |
| pET28a-PIN3HL | His-PIN3HL | Kanamycin |
| **Gateway Entry Vectors** | | |
| pDONR221-PDK1.1-with stop | Entry Vector | Kanamycin |
| pDONR221-PDK1.1-no stop | Entry Vector | Kanamycin |
| pDONR221-PDK1.2-with stop | Entry Vector | Kanamycin |
| pDONR221-PDK1.2-no stop | Entry Vector | Kanamycin |
| pDONR221-PDK1.1N-with stop | Entry Vector | Kanamycin |
| pDONR221-PDK1.1C-with stop | Entry Vector | Kanamycin |
| pDONR221-PDK1.2N-with stop | Entry Vector | Kanamycin |
| pDONR221-PDK1.2C-with stop | Entry Vector | Kanamycin |
| pDONR221-mCherry-PDK1.1-with stop | Entry Vector | Kanamycin |
| pDONR221-mCherry-PDK1.2-with stop | Entry Vector | Kanamycin |
| pDONR221-Venus-PDK1.1-with stop | Entry Vector | Kanamycin |
| pDONR221-Venus-PDK1.2-with stop | Entry Vector | Kanamycin |

| | | |
|---|---|---|
| pDONR221-D6PK-no stop | Entry Vector | Kanamycin |
| pDONR221-D6PK-with stop | Entry Vector | Kanamycin |
| pDONR221-D6PKL1-no stop | Entry Vector | Kanamycin |
| pDONR221-D6PKL1-with stop | Entry Vector | Kanamycin |
| pDONR221-D6PKL2-no stop | Entry Vector | Kanamycin |
| pDONR221-D6PKL2-with stop | Entry Vector | Kanamycin |
| pDONR221-D6PKL3-no stop | Entry Vector | Kanamycin |
| pDONR221-D6PKL3-with stop | Entry Vector | Kanamycin |
| pDONR-P4-P1r-Venus | Entry Vector | Kanamycin |
| pDONR-P4-P1r-mCherry | Entry Vector | Kanamycin |
| pDONR-P4-P1r-pPDK1.1 | Entry Vector | Kanamycin |
| pDONR-P4-P1r-pPDK1.2 | Entry Vector | Kanamycin |

**Binary Vectors**

| **Plasmid Name** | **Transgenic Lines** | **Resistance *in planta*** |
|---|---|---|
| pB7m24GW2 | *p35S::Venus-D6PK* | Basta |
| pB7m24GW2 | *p35S::Venus-D6PK*$^{S345A}$ | Basta |
| pB7m24GW2 | *p35S::Venus-D6PK*$^{S345D}$ | Basta |
| pB7m24GW2 | *p35S::mCherry-D6PK* | Basta |
| pB7m24GW2 | *p35S::mCherry-D6PKL1* | Basta |
| pB7m24GW2 | *p35S::mCherry-D6PKL2* | Basta |
| pB7m24GW2 | *p35S::mCherry-D6PKL3* | Basta |
| pB7m24GW2 | *p35S::Venus-PDK1.1* | Basta |
| pB7m24GW2 | *p35S::Venus-PDK1.2* | Basta |
| pB7m24GW2 | *p35S::mCherry-PDK1.1* | Basta |
| pB7m24GW2 | *p35S::mCherry-PDK1.2* | Basta |
| pB7m24GW3 | *pPDK1.1::Venus-PDK1.1* | Basta |
| pB7m24GW3 | *pPDK1.2::Venus-PDK1.2* | Basta |
| pB7m24GW3 | *pRPS5A::Venus-PDK1.1* | Basta |
| pB7m24GW3 | *pRPS5A::Venus-PDK1.2* | Basta |
| pB7m24GW3 | *pRPS5A::mCherry-PDK1.1* | Basta |
| pB7m24GW3 | *pRPS5A::mCherry-PDK1.2* | Basta |
| pCambia1300+pBI121 | *pPDK1.1::GUS* | Hygromycin |
| pCambia1300+pBI121 | *pPDK1.1::GUS* | Hygromycin |
| pCambia1300 | *pPDK1.1::PDK1.1* | Hygromycin |
